# Mapping enhancer-gene regulatory interactions from single-cell data

**DOI:** 10.1101/2024.11.23.624931

**Authors:** Maya U. Sheth, Wei-Lin Qiu, X. Rosa Ma, Andreas R. Gschwind, Evelyn Jagoda, Anthony S. Tan, Hjörleifur Einarsson, Bram L. Gorissen, Danilo Dubocanin, Christopher S. McGinnis, Dulguun Amgalan, Ansuman T. Satpathy, Thouis R. Jones, Lars M. Steinmetz, Anshul Kundaje, Berk Ustun, Jesse M. Engreitz, Robin Andersson

**Affiliations:** The Novo Nordisk Foundation Center for Genomic Mechanisms of Disease, Broad Institute of MIT and Harvard, Cambridge, Massachusetts, USA; Department of Genetics, Stanford University School of Medicine, Stanford, California, USA; Basic Sciences and Engineering Initiative, Betty Irene Moore Children’s Heart Center, Lucile Packard Children’s Hospital, Stanford, California, USA; Department of Bioengineering, Stanford University, Stanford, California, USA; Section for Computational and RNA Biology, Department of Biology, University of Copenhagen, Copenhagen, Denmark; Analytic and Translational Genetics Unit, Massachusetts General Hospital, Boston, Massachusetts 02114, USA; Department of Pathology, Stanford University, Stanford, CA, USA; Parker Institute for Cancer Immunotherapy, San Francisco, California, USA; Genome Biology Unit, European Molecular Biology Laboratory (EMBL), Heidelberg, Germany; Stanford Genome Technology Center, Stanford University School of Medicine, Palo Alto, California, USA; Department of Computer Science, Stanford University, Stanford, California, USA; Halıcıoğlu Data Science Institute and Department of Computer Science & Engineering, University of California San Diego, San Diego, California, USA; Gene Regulation Observatory, Broad Institute of MIT and Harvard, Cambridge, Massachusetts, USA; Stanford Cardiovascular Institute, Stanford University School of Medicine, Stanford, California, USA

## Abstract

Mapping enhancers and their target genes in specific cell types is crucial for understanding gene regulation and human disease genetics. However, accurately predicting enhancer-gene regulatory interactions from single-cell datasets has been challenging. Here, we introduce a new family of classification models, scE2G, to predict enhancer-gene regulation. These models use features from single-cell ATAC-seq or multiomic RNA and ATAC-seq data and are trained on a CRISPR perturbation dataset including >10,000 evaluated element-gene pairs. We benchmark scE2G models against CRISPR perturbations, fine-mapped eQTLs, and GWAS variant-gene associations and demonstrate state-of-the-art performance at prediction tasks across multiple cell types and categories of perturbations. We apply scE2G to build maps of enhancer-gene regulatory interactions in heterogeneous tissues and interpret noncoding variants associated with complex traits, nominating regulatory interactions linking *INPP4B* and *IL15* to lymphocyte counts. The scE2G models will enable accurate mapping of enhancer-gene regulatory interactions across thousands of diverse human cell types.

## Introduction

The human genome encodes millions of regulatory elements called enhancers that tune gene expression in specific cell types^1–6^. Identifying which enhancers regulate which genes in which cell types (“enhancer-gene regulatory interactions”) is critical for understanding gene regulation and the functions of disease-related genetic variants^7–10^. Yet, we lack maps of enhancer-gene interactions for most cell types in the human body.

Previously, we and others have built computational models that predict enhancer-gene regulatory interactions from bulk measurements of chromatin state and 3D contacts^11–20^, including for example the Activity-by-Contact (ABC)^21^ and ENCODE-rE2G models^20^. These models have enabled creating genome-wide enhancer-gene maps in hundreds of human cell types and tissues^20,21^. However, these maps are limited in their applicability or resolution to study many individual cell types and cell states in complex tissues due to the heterogeneity of cell types within a tissue and challenges in purifying individual cell types for bulk epigenomic assays.

Advances in single-cell profiling of chromatin accessibility and RNA expression now provide a strategy to access virtually any cell type and state across heterogeneous tissues in the human body^22–27^. In theory, predictive models that use single-cell data as inputs could be applied to characterize enhancer-gene regulation in many more cell types and states than possible with bulk models. To this end, various approaches have been explored, mostly based on using unsupervised approaches such as correlating the accessibility of a distal element with the expression of a nearby gene^28–37^. However, it has been challenging to assess the accuracy of these models due to the lack of robust benchmarking datasets and pipelines, and their consistency for building large atlases of enhancer-gene regulatory interactions has not been characterized. Recent work by the ENCODE Consortium to collect large-scale gold-standard datasets of enhancer perturbations now provides an opportunity to systematically evaluate these models, and to develop new models based on supervised learning^20,21,38,39^.

Here we introduce a new family of single-cell enhancer-to-gene (scE2G) models that utilize scATAC or multiomic scATAC and scRNA data to make genome-wide predictions of regulatory enhancer-gene interactions. We systematically benchmark predictions from scE2G models as well as 10 other published models using and extending pipelines developed with the ENCODE Consortium^20^. The scE2G models demonstrate state-of-the-art performance across a range of prediction tasks and diverse single-cell datasets involving CRISPR perturbations, fine-mapped eQTLs, and GWAS variant-gene associations. Important features in the scE2G models include (i) a pseudobulk ABC score, which integrates enhancer and promoter chromatin accessibility, estimates of genomic distance and 3D contact, (ii) promoter class based on expression across cell types, and (iii) the Kendall correlation between chromatin accessibility and gene expression across single cells within a cell type, which appears to detect stochastic transcriptional bursting. We apply scE2G to build maps of enhancer-gene regulatory interactions in 42 cell types and states from complex tissues, and demonstrate the utility of these maps in interpreting risk variants for common diseases. scE2G models are robust to sequencing depth, cell number, and assay type. In combination with expanding efforts to collect single-cell data, scE2G will facilitate future efforts to build genome-wide enhancer maps in many thousands of cell types in the human body.

### Predicting regulatory enhancer-gene interactions from single-cell data

We develop two models to predict enhancer-gene regulatory interactions from single-cell data: scE2G*^ATAC^*, which uses scATAC-seq data, and scE2G*^Multiome^*, which uses as input paired scRNA and scATAC-seq data. Both models are classifiers based on logistic regression that take single-cell data as input and predict which chromatin accessible elements act as enhancers to regulate which genes in a given cell population (**Fig. 1a**). We trained scE2G models using gold-standard CRISPR perturbation data in K562 cells, allowing subsequent application of the trained models to single-cell data from other cell types (**Fig. 1a**). As such, the scE2G models make cell-type-specific predictions of enhancer-gene regulatory interactions by analyzing data from one cell type or state at a time (*e.g.*, one cluster of cells from a single-cell dataset).

**Figure 1.**
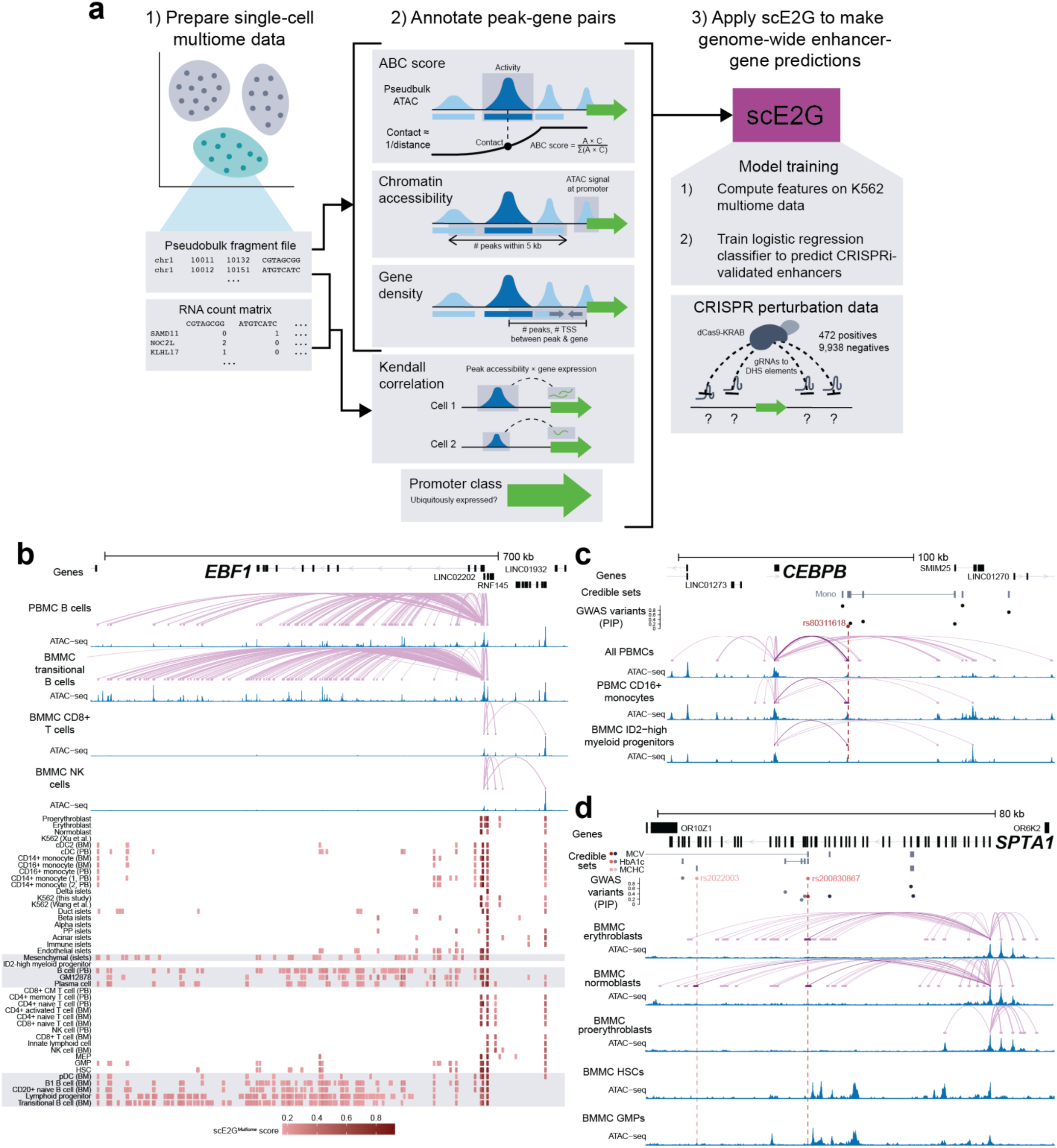
scE2G identifies enhancer-gene regulatory interactions from single-cell data across 42 cell types. **a.** Conceptual diagram of scE2G. **b.** scE2G*^Multiome^* predictions for *EBF1* across 44 cell types for the region chr1:158,605,000-158,702,000 (hg38). Enhancer-gene linking arcs and normalized ATAC signals are shown for PBMC B cells, BMMC transitional B cells, BMMC CD8+ T cells, and BMMC NK cells. The heatmap shows enhancer predictions with a score above the threshold for *EBF1* in each cell type, ordered by similarity in RNA expression profile. The color of the enhancer indicates the magnitude of the scE2G score. Highlighted rows in the heatmap indicate B cell types expected to have more active regulatory landscapes for *EBF1*. “PB” indicates cell type is from PBMCs; “BM” indicates cell type is from BMMCs. **c.** scE2G*^Multiome^* predictions and normalized ATAC signals for *CEBPB* in combined PBMCs, PBMC CD16+ monocytes, BMMC ID2-high myeloid progenitors, and PBMC natural killer cells for the region chr20:50,145,000-50,310,000 (hg38). Noncoding fine-mapped GWAS variants with PIP > 10% and credible sets for monocyte count are shown. The variant rs80311618, which overlaps a myeloid-specific enhancer, is indicated in red and predictions overlapping this variant are shown in darker purple. The width of prediction arcs is proportional to scE2G score. **d.** scE2G*^Multiome^* predictions and normalized ATAC signals for *SPTA1* in five BMMC cell types: erythroblasts, normoblasts, proerythroblasts, hematopoietic stem cells (HSCs), and granulocyte-macrophage progenitors (GMPs) for the region chr1:158,605,000-158,702,000 (hg38). Noncoding fine-mapped GWAS variants with PIP > 10% and credible sets for hemoglobin A1C (HbA1c), mean corpuscular volume (MCV), and mean corpuscular hemoglobin concentration (MCHC) are shown. The variants rs2022003, in a credible set for MCHC, and rs200830867, in credible sets for HbA1c and MCV, are highlighted in red. Predictions overlapping these variants are shown in darker purple. The width of prediction arcs is proportional to scE2G score.

The scE2G*^ATAC^* model uses six features derived from single-cell data: (i) an ABC score, with activity computed from pseudobulk scATAC-seq data and contact estimated as an inverse function of genomic distance; (ii) two features describing chromatin accessibility at or near the enhancer or promoter; (iii) two features related to genomic distance and gene density; and (iv) the class of the promoter, based on whether the gene is ubiquitously expressed across many cell types. Apart from the ABC score being adapted to single cell data, this first set of features is similar to those used in the ENCODE-rE2G model we developed for bulk data^20^. In addition, for the scE2G*^Multiome^* model, we compute the Kendall correlation between element accessibility and gene expression across single cells, then integrate it with the ABC score as a new feature termed the Activity, Responsiveness, and Contact (ARC)-E2G score to improve calibration across different sequencing depths (see **Methods**, **Fig. S1**).

We trained scE2G models on a dataset of 10,410 genomic element-gene pairs (candidate enhancer-gene pairs) tested with CRISPR in K562 erythroleukemia cells. This dataset contains examples that we previously aggregated from 3 studies from ENCODE and others^20^. It includes 472 unique “positive” element-gene pairs where CRISPR perturbation of the element led to a significant decrease in gene expression and 9,938 “negative” element-gene pairs where no significant reduction in expression (at least 15 to 25% effect size) was observed^20^. We constructed features from multiomic single-cell RNA and ATAC-seq (10x Multiome) data in K562 cells^40^, and trained logistic regression classifiers to distinguish positives from negatives and evaluated their performance using hold-one-chromosome-out cross-validation. For feature selection, we used a best-subsets approach to identify the minimal set of features, described above, that can achieve near-optimal performance at classifying the element-gene pairs in the CRISPR dataset (see **Methods**).

Having learnt how to accurately classify CRISPR elements in K562 cells from features derived from single-cell data, the trained scE2G models can be applied to predict interactions from new data from new cell types, unseen in the training process, to infer all enhancer-gene regulatory interactions across the genome. Each element-gene pair is annotated with a score corresponding to the probability of a regulatory effect from the logistic regression model. Because scE2G is designed to detect elements that act as enhancers, as opposed to silencers, topological elements, or other types of regulatory elements, we name elements with scores above a chosen threshold as scE2G “enhancers”.

To demonstrate the utility of scE2G, we applied scE2G*^Multiome^* to build maps of enhancer-gene regulatory interactions in 42 different cell types (defined here as a cell cluster annotated based on expression of marker genes) including those from human peripheral blood mononuclear cells (PBMCs), bone marrow mononuclear cells (BMMCs), and pancreatic islets (**Fig. 1b, Fig. S2**). For each cell type, we (i) pseudobulk the scATAC data to compute the ABC score and other features; (ii) use the single-cell RNA and ATAC data to compute peak-gene correlations; and (iii) pass this set of features to the pretrained model to predict, for a given element-gene pair, whether the element regulates the gene. The resulting predictions provide higher-resolution maps of regulatory landscapes than the corresponding bulk datasets, are cell-type specific, and reflect expected cell-type specific regulation of lineage-specific genes, such as *EBF1*, *CEBPB*, and *SPTA1* (**Fig. 1b-d**, **Fig. S3**).

### Benchmarking of scE2G models

We benchmarked predictions from a total of 14 single-cell models of enhancer-gene regulation, which we applied to the same cell types and datasets. We assessed performance using three consensus pipelines including tasks related to predicting effects of CRISPR enhancer perturbations, fine-mapped eQTLs, and fine-mapped GWAS variants^20^, which together enable evaluating enhancer maps in orthogonal and complementary ways (**Fig. 2a**).

**Figure 2.**
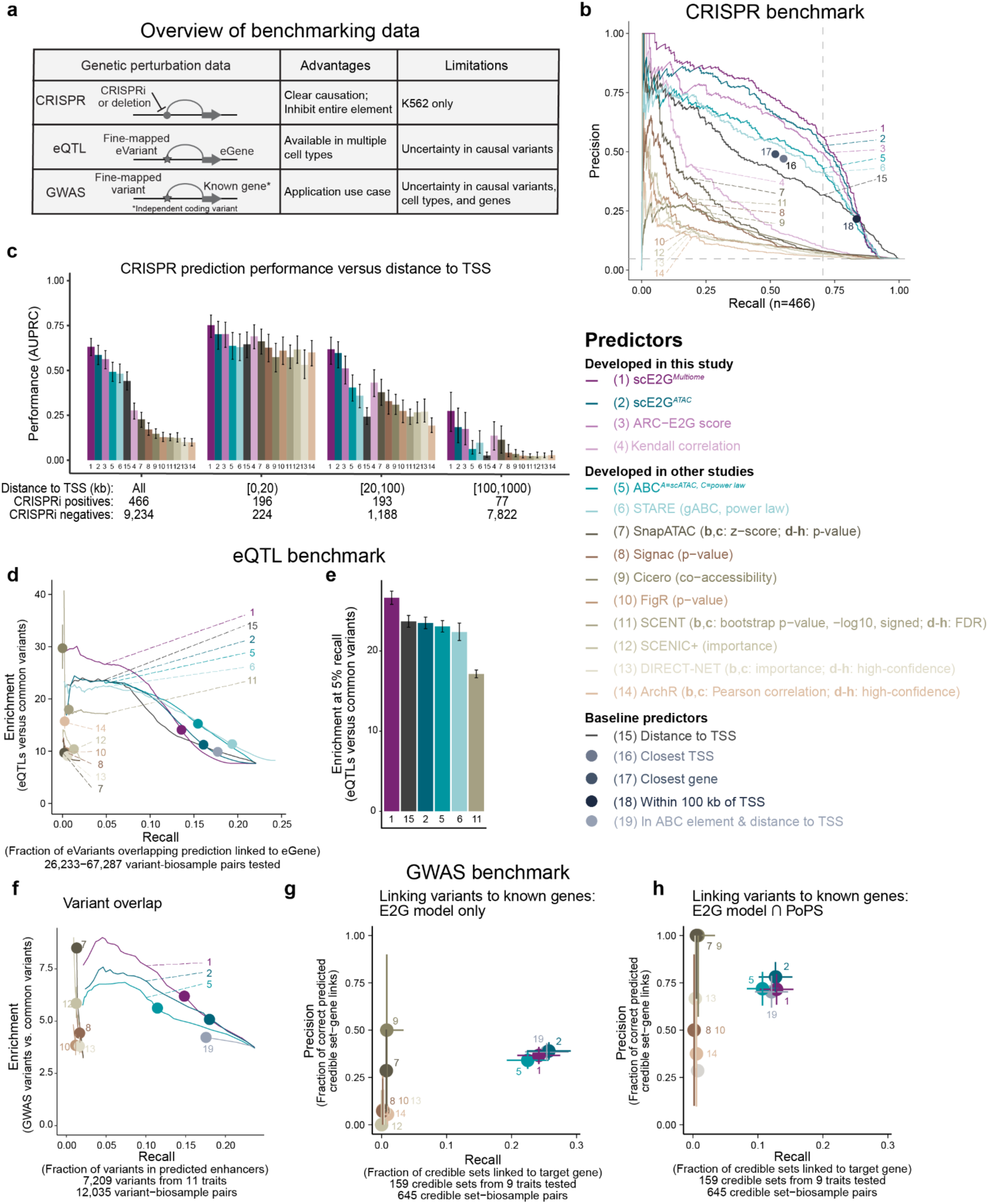
Comparison of scE2G in benchmark tasks. **a.** Overview of CRISPR, eQTL and GWAS benchmarking data (modified from Gschwind *et al.*, 2023^20^) **b.** Precision–recall curves displaying the performance of models on K562 CRISPRi data, which were re-analyzed and assembled in Gschwind *et al.*, 2023^20^ using datasets from Nasser *et al.*, 2021^48^, Gasperini *et al.*, 2019^38^ and Schraivogel *et al.*, 2020^39^. The benchmarking retained only element-gene (E-G) pairs with a TSS distance of less than 1Mb (9,700 tested E-G pairs, 466 positives). Each curve represents a continuous predictor. Each single dot indicates the performance of a binary predictor. Dashed vertical line marks performance at a threshold corresponding to 70% recall; dashed horizontal line indicates the proportion of CRISPRi-positive E-G pairs among all CRISPRi tested E-G pairs (precision value for 100% recall). **c.** CRISPRi benchmarking performance (Area Under the Precision-Recall Curve, AUPRC) of continuous predictors varies with distance to TSS. Error bars show the 95% confidence interval of AUPRC values inferred via bootstrap (1,000 iterations). **d.** Enrichment–recall curves showing performance of models on the eQTL benchmark across matching prediction and eQTL cell type pairs (**Table S4**). Enrichment and recall were aggregated across matching biosamples by summing variant counts across all pairs and were calculated for ∼100 threshold values for continuous predictors. Enrichment (y-axis) is the ratio of fine-mapped distal noncoding eQTLs (PIP > 50%) in predicted enhancers compared to distal noncoding common variants. Recall (x-axis) is the fraction of variants overlapping enhancers linked to the correct gene across the range of score thresholds for enhancers predicted by different predictive models. Points indicate enrichment and recall at suggested thresholds for each model. Error bars at the point represent 95% confidence intervals calculated from the natural log of enrichment. **e.** Enrichment of variants at 5% recall for the same cell type pairs as **d**. Error bars for both plots represent 95% confidence intervals calculated from the natural log of enrichment. **f.** Enrichment–recall curves showing overlap of fine-mapped GWAS variants (PIP > 10%) for 11 blood-related traits in enhancer predictions for 28 paired cell types (**Table S5**). Enrichment (y-axis) is calculated as the ratio of GWAS variants in predicted enhancers compared to distal noncoding variants; recall (x-axis) is calculated as the fraction of variants overlapping enhancers. Performance was evaluated at ∼100 score thresholds for continuous predictors. Points indicate enrichment and recall at suggested thresholds for each model, and error bars represent 95% confidence intervals calculated from the natural log of enrichment. **g.** Precision and recall of models at linking credible sets to known causal genes (inferred from independent signals with coding variants), with enhancers for each model binarized with their suggested threshold. The top two target genes based on model score for each credible set were considered positive predictions. Error bars represent 95% confidence intervals based on the Wilson score interval. Faded points and underlined numbers indicate performance of the models when predictions are intersected with PoPS, as in **h**. **h.** Same as **g**, except enhancer predictions were intersected with the top two genes linked to the credible set by PoPS. Faded points and underlined numbers indicate performance of the models when predictions before intersecting with PoPS, as in **g**.

We compared scE2G models with baselines predictors based on genomic distance and 10 models developed in prior work (**Table S1, Note S1**): (i) ABC-based models ABC^21^ and STARE (gABC)^41^, for which we estimated enhancer activity and 3D contact frequency using pseudobulk ATAC intensity and power law genomic distance-decay, respectively; (ii) correlation-based models Signac^34^, Cicero^29^, ArchR^28^, and FigR^31^, which compute Pearson or Spearman correlation between enhancer accessibility and gene expression (or promoter accessibility); (iii) regression-based models SnapATAC^35^, SCENT^33^, DIRECT-NET^30^, and SCENIC+^32^, which learn a quantitative relationship between the accessibility of one or more enhancers and gene expression. One notable difference among these models is that some (scE2G, ABC, STARE, and SCENT) predict enhancer-gene regulatory interactions by analyzing data from each cell type separately, whereas the remaining models are designed to predict interactions by correlating across all cell types in a dataset simultaneously.

First, we benchmarked the models against the gold-standard CRISPR element perturbation dataset described above. This dataset contains tested element-gene links from epigenetic perturbation experiments in K562 cells and therefore enables evaluating accuracy at predicting causal enhancer-gene regulatory interactions in a given cell type. We generated predictions on a deeply sequenced K562 10x Multiome dataset^40^ (on average 33,387 ATAC fragments and 17,961 RNA UMIs per cell, for a total of 7,821 cells) (**Fig. S2**). For a fair comparison, we restricted predictions to element-gene pairs within 1 Mb, and evaluated against 466 ‘positive’ pairs out of 9,700 CRISPR-validated element-gene pairs (see **Methods**). Because scE2G models were trained on this dataset, we used scores from hold-one-chromosome-out cross-validation in this benchmark. Both scE2G*^Multiome^* and scE2G*^ATAC^* significantly outperformed other existing models on all evaluated metrics, and scE2G*^Multiome^* achieved better performance than scE2G*^ATAC^* (**Table S2**, **Table S3**, delta AUPRC = 0.0255, p_bootstrap_ = 0.0294; delta precision at 70% recall = 1.67%, p_bootstrap_ = 0.065), and scE2G*^ATAC^* achieved better performance than ABC (delta AUPRC = 0.099, p_bootstrap_ = 1 × 10^-16^; delta precision at 70% recall = 7.22%, p_bootstrap_ = 1 × 10^-16^). These trends were more pronounced when considering element-gene pairs separated by larger distances (**Fig. 2a–c**; **Fig. S4**). Notably, only scE2G and other ABC-based models outperformed distance baselines. To confirm the robustness of our analyses, we repeated the benchmark using two other K562 10x Multiome datasets (**Fig. S5**). We also evaluated models using their default settings (**Fig. S6**). We consistently observed that the scE2G models outperformed all other models.

Next, we evaluated how well the models identified distal noncoding eQTL variants and their target genes from matching cell types (**Fig. 2a**, **Fig. S7**). Fine-mapped eQTL variant-gene links provide a complementary source of validation data because eQTLs represent the effects of naturally occurring genetic variation rather than experimentally-induced epigenetic perturbations, and they are currently available in many more cell types and tissues than perturbation data. We compared the enrichment and recall of fine-mapped eQTL variant-gene links from 19 biosamples^42^ in enhancers predicted by each model in 33 matching cell types (*e.g.*, comparing eQTLs in CD8+ T cells with scE2G predictions in CD8+ T cell types from PBMCs and BMMCs, **Fig. S7a**, **Table S4**). In this type of evaluation, we have previously observed recall in the range of 10–20% for models using bulk data (**Fig. N1c**). Higher recall is not expected because of uncertainty in fine-mapping (we consider variants with a posterior inclusion probability >50%) and localization of many fine-mapped eQTLs outside of chromatin accessible elements^20^. Across 67 pairs of matching eQTL and prediction cell types, we found that scE2G achieved higher recall and comparable enrichment to other methods when predictions were binarized by their suggested thresholds (**Fig. 2d**, 14.1-fold enrichment and 13.6% recall for scE2G*^Multiome^* versus 10.1-fold enrichment and 1.3% recall for SCENIC+, the non-ABC-based model with the next highest recall). To compare models agnostic of threshold, we also evaluated enrichment and recall across the full range of scores. At a recall of 5% (the highest achieved by any non-ABC model), variants were significantly more enriched in scE2G*^Multiome^* enhancers than all other models (**Fig. 2e**, 26.6-fold for scE2G*^Multiome^* versus 17.1-fold for SCENT, p_adjusted_=5.29×10^-94^).

Finally, we assessed the ability of the models to enrich for fine-mapped GWAS variants and link credible sets to their target gene, using 15,648 fine-mapped distal noncoding variants for 11 traits (PIP > 10%)^43^ (**Fig. 2a, Fig. S8**). Because GWAS variants impact more highly-constrained genes and tend to be further from their target gene TSS than eQTLs^44^, this benchmark provides an evaluation of the models on a different set of genetic “perturbations”. First, we tested model predictions made on 28 cell types from PBMC and BMMC 10x Multiome experiments against three blood count traits and eight red blood cell traits to evaluate performance on cell types within a single-cell dataset (**Fig. 2f,g,h, Table S5**). For models built from bulk data, we have previously observed a recall of 5–15% and enrichment of 5–15-fold in this type of evaluation^20,45^ (**Fig. N1e**). This low recall is expected due to uncertainty in which variants are causal (considering fine-mapped variants with PIP > 10%), uncertainty in which cell types are causal, and incomplete representation of all relevant cell types. When binarized at the suggested threshold, only scE2G and other ABC-based predictions achieved recall in this range, and they had comparable enrichment of variants to non-ABC-based models (**Fig. 2f**). When evaluating models at linking variants to likely causal genes (nearby genes with independent coding variants associated with the same traits^46^, see **Methods**), we previously saw recall of 20–50% and precision of 25–75%^20^ (**Fig. N1f,g**). scE2G achieved performance in this range, with higher recall and similar precision to non-ABC models (**Fig. 2g**). The same trend was observed for this task when intersecting model-predicted genes with those prioritized by the Polygenic Priority Score (PoPS), which is based on gene function^46^. scE2G*^Multiome^* achieved a recall of 24.2% and 12.9% before and after intersecting with PoPS, while non-ABC models did not surpass a recall of 0.8% in either case (**Fig. 2h**).

Overall, scE2G models outperform other single-cell models across multiple benchmarking tasks and achieve similar performance to the ENCODE-rE2G model based on bulk DNase-seq data (**Note S2**). Each benchmarking dataset has its advantages and limitations: (i) CRISPR element perturbations most directly identify links between enhancers and genes, by characterizing the effects of inhibiting an entire element on gene expression in a specific cell type. However, this CRISPR dataset is limited to K562 cells, and contains some element-gene links likely attributable to indirect effects or mechanisms other than enhancers^21^. (ii) Fine-mapped eQTL data are available in many primary cell types, but there is a level of uncertainty in identifying the causal variant, and the absolute recall is low because eQTL variants can act through other mechanisms beyond affecting enhancers^47^. (iii) Interpreting GWAS signals is an important use case, but, for the purposes of benchmarking models of enhancer-gene regulatory interactions, there is uncertainty in causal variants, causal cell types, and the “known” genes as annotated by independent coding variants^46^ (**Fig. 2a**). The fact that scE2G performs well across these three orthogonal types of perturbation-based benchmarking datasets, covering many cell types and input datasets, suggests that scE2G has indeed learned predictive and generalizable rules of enhancer-gene regulation.

### Model interpretation

While developing the scE2G models, we analyzed the importance of model features to identify a minimal set of features needed for good performance and thereby increase their applicability in practice. We performed three feature importance analyses using the CRISPR benchmark, including evaluating all subsets of the 11 input features, sequential feature selection, and ablation-based feature importance (**Fig. 3a, Fig. S9**). In each analysis, the ABC score, Kendall correlation, and promoter class (whether the gene is ubiquitously expressed) had large significant effects on AUPRC and precision at 70% recall. We selected these features along with 4 others related to genomic distance and ATAC signal that yielded optimal performance for the final scE2G*^Multiome^* model (**Fig. 3a**).

**Figure 3.**
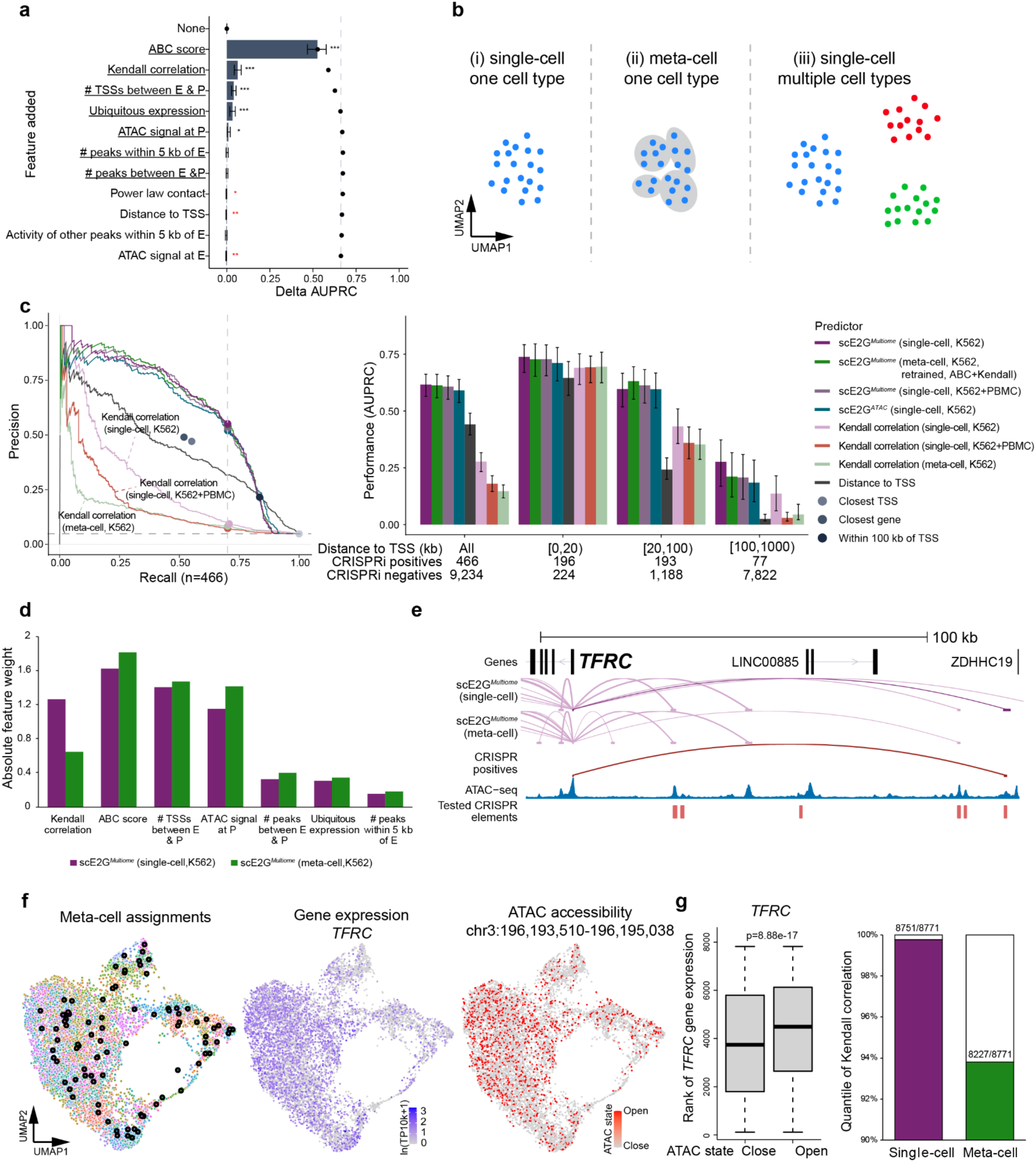
Interpretation of scE2G and the Kendall correlation. **a.** Forward sequential feature selection for scE2G*^Multiome^*. Bars represent the change in AUPRC from the previous model including features above it. Dots indicate the accumulated change in AUPRC for the model including the feature labeled and all above it, and the dashed vertical line represents the total accumulated AUPRC across all features. Error bars represent 95% range of AUPRC values inferred via bootstrap (1,000 iterations). One, two, and three stars indicate p-values less than 0.05, 0.01, and 0.001, respectively; black stars represent a positive change in AUPRC and red stars represent a negative change in AUPRC. The underlines indicate the features in the final model. **b.** Overview of three distinct approaches to calculating Kendall correlation between gene expression and ATAC accessibility across (i) single-cells within one cell type, (ii) meta-cells within one cell type, and (iii) single-cells across multiple cell types. **c.** CRISPR benchmarking of the three approaches to calculating Kendall correlation (**b**) and corresponding scE2G models. **Left**: Precision–recall curves display the performance of models on the K562 CRISPRi data. As in Fig. 2b, each curve represents a continuous predictor. Each single dot indicates the performance of a binary predictor. Dashed vertical line marks performance at a threshold corresponding to 70% recall; dashed horizontal line indicates the proportion of CRISPRi^+^ E-G pairs among all CRISPRi validated E-G pairs (precision value for 100% recall). **Right**: CRISPRi benchmarking performance (AUPRC) of continuous predictors varies with distance to TSS. Error bars show the 95% confidence interval of AUPRC values inferred via bootstrap (1,000 iterations). **d.** Barplots showing the comparison of absolute feature weights between scE2G*^multiome^* models based on single-cell and meta-cell data. **e.** scE2G*^Multiome^* (single-cell) and scE2G*^Multiome^* (meta-cell) predictions for *TFRC* in K562 for the region chr3:196,070,000-196,197,500 (hg38). Enhancer-gene linking arcs in purple are shown for scE2G*^Multiome^* (single-cell) and scE2G*^Multiome^* (meta-cell) predictions binarized according to the 70% recall threshold in **c**, and the width of the arc indicates the magnitude of the scE2G score. The enhancer-gene linking arc in red indicates the positive CRISPRi enhancer at chr3:196,193,510-196,195,038 for *TFRC* in this region, and the corresponding prediction made only by scE2G*^Multiome^* (single-cell) is in a darker purple. Normalized ATAC signal for K562 (Xu *et al.*) and all tested CRISPR elements are also shown. **f.** The expression of gene *TFRC* and the accessibility of enhancer chr3:196,193,510-196,195,038 indicate stochastic transcription bursting and associated enhancer accessibility across single cells. **Left**: Assignment of single cells to meta-cells using SEACells^50^. Each point represents a single cell, and each circle indicates a meta-cell. Meta-cells and their corresponding single cells are color-coded with the same color. **Middle**: Normalized single-cell gene expression levels of *TFRC*. **Right**: Binarized single-cell ATAC accessibility. **g.** Comparison of Kendall correlation of *TFRC* gene expression and chr3:196,193,510-196,195,038 enhancer accessibility between single-cell and meta-cell data. **Left**: Box plot comparing the rank of gene expression between single cells with open and closed ATAC states. Wilcoxon rank-sum test. **Right**: Barplot showing the quantile of Kendall correlation based on single-cell and meta-cell data across 8,771 E-G pairs available. The ranks are in ascending order (larger ranks corresponding to larger values).

Compared to the bulk ENCODE-rE2G model^20^, the major feature added to the scE2G*^Multiome^* model was the Kendall correlation between element accessibility and gene expression in single cells. Incorporating the Kendall correlation improved the model’s ability to predict long-range enhancer-gene regulatory interactions (*e.g.*, >100 kb, **Fig. 2c, Fig. S9c**), significantly improving over the ABC score (delta AUPRC = 0.214, p_bootstrap_ = 1×10^-16^) or the scE2G*^ATAC^* model (delta AUPRC = 0.0917, p_bootstrap_ = 3×10^-4^).

For the scE2G*^Multiome^* model, the Kendall correlation is calculated across single cells within a single cell type or state (**Fig. 1a**). This approach differs from previous correlation- and regression-based methods, such as Signac^34^ and SnapATAC^35^, that quantify enhancer-gene associations across single cells or meta-cells across multiple cell types in a single dataset^28–32,34,35^. To directly compare these approaches, we computed three Kendall correlation metrics between enhancer accessibility and gene expression: (i) across single cells within a single cell type (single-cell, K562); (ii) across single cells that include multiple cell types (single-cell, K562+PBMC); and (iii) across meta-cells within a single cell type (meta-cell, K562) (**Fig 3b**). The Kendall correlation calculated across single cells within a cell type (single-cell, K562) performed better than the other two metrics when tested alone and when integrated into the scE2G*^Multiome^* model (**Fig. 3c**). While performance differences among scE2G*^Multiome^* models were smaller, the retrained meta-cell scE2G*^Multiome^* model had down-weighted the Kendall correlation feature (**Fig. 3d**), leading to worse predictive performances for long-distance enhancer-gene pairs (**Fig. 3c**). This suggests that stochastic variation in enhancer-gene co-activities across single cells within a cell type is informative for linking enhancers to their target genes. Consistent with this interpretation, enhancer-gene pairs that were correctly predicted by the scE2G*^Multiome^* model but not the scE2G*^ATAC^* model, which does not use the Kendall correlation, did not show any apparent K562 cell subtype clustering (**Fig. 3e,f**) or correlated enhancer and gene activity across K562 meta-cells (**Fig. 3g**). These correlations may result from a temporal relationship between when an enhancer is accessible and when a gene is transcribed^49^. Therefore, the improved performance of scE2G*^Multiome^* over scE2G*^ATAC^* appears to stem from the detection of regulatory heterogeneity across single cells within a cell type by the Kendall correlation. Models that consider enhancer-gene correlation across cell types^28–32,34,35^ may not capture such a temporal relationship.

### Performance across sequencing depths

Models that will be used to predict interactions across cell types should be robust to variation in sequencing depth. This is because depth can vary dramatically both across datasets as well as across cell types within a single dataset. For example, the datasets used for benchmarking (**Fig. 2**) included cell types with between 3,960 to 37,442 ATAC fragments/cell and 108 to 42,072 cells, leading to a difference of over three orders of magnitude in the total number of ATAC fragments per cell type.

To systematically evaluate the robustness of scE2G to variation in sequencing depth and cell counts, we downsampled K562 10x Multiome data^40^ across cell counts as well as across ATAC fragments/cell and RNA UMIs/cell, yielding datasets with 100 to 7,821 cells; 100,000 to 261 million total ATAC fragments; and 50,000 to 600 million total RNA UMIs (**Fig. S10a**). We evaluated scE2G predictions on each downsampled dataset using the CRISPR benchmark (**Fig. 4, Fig. S10**).

**Figure 4.**
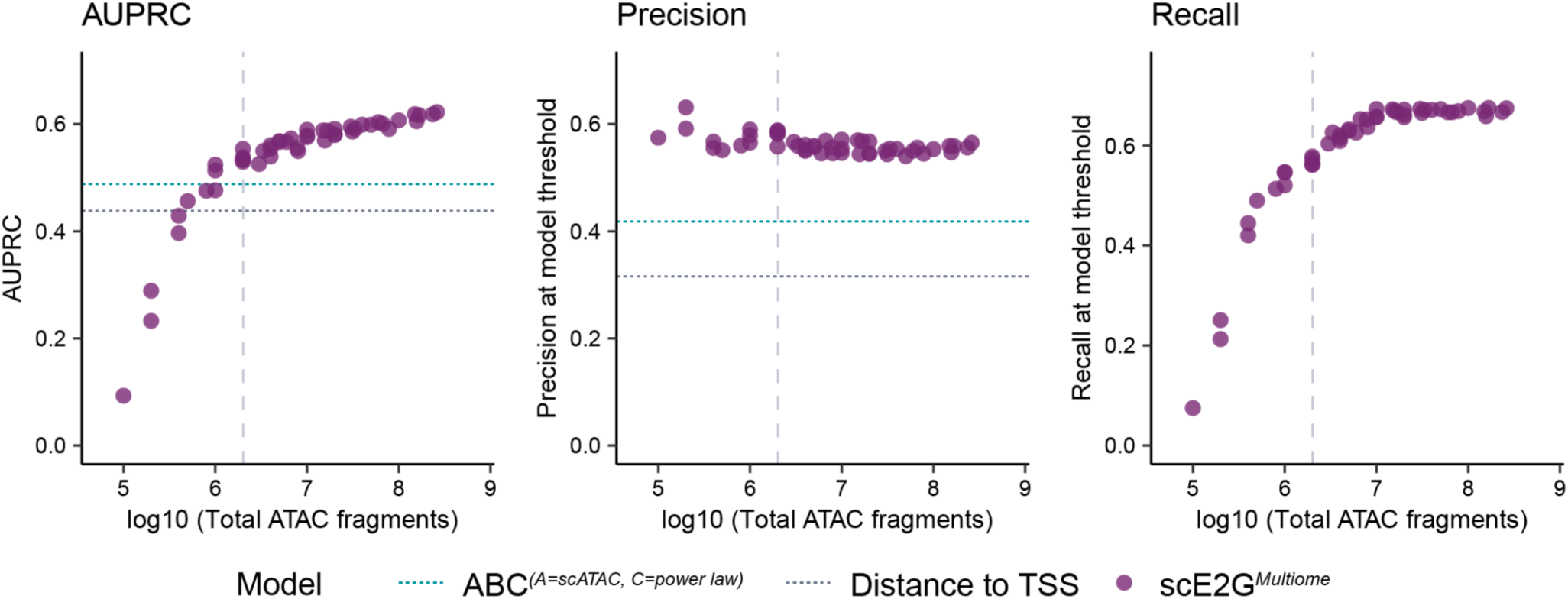
Performance of scE2G across sequencing depths. AUPRC (left), precision at the model threshold (center) and recall at the model threshold (right) of scE2G*^Multiome^* on the CRISPR benchmark across ATAC fragment counts. Performance of distance to TSS (grey line) and ABC on the full-depth dataset (teal line, the next best model after scE2G*^Multiome^* and scE2G*^ATAC^*) are shown for reference. These reference lines are not shown for recall because the score threshold for models is defined based on 70% recall. The vertical dashed lines indicate 2 million fragments (and 1 million RNA UMIs).

Above a threshold of 2 million ATAC fragments and 1 million RNA UMIs, the precision of scE2G*^Multiome^* was stable, and both scE2G*^Multiome^* and scE2G*^ATAC^* significantly outperformed genomic distance and all other tested models applied to the full dataset, by both AUPRC and precision (**Fig. S10b,c**). Below 2 million ATAC fragments, the models were unstable and recall showed a dramatic decrease. This is because the quality of ATAC peak calls (with MACS2) degrades, for example missing >25% of tested CRISPR elements (**Fig. S10b**). Above 2 million ATAC fragments, both AUPRC and recall increased slightly, up to and beyond 100 million fragments. This appeared to be due to an increased utility of the Kendall correlation feature, which has worse performance at lower coverages due to counting noise (also true for other correlation-based approaches, **Fig. S11**). In developing scE2G, we accounted for this variation by combining the ABC scores and Kendall correlations into a single feature for input into the logistic regression (see **Methods**, **Fig. S12)**.

Validating the stability of scE2G across cell types of differing cell count and sequencing depth, we observe a consistent number of base pairs included in enhancers for thresholded predictions for cell types with over 2 million fragments, varying from 33 to 57 Mb for scE2G*^Multiome^*. In contrast, the enhancer set sizes for non-ABC-based models vary much more with fragment counts, ranging over 3 to 4 orders of magnitude (**Fig. S11**).

To summarize, the final scE2G models require a minimum of 2 million total ATAC fragments and 1 million RNA UMIs (corresponding to at least 100 single cells), above which performance is robust to sequencing depth (**Fig. 4, Fig. S10, Fig. S11**). Performance slightly increases with more total ATAC fragments, up to and beyond 100 million fragments per cell type.

### Cell-type resolution of scE2G predictions

We next explored the properties of scE2G predictions across cell types from three single-cell multiome datasets (PBMCs, BMMCs, and pancreatic islets) (**Fig. S13**, **Table S6**, **Table S7**). 39 cell types had more than the recommended 2 million ATAC fragments (**Fig. S13a**, range: 4.9 million–822 million). In each of these cell types, scE2G*^Multiome^* predicted an average of 54,032 enhancer-gene regulatory interactions (**Fig. S13b**, range: 46,486 to 60,024), including an average of 40,160 unique enhancers per cell type (**Fig. S13c**) and 8,533 genes with at least one distal enhancer (**Fig. S13f**). The average enhancer was located 94.9 kb from the promoter of its target gene (**Fig. S13h**). Across all cell types, scE2G*^Multiome^* predicted 6.3 enhancers to regulate each expressed gene, and each enhancer to regulate 1.4 genes (**Fig. S13i,j**).

To validate that scE2G predictions reflect the expected cell-type specificity, we analyzed the similarity of enhancer-gene predictions across all cell types. We computed two metrics to quantify the similarity of each pair of binarized predictions: the Jaccard similarity, which represents the fraction of shared enhancer-gene predictions between two cell types, and the Pearson correlation, which represents how consistent the scE2G scores for each prediction are between two cell types (**Fig. 5a)**. We observed higher similarity between closely-related cell types by both measurements, indicating that the set of predicted enhancer-gene pairs and the strength of the scE2G score capture cell-type specific patterns both within a dataset from one tissue and between similar cell types across datasets (**Fig. S14a**).

**Figure 5.**
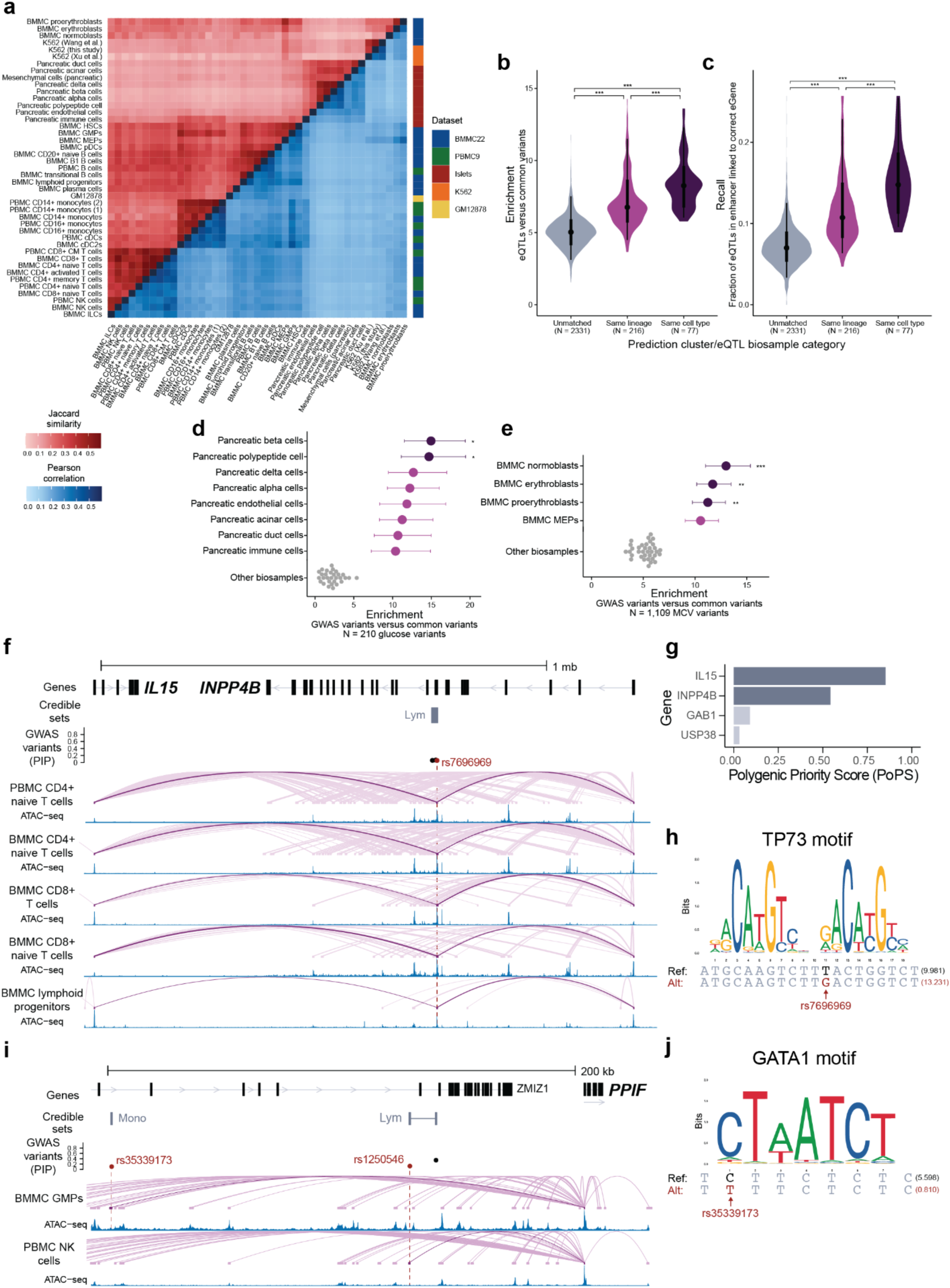
Building and interpreting maps across tissues. **a.** Heatmap of the Jaccard similarity (upper left, red) and Pearson correlation (lower right, blue) between scE2G*^Multiome^* enhancer-gene predictions with scores above the suggested threshold between predictions for each pair of 43 cell types. An enhancer-gene link in which the element overlapped by at least 50% in two sets of predictions was considered shared. **b.** Distribution of enrichment of eQTLs in scE2G*^Multiome^* predictions from biosamples of the same cell type, same cell lineage, or unmatched biosamples (see **Methods**). Enrichment was computed as the fraction of fine-mapped distal noncoding eQTLs (PIP > 50%) in predicted enhancers compared to distal noncoding common variants. Three stars indicate a p-value less than 0.001 from a two-sided t-test between distributions, adjusted for multiple comparisons with the Bonferroni correction. See Methods, ‘Data visualization’ section for definition of point interval plot elements. **c.** Same as **b**, but comparing distribution of recall of eQTLs in scE2G*^Multiome^* predictions. Recall was computed as the fraction of fine-mapped distal noncoding eVariant–eGene links (PIP > 50%) for which the eVariant is located in a predicted enhancer linked to the correct eGene. Stars indicate significance as described in **b**. See **Methods**, ‘**Data visualization**’ section for definition of point interval plot elements. **d.** Enrichment of fine-mapped GWAS variants associated with random blood glucose compared to common variants in scE2G*^Multiome^* predictions from different cell types. Stars indicate that enrichment of variants in pancreatic beta and polypeptide cells (dark purple) was significantly higher than enrichment in predictions from all other cell types evaluated (grey) except for six other pancreatic islet cell types (light purple) with all padjusted < 0.05. Error bars represent 95% confidence intervals computed from the natural log of enrichment. **e.** Same as **d**, but for fine-mapped GWAS variants associated with mean corpuscular volume (MCV). Stars indicate that enrichment in the indicated cell type (dark purple) was significantly greater than enrichment in predictions from all other cell types evaluated (grey) except for megakaryocyte-erythroid progenitors (MEPs), with two stars indicating all padjusted < 0.01 and three stars indicating all padjusted < 0.001. **f.** scE2G*^Multiome^* predictions and normalized ATAC signals for *IL15* and *INPP4B* in several T and lymphoid progenitor cell types for the region chr4:141,612,977-142,900,000 (hg38). Fine-mapped GWAS variants with PIP > 10% and credible sets for lymphocyte count are shown. The variant rs7696969 is indicated in red, and predictions overlapping this variant are shown in darker purple. The width of prediction arcs is proportional to scE2G score. **g.** Polygenic Priority Scores (PoPS) ranking genes within 1 Mb of rs7696969 for lymphocyte count. **h.** Logo showing position weight matrix for the TP73 motif (MA0861.1) altered by the reference and alternate alleles of rs7696969. The number after each sequence is the motif match score computed with FIMO^65^. The match to the reference allele has a p-value of 3.8×10^-5^, and the match to the alternate allele has a p-value of 9.23×10^-6^. **i.** scE2G*^Multiome^* predictions and normalized ATAC signals for *PPIF* in granulocyte-macrophage progenitors and natural killer cells for the region chr10:79,136,379-79,375,225 (hg38). Fine-mapped GWAS variants with PIP > 10% and credible sets for monocyte count and lymphocyte count are shown. The variants rs35339173 (monocyte count) and rs1250546 (lymphocyte count) are indicated in red, and predictions overlapping these variants are shown in darker purple. The width of prediction arcs is proportional to scE2G score. **j.** Logo showing position weight matrix for the GATA1 motif (MA0035.5)^66^ altered by the reference and alternate alleles of rs35339173. The number after each sequence is the motif match score computed with FIMO^65^. The match to the reference allele has a p-value of 2.41×10^-3^, and the match to the alternate allele has a p-value of 0.017.

Next, we asked whether noncoding, fine-mapped eQTL variants from 64 biosamples were more highly enriched in the scE2G*^Multiome^* enhancer predictions for matching cell types, which would indicate that cell-type specificity of predictions correspond to actual cell-type specific regulatory landscapes. We found that eQTLs were indeed better predicted by scE2G*^Multiome^* when applied to the same cell type (*e.g.* eQTLs from CD4+ T cells in predictions from any T cell type) and of the same lineage (*e.g.* eQTLs from CD4+ T cells in predictions from any lymphoid cell type) than in unmatched cell types (**Fig. 5b,c**, **Fig. S14b**, mean enrichment of 8.36-fold versus 5.07-fold, p_adjusted_ = 5.39×10^-27^; mean recall of 15.4% versus 7.2% of 8.36-fold versus 5.07-fold, p_adjusted_ = 8.01×10^-28^).

The cell-type resolved predictions of the scE2G*^Multiome^* indeed expanded the number of unique enhancers detected across all cell types, versus what could be identified from bulk or pseudobulk samples. For example, to quantify this, we considered three related B cell types from BMMCs: B1 B cells, naive CD20+ B cells, and transitional B cells. Applying scE2G*^Multiome^* to each of the three B cell types individually identified a total of 105,774 enhancer-gene regulatory interactions, 4.7% more than observed when applying scE2G*^Multiome^* to similar numbers of cells randomly sampled from the combination of all three cell types (90/100 replicates, **Fig. S14d, Methods**).

At individual genes with known cell-type specific patterns of expression, scE2G predictions of enhancer-gene regulatory interactions indeed highlighted relevant cell types. For example, *EBF1* showed many predictions in B cells, and *CEBPB* showed many predictions in myeloid cells (**Fig. 1**, **Fig. S3**). In contrast, scE2G applied to the pseudobulk of all cells in the same dataset obscured this variability (**Fig. 1**, **Fig. S3**).

Overall, we demonstrate that scE2G can identify many enhancer-gene links specific to individual cell types that are missed when analyzing aggregated cell populations.

### Linking GWAS variants to target genes

Having established confidence in the cell-type specificity of scE2G enhancer-gene predictions, we applied scE2G enhancer-gene maps in 40 cell types to interpret fine-mapped GWAS variants from 17 relevant traits.

We first examined the global enrichment of fine-mapped GWAS variants in elements linked to genes by scE2G*^Multiome^* and found the strongest enrichment in expected cell types (**Fig. S14d**). For example, variants associated with random blood glucose were most enriched in enhancers for pancreatic beta and polypeptide cells (enrichment = 15.0-fold and 14.7-fold) and both were significantly more enriched than all non-pancreatic islet enhancers (**Fig. 5d**, all p_adjusted_ < 0.05). On the other hand, variants associated with mean corpuscular volume were most enriched for enhancers in normoblasts, erythroblasts, and proerythroblasts (average enrichment = 12.0-fold) and all were significantly more enriched than enhancer for any cell type except for megakaryocyte-erythroid progenitors (**Fig. 5e**, all p_adjusted_ < 0.01). We also used stratified linkage disequilibrium score regression (S-LDSC)^51,52^ to identify cell types whose enhancers were enriched for genome-wide heritability of GWAS traits, and observed similar trends as with enrichment of fine-mapped variants (**Note S3**, **Table S8**).

We used scE2G*^Multiome^* to nominate causal gene(s) and cell type(s) for 1,906 noncoding credible sets (associated with traits with at least a 5-fold significant enrichment in predicted enhancers from at least one cell type) comprising 3,705 variants (PIP > 10%), considering the top two genes linked to enhancers overlapping variants based on scE2G score across all cell types (**Table S9**). In total, scE2G nominated 1,470 fine-mapped variants overlapping enhancers linked to 1,366 genes, leading to 2,925 unique credible set-gene-cell type predictions. Notably, 500 credible sets were most strongly linked to genes that did not have the closest transcription start site to a variant in the credible set.

scE2G predictions at the 4q31.21 locus generated variant-to-function hypotheses and illustrate the ability of scE2G to make long-range predictions. At this locus, rs7696969 is associated with lymphocyte count (fine-mapping PIP = 22%) and is a lead variant associated with platelet distribution width (p = 5.4×10^- 48^)^53^. The variant, which is in an intron of *INPP4B,* overlaps an enhancer predicted by scE2G to regulate both *INPP4B* and another nearby gene, *IL15*, in many lymphatic cell types, with the strongest predictions in natural killer and T cells, and weaker predictions in hematopoietic stem cells (**Fig. 5f, Fig. S15a**). Both promoters are located far from the variant (441 kb for *INPP4B* and 769 kb for *IL15*). Both genes have plausible roles in lymphocyte counts. *INPP4B* (inositol polyphosphate 4-phosphatase type II) acts upstream of the AKT pathway, and *Inpp4b* knockout in murine hematopoietic cells affects the abundance of certain B cell populations^54^. *IL15* encodes a cytokine required for natural killer and memory CD8+ T cell development and maintenance^55^. By an independent GWAS gene prioritization method, PoPS^46^, both genes have similarly high scores, stronger than other genes in the locus (ranks 1 and 2 for *IL15* and *INPP4B*, respectively) (**Fig. 5g**). In addition, both IL15 and INPP4B have been shown to mediate erythropoietin-induced erythropoiesis in murine hematopoietic stem cells^56,57^. Supporting a regulatory role for this variant, rs7696969 is both a fine-mapped eQTL for *INPP4B* (PIP = 66.6%) and a chromatin accessibility QTL in lymphoblastoid cell lines^42,58^. The variant also overlaps transcription factor footprints for TP73 in multiple T cell types and alters a TP73 motif (**Fig. 5h**). Together these data suggest that rs7696969 might affect multiple blood traits through cell-type specific, long-range regulation of *INPP4B* and *IL15*.

scE2G also nominated variant-to-function relationships in cases where bulk models have not. For example, two variants at the 10q22.3 locus, one associated with monocyte count (rs35339173, PIP = 10.4%) and one associated with lymphocyte count (rs1250546, fine-mapping PIP = 12.3%), fall within introns for *ZMIZ1* but have unknown causal genes. Neither variant overlaps an enhancer from bulk ENCODE-rE2G predictions in over 1,400 biosamples. However, using scE2G, we found that both variants fall within predicted, cell-type-specific enhancers for *PPIF*: the monocyte count variant in granulocyte-monocyte progenitors and the lymphocyte count variant in PBMC natural killer (NK) cells (**Fig. 5i**, **Fig. S15b**). Supporting a regulatory role for each variant, the monocyte count variant disrupts a motif for GATA1, a transcription factor known to regulate hematopoiesis (**Fig. 5j**), and the lymphocyte count variant overlaps an inferred transcription factor footprint for GLI2 in NK cells. Furthermore, several studies on the biological role of *PPIF*, a gene encoding the widely-expressed protein cyclophilin D which regulates metabolism and cell death, support its prioritization by scE2G for these variants^59^. CRISPRi knockdown of *PPIF* expression in the monocytic cell line THP1 has been shown to affect metabolic state of mitochondria, which is known to affect monocyte activation and differentiation^60^. NK cells from *Ppif* knockout mice exhibit phenotypic immaturity^61^. *Ppif* also has a critical role in regulating hematopoietic stem cell function, proliferation, and differentiation^62–64^. In sum, these data illustrate the utility of scE2G in nominating causal genes and cell types affected by GWAS variants.

## Discussion

Here we present scE2G, a new family of models to predict enhancer-gene regulatory interactions from single-cell data. By making accurate inferences of enhancer-gene regulatory interactions from single-cell datasets, scE2G will enable building maps across many cell types and states to understand gene regulation, developmental biology, and disease-related genetic variation.

scE2G models use a distinct architecture from previous single-cell models, yielding improvements in accuracy and stability. While most existing models rely on either (1) correlation or regression between enhancer accessibility and gene expression or accessibility or (2) a simple combination of accessibility and distance, scE2G uses a supervised learning framework that involves training directly on CRISPR data. This approach learns a combination of features, including the ABC score, element-gene correlation, and promoter class, that substantially outperforms any individual feature or model. We introduce a new strategy for capturing element-gene correlations, which appears to reflect stochastic temporal relationships between element accessibility and gene expression across single cells. This metric applies the non-parametric Kendall correlation across single cells within a given cell type, and outperforms correlation approaches that either combine single cells into meta-cells within a cell type, or compare single cells across distinct cell types. This correlation provides insights into the regulatory roles of enhancers, and notably can only be captured by paired multimodal single-cell assays.

scE2G achieves stable performance across different datasets with different sequencing depths, and across cell types with different frequencies within the same dataset. This is achieved, in part, by integrating the element-gene correlation metric (which is sensitive to sequencing depth) with the ABC score (which is less so). This method and accompanying guidelines (see **Code Availability**) will enable building maps of enhancers across many additional datasets.

Our study applies and extends a benchmarking framework we previously developed with the ENCODE Consortium^20^. This approach combines three orthogonal genetic perturbation datasets, each with its own set of strengths and limitations, to test the generalizability of models across cell types and prediction tasks. Here, we extended this framework by curating additional eQTL datasets and GWAS traits with matched single-cell datasets. We provide these benchmarking pipelines and curated datasets to facilitate the development and consistent evaluation of future models (see **Code Availability**).

scE2G maps of enhancer-gene regulatory interactions will improve interpretation of noncoding variants associated with complex traits, by enabling a more comprehensive assessment of all cell types and states in a tissue. Here we illustrate this by interpreting variants associated with metabolic and blood cell traits, and identify novel variant-gene links to *IL15* and *INPP4B* and to *PPIF*. We note that, in interpreting GWAS loci, we expect that predictions linking variants to target genes should be combined with orthogonal information about gene function, as in prior work^20,46,67,68^. Consistent with this, combining scE2G with PoPS^46^ increases precision at identifying silver-standard causal genes (**Fig. 2g,h**).

We note several limitations and opportunities for further development. First, scE2G is designed to predict the effects of distal elements that act as enhancers, and not the effects of elements that act through other mechanisms, such as topological elements that bind CTCF. Second, the CRISPR perturbation data upon which scE2G is trained contains only 472 positive element-gene links and is limited to K562 cells. Expanding this training dataset by characterizing true regulatory interactions in additional cell types should enable further improvements to scE2G. Third, while scE2G models achieve performance similar to that of models based on bulk DNase-seq data (ENCODE-rE2G^20^), we have observed that even more accurate models can be achieved by incorporating additional bulk epigenomic assays such as cell-type specific Hi-C data or H3K27ac ChIP-seq data^20^. Thus, we have not yet reached the maximum performance achievable in this dataset. Fourth, although we leverage eQTL and GWAS data for orthogonal benchmarking of scE2G, optimal performance at these prediction tasks will require additional models that consider information beyond element-gene links, for example by considering the effects of single-nucleotide variants on elements and the effects of genes on phenotypes.

In summary, this study presents a state-of-the-art model to predict enhancer-gene regulatory interactions from single-cell data and demonstrates its utility in interpreting the function of genetic variation. The scE2G models can be applied to datasets of varying sizes and sequencing depth, and can build comprehensive and accurate maps of enhancer-gene regulatory interactions across thousands of cell types and states in the human body. For example, in parallel work we have applied scE2G to single-cell atlases of the human coronary arteries^69^ and human fetal heart^70^, identifying and validating enhancers containing variants linked to cardiovascular diseases. By creating these maps of enhancer landscapes, we expect scE2G to provide the foundation for a widely-useful approach to study many common diseases and complex traits.

## Data Availability

The data sources for each 10x Multiome sample are listed in **Table S10**. The locations of the scE2G*^Multiome^* prediction files are listed in **Table S6**.

## Code Availability

scE2G: https://github.com/EngreitzLab/scE2G/releases/tag/v1.0

CRISPR benchmarking:https://github.com/EngreitzLab/CRISPR_comparison/tree/main

eQTL benchmarking: https://github.com/EngreitzLab/eQTLEnrichment/tree/integrated

GWAS benchmarking: https://github.com/EngreitzLab/GWAS_E2G_benchmarking

Data analysis for this study: https://github.com/anderssonlab/scE2G_analysis/releases/tag/v1.0

## Acknowledgements

M.U.S. acknowledges the support of an NSF Graduate Research Fellowship (DGE-1656518) and a graduate fellowship award from Knight-Hennessy Scholars at Stanford University. A.R.G. and L.M.S acknowledge the support of R01HG011664 and the NHGRI Impact of Genomic Variation on Function Consortium (UM1HG011972). B.L.G. acknowledges support from the Novo Nordisk Foundation Center for Genomic Mechanisms of Disease (NNF21SA0072102). D.A. acknowledges support from an AHA Postdoctoral Fellowship (821920 and 23POSTCHF1019753) and the Stanford Maternal and Child Health Research Institute (MCHRI) Postdoctoral fellowship. C.S.M. and A.T.S. acknowledge support from the NHGRI Impact of Genomic Variation on Function Consortium (UM1HG012076). B.U. acknowledges the NIH Bridge2AI Center Grant U54HG012510. A.Kundaje acknowledges support from the NHGRI Impact of Genomic Variation on Function Consortium (U01HG012069). R.A. acknowledges support from the Novo Nordisk Foundation (NNF20OC0059796) and the Novo Nordisk Foundation Center for Genomic Mechanisms of Disease (NNF21SA0072102). J.M.E. acknowledges support from the NHGRI Impact of Genomic Variation on Consortium (UM1HG011972); the Novo Nordisk Foundation Center for Genomic Mechanisms of Disease (NNF21SA0072102); NHLBI R01HL159176; the NHGRI Genomic Innovator Award (R35HG011324); the Applebaum Foundation; Gordon and Betty Moore and the BASE Research Initiative at the Lucile Packard Children’s Hospital at Stanford University; and the Chan Zuckerberg Initiative DAF (2022-249191), an advised fund of Silicon Valley Community Foundation. We thank Stanford University and the Stanford Research Computing Center for providing computational resources and support as part of the Sherlock High-Performance Compute Cluster.

We thank members of the Andersson lab, Engreitz lab, IGVF Noncoding Variants Focus Group, and the Novo Nordisk Foundation Center for Genomic Mechanisms of Disease for feedback on the scE2G model and results. We thank Kushal Dey, Elizabeth Dorans, Alkes Price, Alireza Karbalayghareh, and William Greenleaf for discussions and feedback on the manuscript.

## Author Contributions

W.Q. and M.U.S. co-led the project as co-first authors, and agreed that either author can be listed first in reporting the manuscript. W.Q. developed the Kendall correlation metric and ARC-E2G score, computed previous models, conducted downsampling analysis, analyzed multiome data, and benchmarked these methods versus CRISPR data. M.U.S. trained the scE2G models, conducted feature selection and interpretation, developed and applied eQTL and GWAS benchmarking pipelines, conducted downsampling analysis, compared predictions across cell types, and applied scE2G maps to interpret

GWAS variants. M.U.S., W.Q., and A.S.T. developed the software package to compute scE2G. A.R.G., L.S., and J.M.E. developed the CRISPR benchmarking pipeline and dataset. X.R.M. and D.D. contributed to development and validation of scE2G models and peak-calling methods. E.J. contributed to data interpretation and application of scE2G. H.E. contributed to comparison of correlation metrics. W.Q., B.L.G., and T.R.J. implemented software to compute Kendall correlation. C.S.M., D.A., and A.T.S. generated 10x Multiome data. B.U. contributed to strategy for feature selection. A.K. contributed to development of peak-calling methods. W.Q., M.U.S., X.R.M., A.R.G., E.J., J.M.E., and R.A. analyzed and interpreted data in the development of scE2G. M.U.S., W.Q., R.A., and J.M.E. wrote the manuscript with input from all authors. R.A. and J.M.E. provided overall supervision for the project, contributed equally as co-last authors, and agreed that either author can be listed last in reporting the manuscript.

## Supplementary Information

### Supplementary Figures

**Fig. S1.**
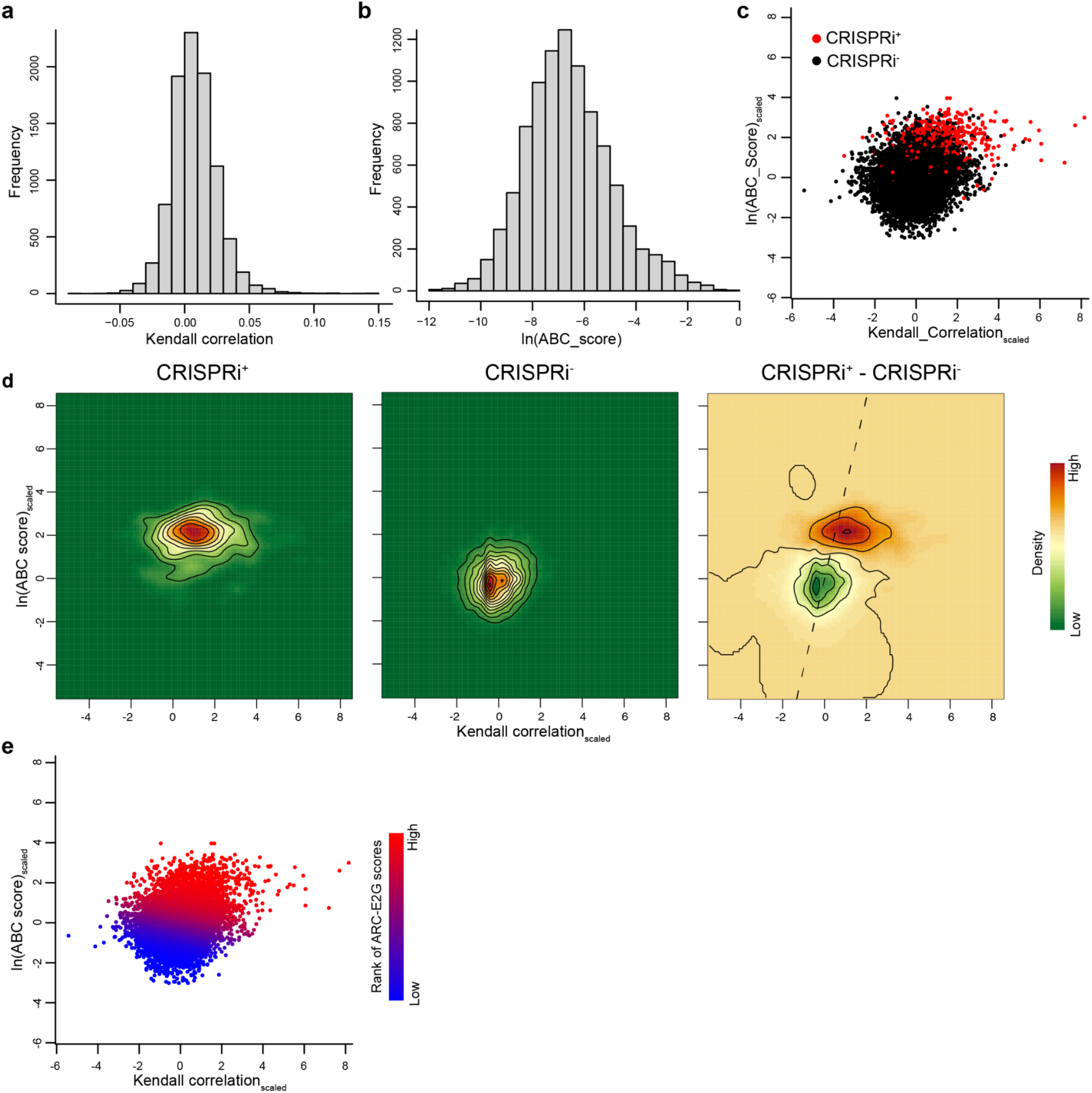
Integrating ABC score and Kendall correlation into ARC-E2G score. **a** and **b.** Histogram plots showing the distribution of the element-gene Kendall correlation (**a**) and ln(ABC_score) (**b**) in CRISPRi validated enhancer-gene pairs. **c.** Scatter plot showing the relationship between scaled ln(ABC_score) and scaled Kendall correlation. We scaled values by subtracting the mean and then dividing by the standard deviation. Each point represents a CRISPRi validated enhancer-gene pair. Red: Element-gene pairs with significant effects in the CRISPR experiment (CRISPRi^+^). Black: Element-gene pairs that did not have significant effects in the CRISPR experiment(CRISPRi^-^). **d.** Left: Density of CRISPRi^+^ element-gene pairs; Middle: Density of CRISPRi^-^ element-gene pairs; Right: Density difference between CRISPRi^+^ and CRISPRi^-^ element-gene pairs. The dashed line represents the linear regression line. **e.** We fit a linear regression between scaled ln(ABC_score) and scaled Kendall correlations, and define the integrated Activity, Responsiveness and Contact(ARC)-E2G values along the linear regression line (see **Methods**). Projected ARC-E2G score ranks to the same plot as in (a), with a color gradient from blue to red representing low to high ARC-E2G scores.

**Fig. S2.**
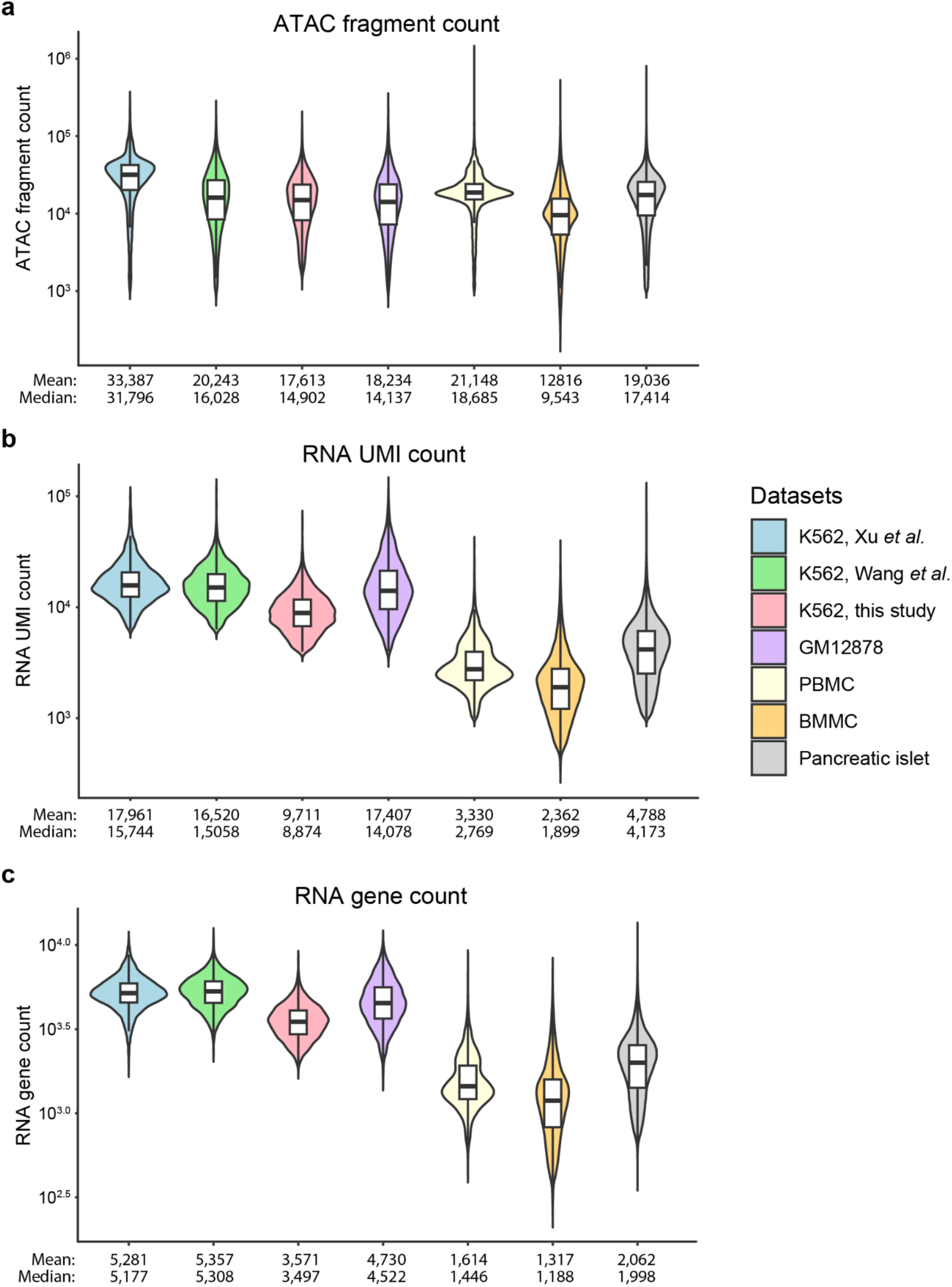
Data quality of single-cell multiome datasets. **a-c**. Violin plots showing the distribution of ATAC fragment counts per cell (**a**), RNA UMI counts per cell (**b**), and RNA gene counts per cell (**c**) across seven single-cell 10x Multiome datasets used in this study. Each violin density contains a box that indicates the median (thick horizontal line), and the first and third quartiles (lower and upper boundaries of the box) of the data. Both mean and median values are labeled below the corresponding violin density.

**Fig. S3.**
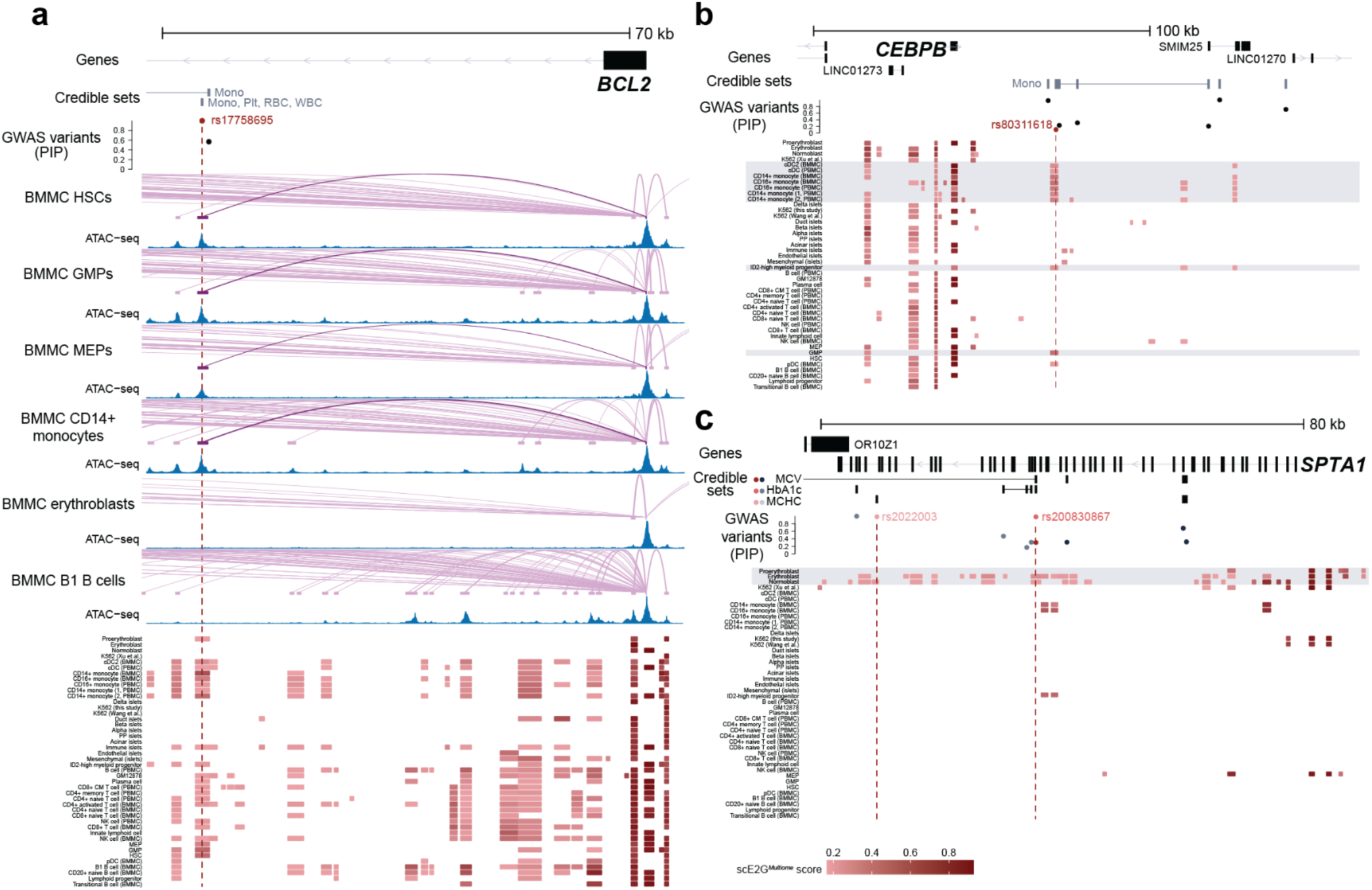
Cell-type specific enhancer-gene regulatory predictions by scE2G*^Multiome^*. **a.** scE2G*^Multiome^* predictions for *BCL2* across 44 cell types for the region chr18:63,245,257-63,325,893 (hg38). Noncoding fine-mapped GWAS variants with PIP > 10% and credible sets for monocyte, platelet, red blood cell, and white blood cell count in the region are shown. Indicated in red is the variant rs17758695, which has a PIP > 99% for each of these traits and has been previously reported to be active in several progenitor cell types and monocytes^71^. Enhancer-gene linking arcs and normalized ATAC signals are shown for several cell types; predictions overlapping rs17758695 are shown in darker purple. The heatmap shows elements linked to *BCL2* with a scE2G*^Multiome^* score above the threshold in each cell type, with cell types ordered by similarity in RNA expression profile. The color of the enhancer indicates the magnitude of the scE2G score for *BCL2*. **b.** Heatmap, as described in **a**, showing elements linked to *CEBPB* by scE2G*^Multiome^* with a score above the threshold for the region chr20:50,145,000-50,310,000 (hg38), the locus shown in **Fig. 1c**. Noncoding fine-mapped GWAS variants with PIP > 10% and credible sets for monocyte count are also shown. The variant rs80311618, which overlaps a myeloid-specific enhancer, is indicated in red. Highlighted rows in the heatmap indicate myeloid cell types in which the variant may be expected to be active. **c.** Heatmap, as described in **a**, showing elements linked to *SPTA1* by scE2G*^Multiome^* with a score above the threshold for the region chr1:158,605,000-158,702,000 (hg38), the locus shown in **Fig. 1d**. Noncoding fine-mapped GWAS variants with PIP > 10% and credible sets for hemoglobin A1C (HbA1c), mean corpuscular volume (MCV), and mean corpuscular hemoglobin concentration (MCHC) are shown. The variants rs2022003, in a credible set for MCHC, and rs200830867, in credible sets for HbA1c and MCV, are highlighted in red. Highlighted rows in the heatmap indicate erythroid cell types in which the variants may be expected to be active.

**Fig. S4.**
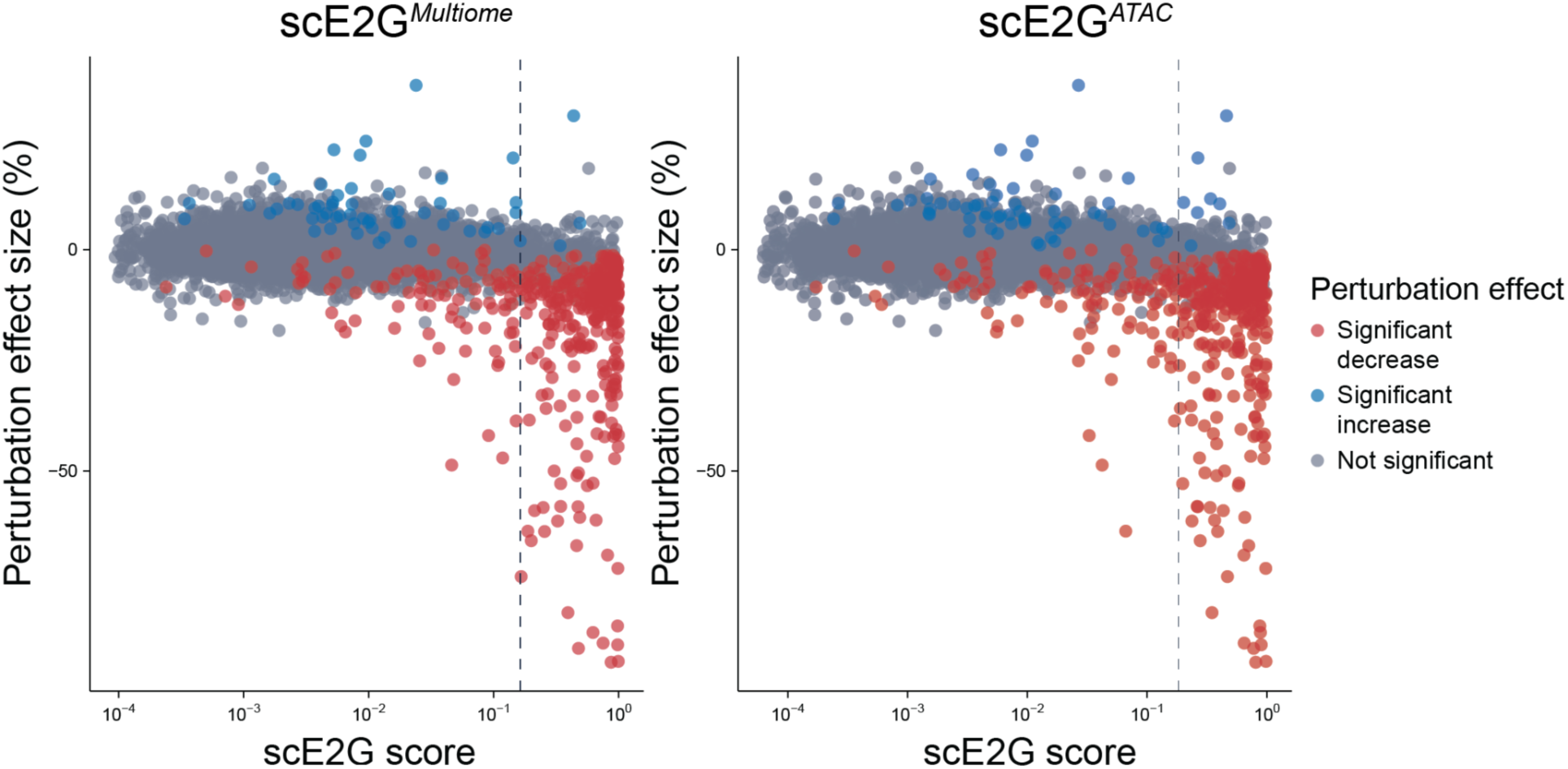
scE2G scores versus CRISPRi effect size. CRISPRi effect sizes on gene expression versus scE2G*^Multiome^* (**left**) and scE2G*^ATAC^* (**right**) scores. Each point represents one tested element-gene pair. CRISPRi effect sizes represent the % change in gene expression relative to unperturbed cells. Model threshold is indicated with a vertical dashed line. scE2G scores were computed from the K562 10x Multiome data from Xu *et al.* using models trained on hold-one-chromosome-out data. To annotate CRISPR-tested element-gene pairs with scE2G scores, we used the same approach as in the CRISPR benchmarking pipeline: the maximum score of overlapping scE2G element-gene pairs was assigned.

**Fig. S5.**
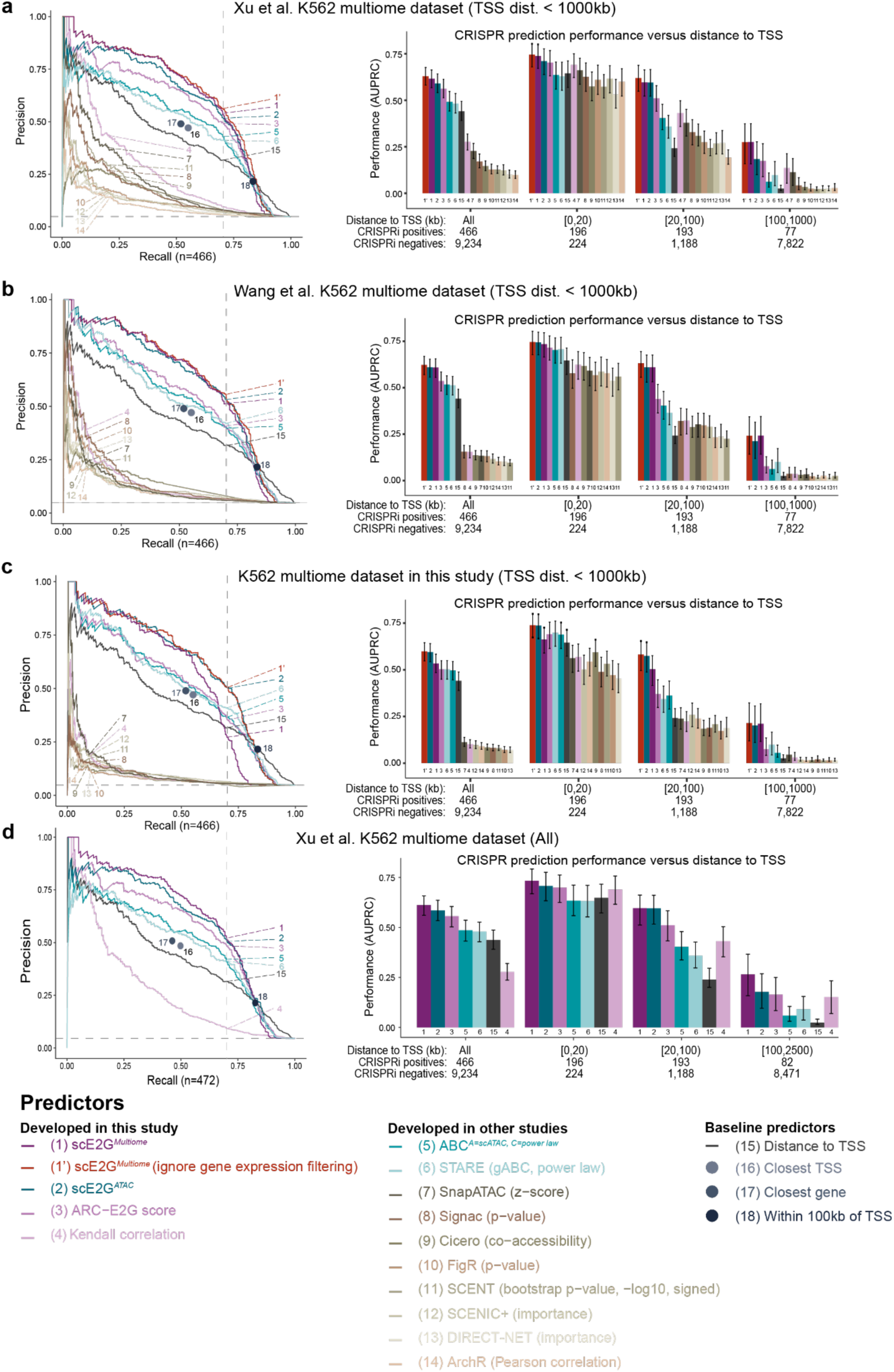
CRISPR benchmarking on three different K562 multiome datasets. **a-c**. CRISPR benchmarking (TSS distance < 1Mb) of scE2G models on K562 single-cell datasets from Xu et al.^40^ (**a**), Wang et al.^72^ (**b**) and this study (**c**). **Left**: Precision–recall curves displaying the performance of models on K562 CRISPRi data, as in **Fig. 2b**. The benchmarking retained only enhancer-gene pairs with a TSS distance of less than 1Mb (9,700 tested element-gene pairs, 466 positives). Each curve represents a continuous predictor. Each single dot indicates the performance of a binary predictor. Dashed vertical line marks performance at a threshold corresponding to 70% recall; dashed horizontal line indicates the proportion of CRISPRi^+^ E-G pairs among all CRISPRi validated E-G pairs (precision value at 100% recall). **Right**: CRISPRi benchmarking performance (AUPRC) of continuous predictors varies with distance to TSS. Error bars show the 95% confidence interval of AUPRC values inferred via bootstrap (1,000 iterations). In the default scE2G*^Multiome^* results (1), we assign predictions only to genes that are sufficiently highly expressed in a given cell type (pseudo-bulk TPM > 1, see **Methods**). Here, we also compared scE2G*^Multiome^* results without gene expression filtering (1’). We note that gene expression filtering primarily impacts the scE2G*^Multiome^* performance on the K562 dataset in panel (**c**), indicating worse gene expression quantification than in the K562 single-cell datasets from Xu *et al.*^40^ (**a**) and Wang *et al.*^72^ (**b**) (See also **Fig. S2b,c**). **d.** Similar to **a**, but comparing to all CRISPR-tested element-gene pairs in the dataset (10,375 tested element-gene pairs, 472 positives), including those located at distances > 1 Mb.

**Fig. S6.**
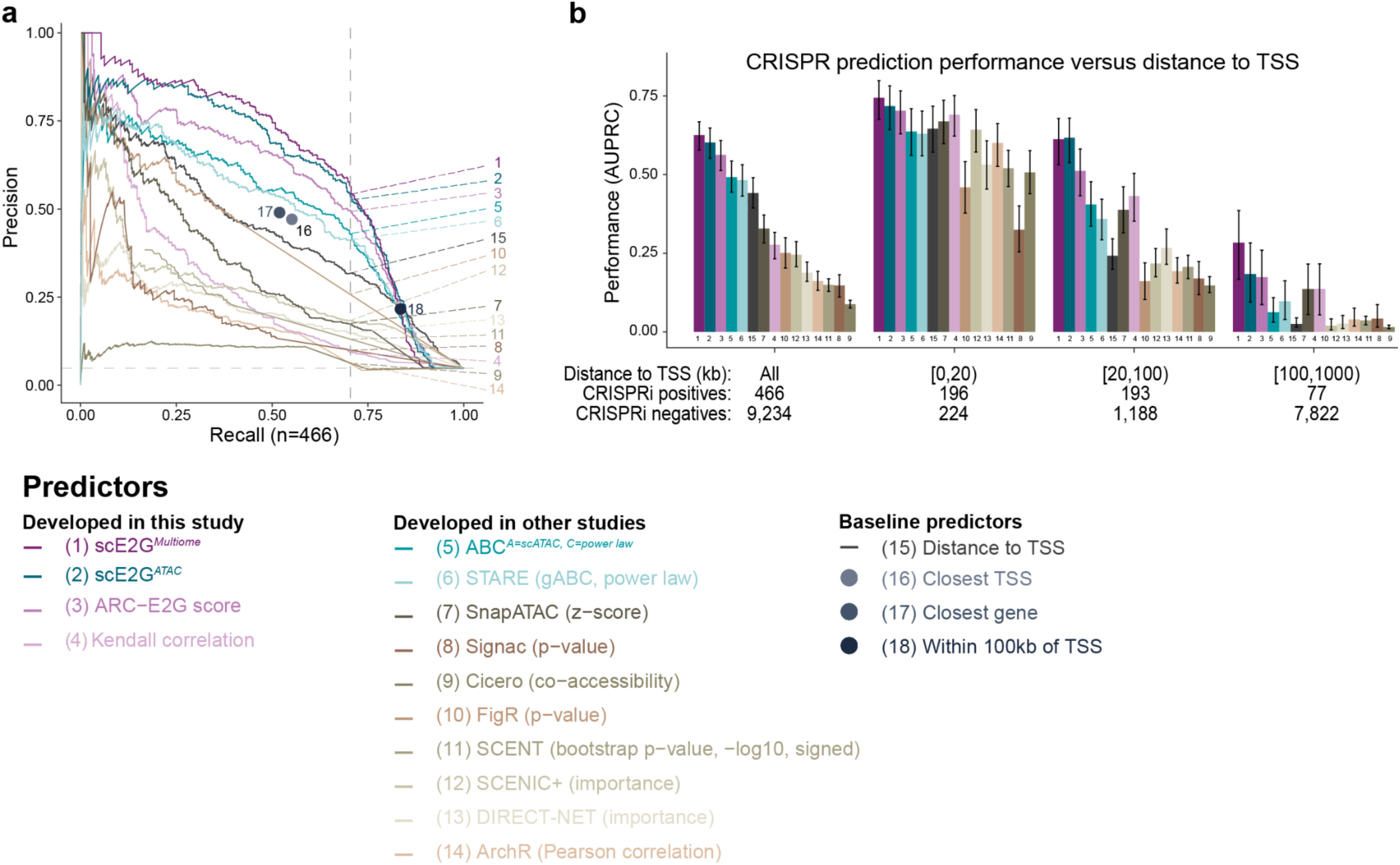
CRISPR benchmarking of existing models using their default settings. **a.** Precision–recall curves displaying the performance of models on K562 CRISPRi data, which were re-analyzed and assembled in Gschwind *et al.*, 2023^20^ using datasets from Nasser *et al.*, 2021^48^, Gasperini *et al.*, 2019^38^ and Schraivogel *et al.*, 2020^39^. The benchmarking retained only enhancer-gene pairs with a TSS distance of less than 1Mb (9,700 tested element-gene pairs, 466 positives). Each curve represents a continuous predictor. Each single dot indicates the performance of a binary predictor. Dashed vertical line marks performance at a threshold corresponding to 70% recall. This figure uses the published recommended thresholds (*e.g.*, distance thresholds) for each method. In contrast, **Fig. 2b** uses a consistent set of thresholds across methods. **b.** CRISPRi benchmarking performance (AUPRC) of continuous predictors varies with distance to TSS. Error bars show the 95% confidence interval of AUPRC values inferred via bootstrap (1,000 iterations).

**Fig. S7.**
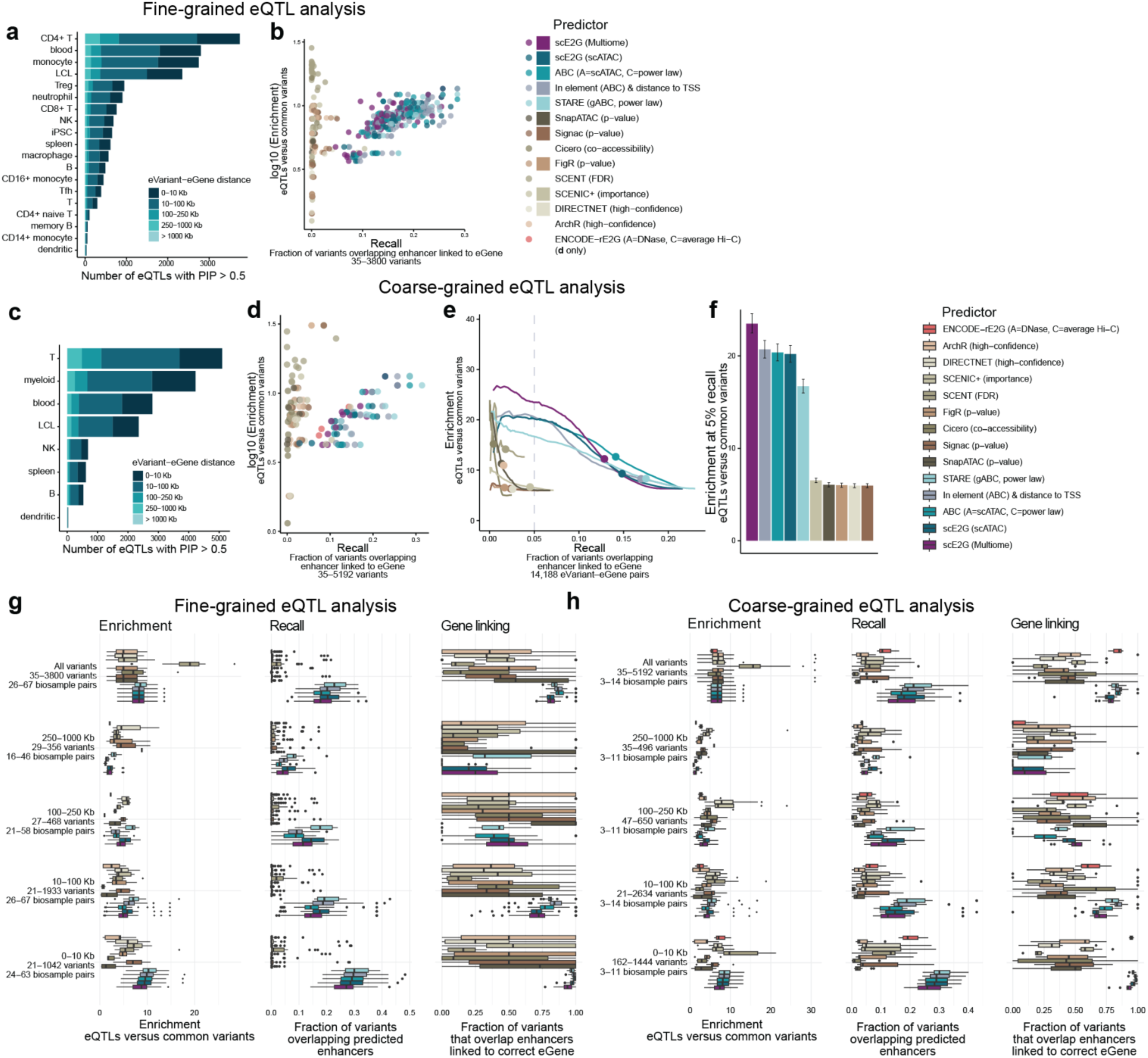
Benchmarking scE2G using fine-mapped eQTLs. **a.** Number of fine-mapped eQTL variants (eVariants), stratified by eVariant–eGene distance, for each eQTL biosample considered in the “fine-grained” eQTL analysis shown in **Fig. 2d,e** (*e.g.*, eQTLs in CD4+ naive T cells matched to model predictions from T cell types, **Table S4**). **b.** Enrichment and recall for predicted enhancers containing fine-mapped eVariants from the “fine-grained” eQTL analysis. Each dot represents one matched eQTL-model biosample. Note that the number of matched biosamples varies per model (see **Table S4**). **c.** Number of eVariants, stratified by eVariant–eGene distance, for each eQTL biosample considered in a “coarse-grained” eQTL analysis (*e.g.*, eQTLs from T cells matched to model predictions from T cell type “supergroup”, **Table S4**). The “coarse-grained” eQTL analysis is shown to compare all single-cell models on the same set of matched eQTL-model biosamples. **d.** Enrichment and recall for predicted enhancers containing fine-mapped eVariants from the “coarse-grained” eQTL analysis. Each dot represents one matched eQTL-model biosample. There are 3 matched biosamples for ENCODE-rE2G and 13 matched biosamples for all other models (see **Table S4**). **e.** Enrichment–recall curves showing performance of models from the “coarse-grained” eQTL analysis as described in **d**. Enrichment and recall were aggregated across matching biosamples by summing variant counts across all biosample pairs and were calculated for ∼100 threshold values for continuous predictors. Points indicate enrichment and recall at suggested thresholds for each model. The dashed vertical line indicates a recall of 5%. Error bars at the point represent 95% confidence intervals calculated from the natural log of enrichment. **f.** Enrichment of variants at 5% recall aggregated across all biosample pairs in the “coarse-grained” eQTL analysis in **e**. Error bars represent 95% confidence intervals calculated from the natural log of enrichment. **g.** Boxplots showing enrichment, recall (fraction of variants overlapping any enhancer, not accounting for eGene linking), and fraction of variants overlapping an enhancer linked to the correct eGene given that they overlap an enhancer, for the “fine-grained” eQTL analysis. Results are stratified by distance between eVariant and eGene. See Methods, ‘Data visualization’ section for definition of box plot elements. **h.** Same as **g**, but for the “coarse-grained” eQTL analysis.

**Fig. S8.**
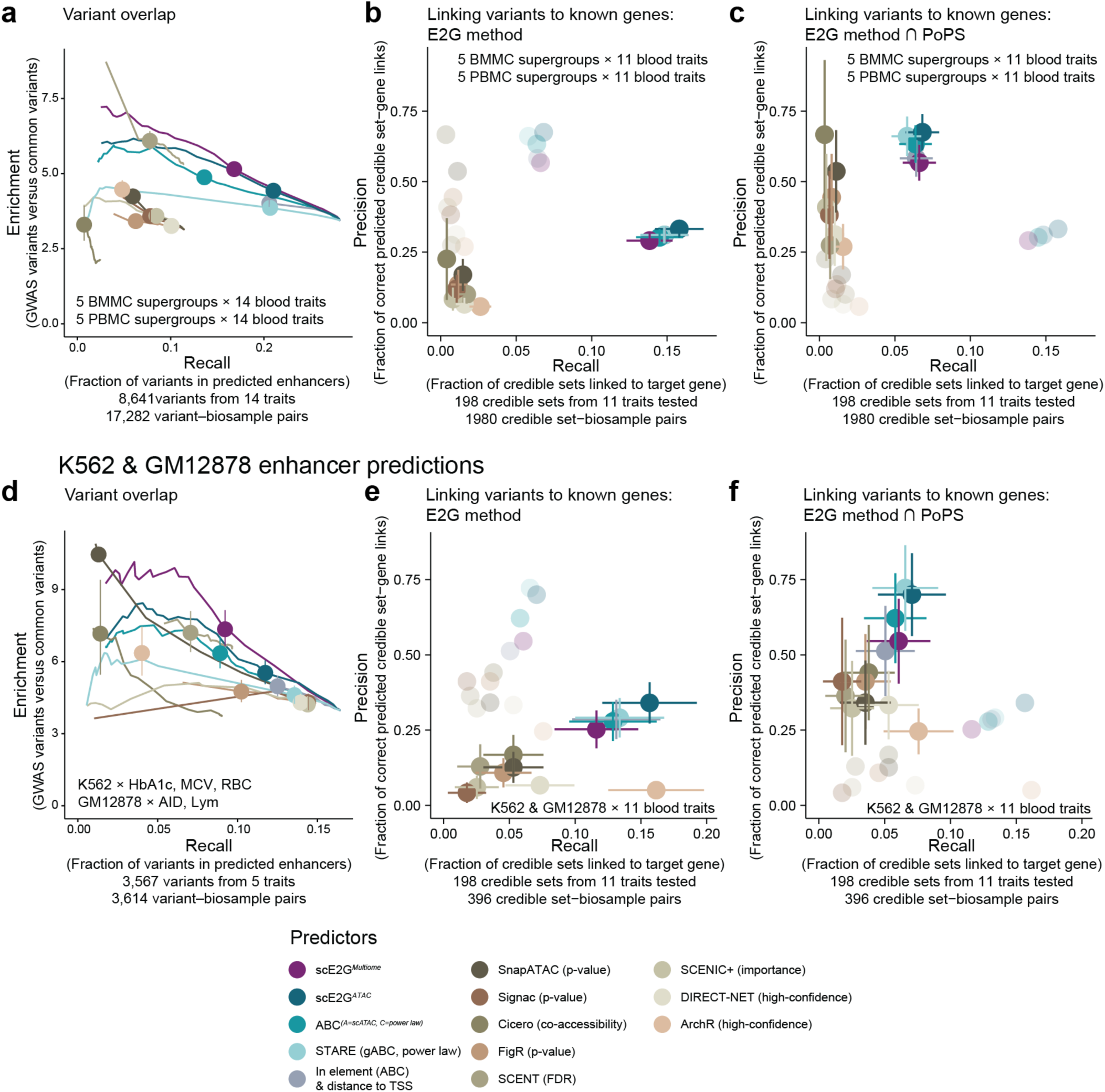
Benchmarking scE2G using fine-mapped GWAS variants. **a.** Enrichment–recall curves showing overlap of fine-mapped GWAS variants for 14 blood-related traits in enhancer predictions for 10 cell type “supergroups” from BMMCs and PBMCs (**Table S5**). Evaluation of predictions in cell type “supergroups” is shown here to compare all single-cell models on the same set of biosamples. Enrichment (y-axis) is calculated as the ratio of GWAS variants in predicted enhancers compared to distal noncoding variants; recall (x-axis) is calculated as the fraction of variants overlapping enhancers. Performance was evaluated at ∼100 score thresholds for continuous predictors. Points indicate enrichment and recall at suggested thresholds for each model, and error bars represent 95% confidence intervals calculated from the natural log of enrichment. **b.** Precision and recall of models at linking credible sets to causal genes (inferred from independent signals with coding variants) for enhancer predictions from the same 10 cell type supergroups as **a**, and credible sets from 11 blood-related traits (MCH, MCHC, MCV, Hb, HbA1c, RBC, Mono, Lym, Eosino, Neutro, Plt). Predictions are binarized with each model’s suggested threshold. The top two target genes based on model score for each credible set were considered positive predictions. Error bars represent 95% confidence intervals based on the Wilson score interval. Faded points and underlined numbers indicate performance of the models when predictions are intersected with PoPS, as in **c**. **c.** Same as **b**, except enhancer predictions were intersected with the top two genes linked to the credible set by PoPS. Faded points and underlined numbers indicate performance of the models when predictions before intersecting with PoPS, as in **c**. **d.** Enrichment–recall curves as in **a** showing overlap of fine-mapped GWAS variants for hemoglobin A1c (HbA1c), mean corpuscular volume (MCV), and red blood cell count (RBC) in K562 predictions and variants for autoimmune disorders (AIDs) and lymphocyte count (Lym) in GM12878 predictions. **e.** Precision and recall of models at linking credible sets to known causal genes for binarized enhancer predictions, as in **b**. Enhancer predictions for K562 and GM12878 were combined and evaluated against credible sets from 11 blood-related traits (**Table S5**). Faded points indicate performance of the models when predictions are intersected with PoPS, as in **g**. Evaluation of predictions in K562 and GM12878 cells is shown as a direct comparison to our previous evaluation of bulk predictors^20^. **f.** Same as **e**, except enhancer predictions were intersected with the top two genes linked to the credible set by PoPS. Faded points indicate performance of the models when predictions before intersecting with PoPS, as in **e**.

**Fig. S9.**
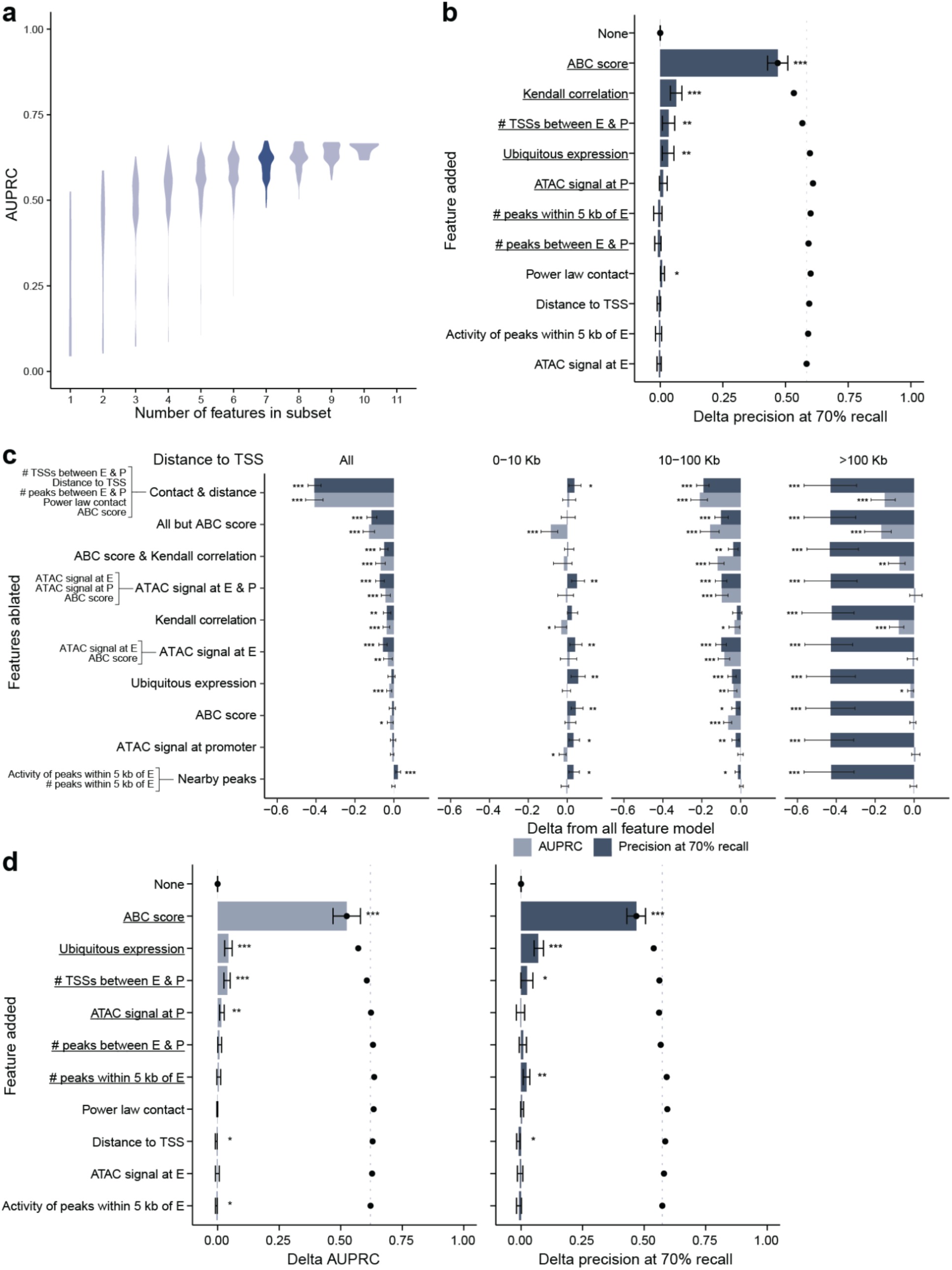
Feature interpretation of scE2G*^Multiome^* and scE2G*^ATAC^*. **a.** Violin plots showing AUPRC on the CRISPR benchmark for all possible subsets of the 11 starting features (ABC score and Kendall correlation as separate features) for each subset size. The distribution of AUPRC for 7-feature subsets, which contains the features used in the final scE2G*^Multiome^* model at which the maximum AUPRC is achieved, is highlighted. **b.** Forward sequential feature selection for scE2G*^Multiome^* (ABC score and Kendall correlation as separate features), as shown in **Fig. 3a**, but bars represent the change in precision at 70% recall (instead of AUPRC) from the previous model including features above it. The order of added features was determined by maximum change in AUPRC. Dots indicate the accumulated change in precision for the model including the feature labeled and all above it, and the dashed vertical line represents the total accumulated precision across all features. Error bars represent 95% range of values inferred via bootstrap (1,000 iterations). One, two, and three stars indicate p-values less than 0.05, 0.01, and 0.001, respectively; black stars represent a positive change and red stars represent a negative change. The underlines indicate the features in the final model. **c.** Category feature ablation for scE2G*^Multiome^* (ABC score and Kendall correlation as separate features) showing change in AUPRC and precision at 70% recall on the CRISPR benchmark, with results shown for element-gene pairs stratified by distance to TSS. Error bars and significance indicators are as described in **b**. **d.** Forward sequential feature selection for scE2G*^ATAC^*, with bars representing change in AUPRC (left) and precision at 70% recall (right). The order of added features was determined by maximum change in AUPRC. Error bars, significance indicators, and underlined features are as described in **b**.

**Fig. S10.**
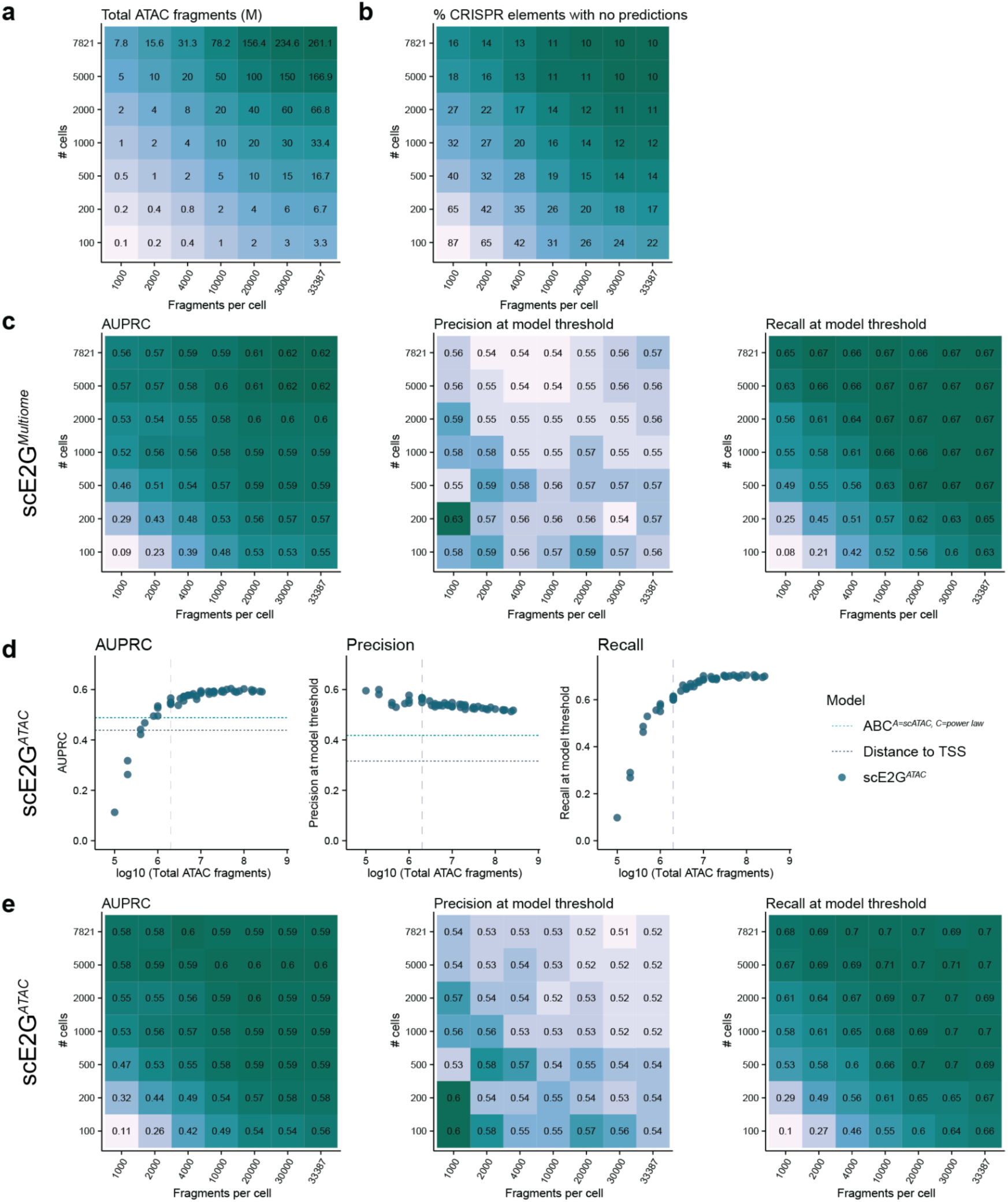
Robustness of scE2G to sequencing depth. **a.** Heatmap of fragment numbers for downsamples of the Xu et al.^40^ K562 10x Multiome dataset. **b.** Heatmap showing percent of elements tested in the CRISPR benchmark (most of which are accessible) that do not overlap candidate elements considered in scE2G predictions for each downsample. Here, overlap is defined as sharing at least 50% of base pairs. **c.** Heatmaps of performance on the CRISPR benchmark for scE2G*^Multiome^* across downsamples. **d.** AUPRC (left), precision at the model threshold (center) and recall at the model threshold (right) of scE2G*^ATAC^* on the CRISPR benchmark across ATAC fragment counts. Performance of distance to TSS (grey line) and ABC on the full-depth dataset (teal line, the next best model after scE2G*^Multiome^* and scE2G*^ATAC^*) are shown for reference. These reference lines are not shown for recall because the score threshold for models is defined based on 70% recall. The vertical dashed lines indicate 2 million fragments (and 1 million RNA UMIs). **e.** Heatmaps of performance on the CRISPR benchmark for scE2G*^ATAC^* across downsamples.

**Fig. S11.**
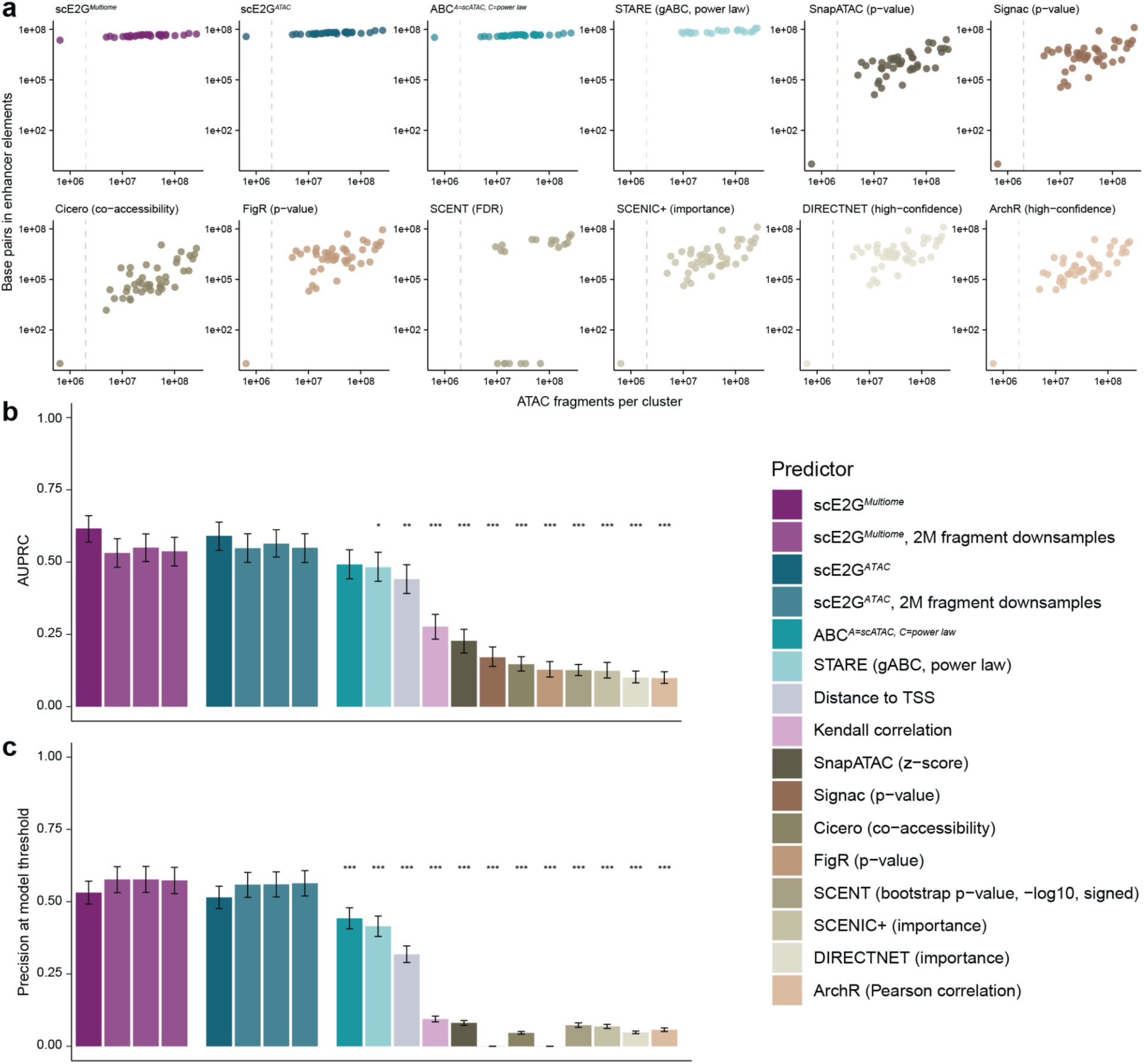
Robustness of scE2G to sequencing depth compared to other models. **a.** Number of basepairs included in predicted enhancers as a function of ATAC fragments per cell type. Predictions are binarized at the thresholds for each model. Each dot is one model biosample, including PBMC and BMMC cell types, cell type supergroups, K562, and GM12878. To obtain model predictions for a given cell type (*e.g.* PBMC B cells) for published predictors, we filtered predictions generated on the full dataset (e.g. all PBMCs) to those with detected peaks and genes (see **Methods**). **b.** AUPRC on the CRISPR benchmark in K562 cells, comparing scE2G predictions made on the full-depth dataset, scE2G predictions made on three downsampled datasets, and other models made on the full-depth dataset. The three downsampled datasets each comprised 2 million ATAC fragments and 1 million RNA UMIs and were made up of different numbers of cells: 1) 500 cells with 4000 ATAC fragments and 2000 RNA UMIs per cell, 2) 1000 cells with 2000 ATAC fragments and 1000 RNA UMIs per cell, 3) 2000 cells with 1000 ATAC fragments and 400 RNA UMIs per cell. Note that some existing models were scored based on different metrics than in **a**. Error bars represent 95% confidence intervals inferred via bootstrapping with 10,000 replicates. Stars signify p-values for the delta performance (also inferred via bootstrapping with 10,000 replicates) between the model with the stars applied to the full-depth dataset and the six predictions from scE2G*^Multiome^* and scE2G*^ATAC^* on each of the three downsamples. Three stars represents all p-values < 0.001, two stars represent all p-values < 0.01, one star represents all p-values < 0.05. **e.** Same as **d**, but for precision at the model threshold.

**Fig. S12.**
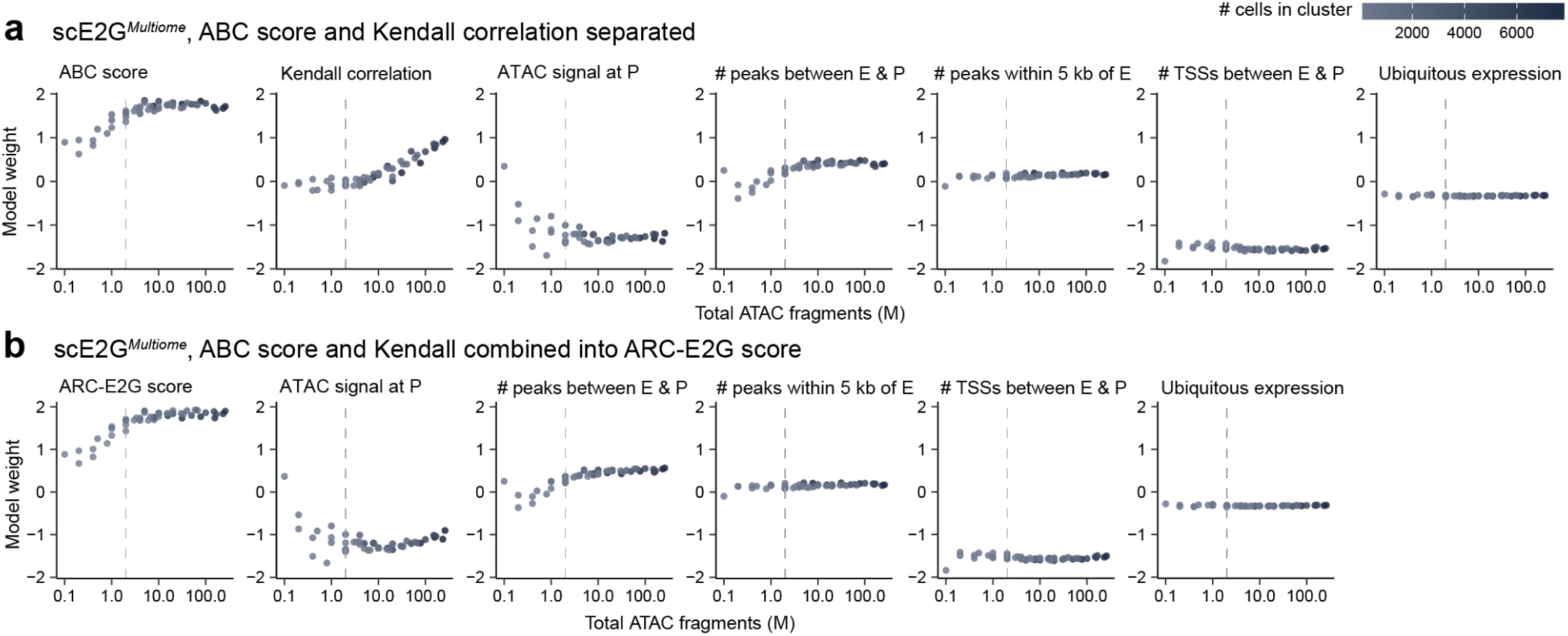
Model weights robustness to sequencing depth. **a.** Model weights logistic regression models trained on each downsample as a function of log10(total ATAC fragments) for a model trained with ABC and the Kendall correlation separately. The vertical dashed line indicates 2 million total ATAC fragments. Note that the model weight for the Kendall correlation increases beyond the 2 million fragment mark. **b.** Model weights logistic regression models trained on each downsample as a function of log10(total ATAC fragments) for a model trained with ARC-E2G integrating ABC and the Kendall correlation. The vertical dashed line indicates 2 million total ATAC fragments. Note that the model weight for ARC-E2G score is stable beyond 2 million fragments.

**Fig. S13.**
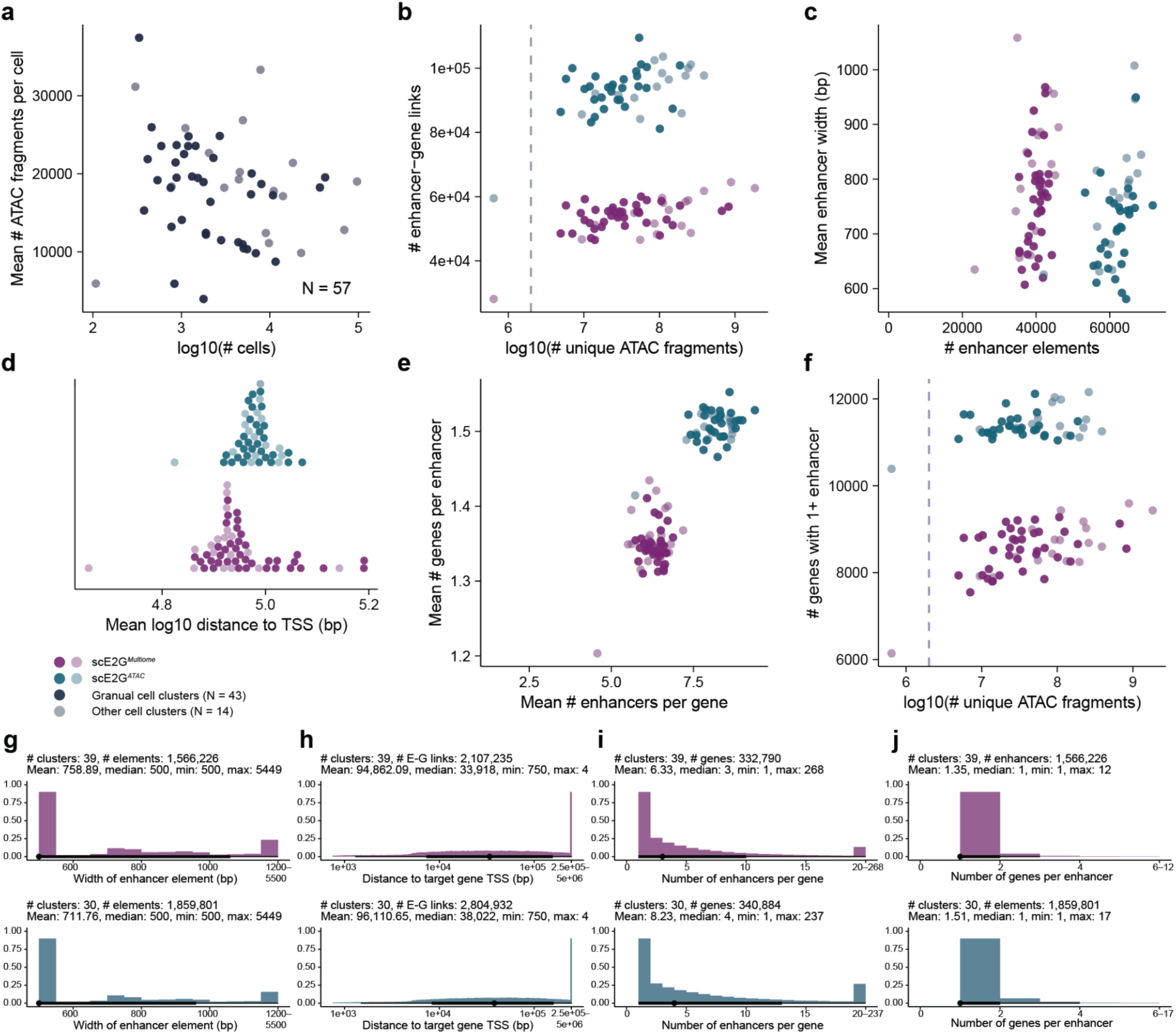
Properties of genome-wide scE2G predictions. **a–f**: Each point represents the average value of a property for enhancer-gene predictions in one cell type or cell type supergroup. Purple points correspond to scE2G*^Multiome^* predictions (N = 57), and teal points correspond to scE2G*^ATAC^* predictions (N = 46). Darker points represent the 39 cell types from the PBMC, BMMC, and pancreatic islet datasets with more than 2 million ATAC fragments. All properties were computed on binarized predictions omitting promoter elements. **a.** Average number of unique ATAC fragments per cell by number of cells per cell type. **b.** Number of enhancer-gene links by number of unique ATAC fragments. The dashed vertical line indicates 2 million ATAC fragments. **c.** Mean enhancer width by number of unique enhancers. **d.** Mean distance between enhancer and target gene transcription start site. **e.** Mean number of enhancers per gene by mean number of genes per enhancer. **f.** Number of genes with at least one non-promoter enhancer by number of unique ATAC fragments. The dashed vertical line indicates 2 million ATAC fragments. **g–j**: Histograms showing the distribution of properties across all predictions in the 39 cell types from the PBMC, BMMC, and pancreatic islet datasets with more than 2 million ATAC fragments (scE2G*^Multiome^*, purple) or 30 cell types from the PBMC and BMMC datasets with more than 2 million ATAC fragments (scE2G*^ATAC^*, teal). As in **g–j**, all properties were computed on binarized predictions omitting promoter elements. See Methods, ‘Data visualization’ section for definition of point interval elements below each histogram.

**Fig. S14.**
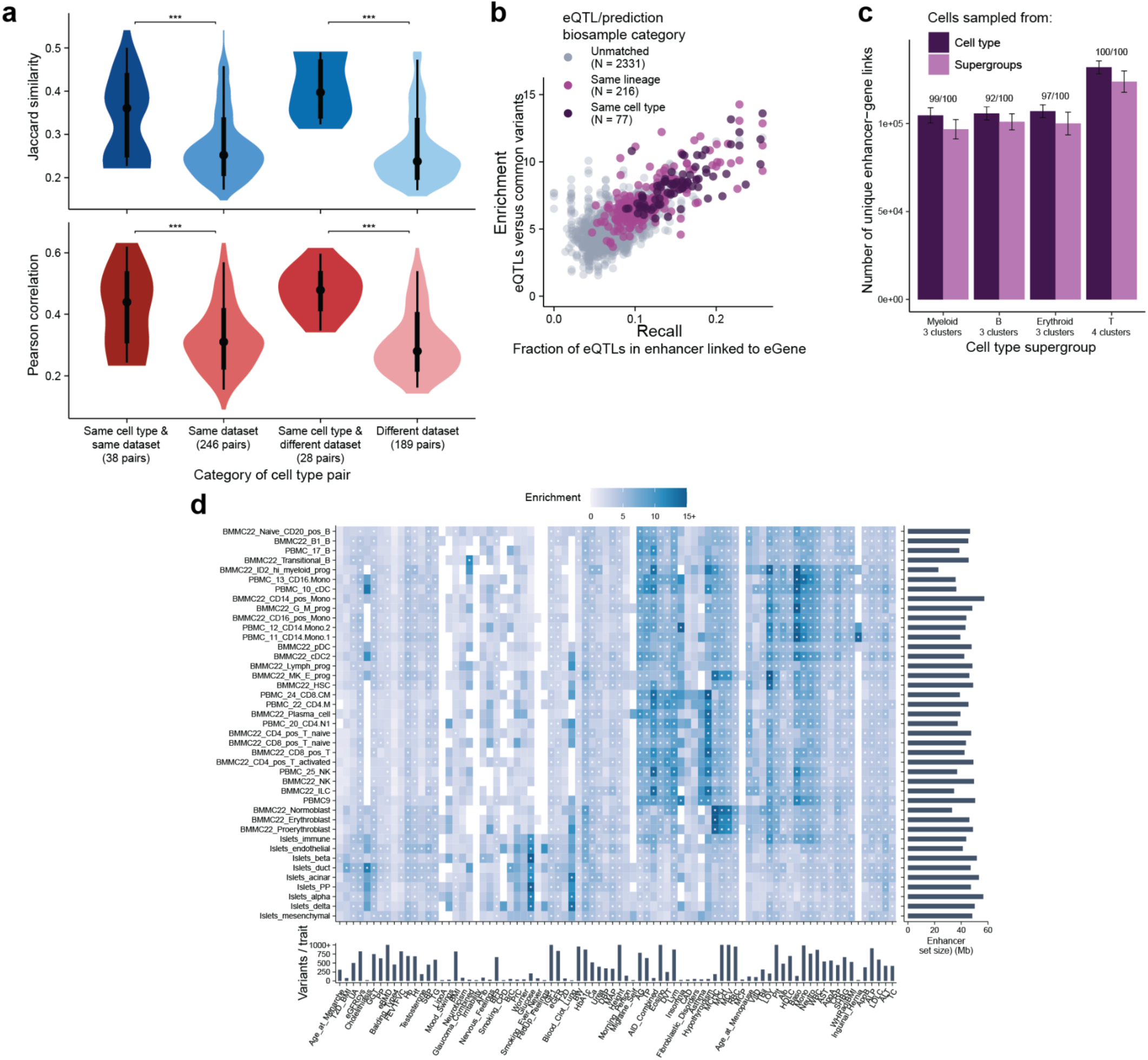
Cell-type resolution of scE2G*^Multiome^* predictions. **a.** Jaccard similarity (top, blue) and Pearson correlation (bottom, red) between scE2G*^Multiome^* predictions for different categories of cell type pairs for 30 PBMC and BMMC cell types: 1) same cell type and same dataset (*e.g.* PBMC CD14+ monocytes and PBMC CD16+ monocytes); 2) same dataset (*e.g.* all pairs of PBMC cell types); 3) same cell type and different dataset (*e.g.* PBMC CD14+ monocytes and BMMC CD16+ monocytes); 4) different dataset (all pairs of one PBMC and one BMMC cell type). Three stars indicate a Bonferroni-corrected p-value less than 0.001 from a two-sided t-test between distributions. See Methods, ‘Data visualization’ section for definition of point interval plot elements. **b.** Enrichment and recall of eVariants in scE2G*^Multiome^* predictions from biosamples of the same cell type (dark purple), same cell lineage (light purple), or unmatched biosamples (grey). Each point represents one eQTL, prediction biosample pair. Enrichment was computed as the fraction of fine-mapped distal noncoding eQTLs (PIP > 50%) in predicted enhancers compared to distal noncoding common variants. Recall was computed as the fraction of fine-mapped distal noncoding eVariant–eGene links (PIP > 50%) for which the eVariant is located in a predicted enhancer linked to the correct eGene. **c.** Comparison of number of unique predicted enhancer-gene links by scE2G*^Multiome^* in BMMC cell type supergroups for either random, non-overlapping samples of cells selected from fine-grained cell types (dark purple) or from the same number of non-overlapping samples selected from the cell type supergroup (see Methods). After dividing all cells in the dataset with both sampling methods, unique combinations of samples were selected and compared for each supergroup 100 times. Error bars represent 95% confidence intervals for the total number of enhancer-gene predictions across the 100 iterations. Fractions above the bars indicate the number of iterations in which scE2G*^Multiome^* predictions from cells sampled from cell types had more enhancer-gene links than from cells sampled from the supergroup. **d.** Enrichment of fine-mapped GWAS variants from 86 traits versus common variants in scE2G*^Multiome^* predictions for all cell types. Horizontal margin plot shows the number of variants per trait, and the y-axis is capped at 1000. The vertical margin plot shows the number of base pairs predicted to be in enhancers. The color scale for enrichment is capped at 15. Stars indicate significant enrichment (padjusted < 0.05). Both axes are hierarchically-clustered with the Ward2 method based on Euclidean distance of the correlation across the enrichment matrix.

**Fig. S15.**
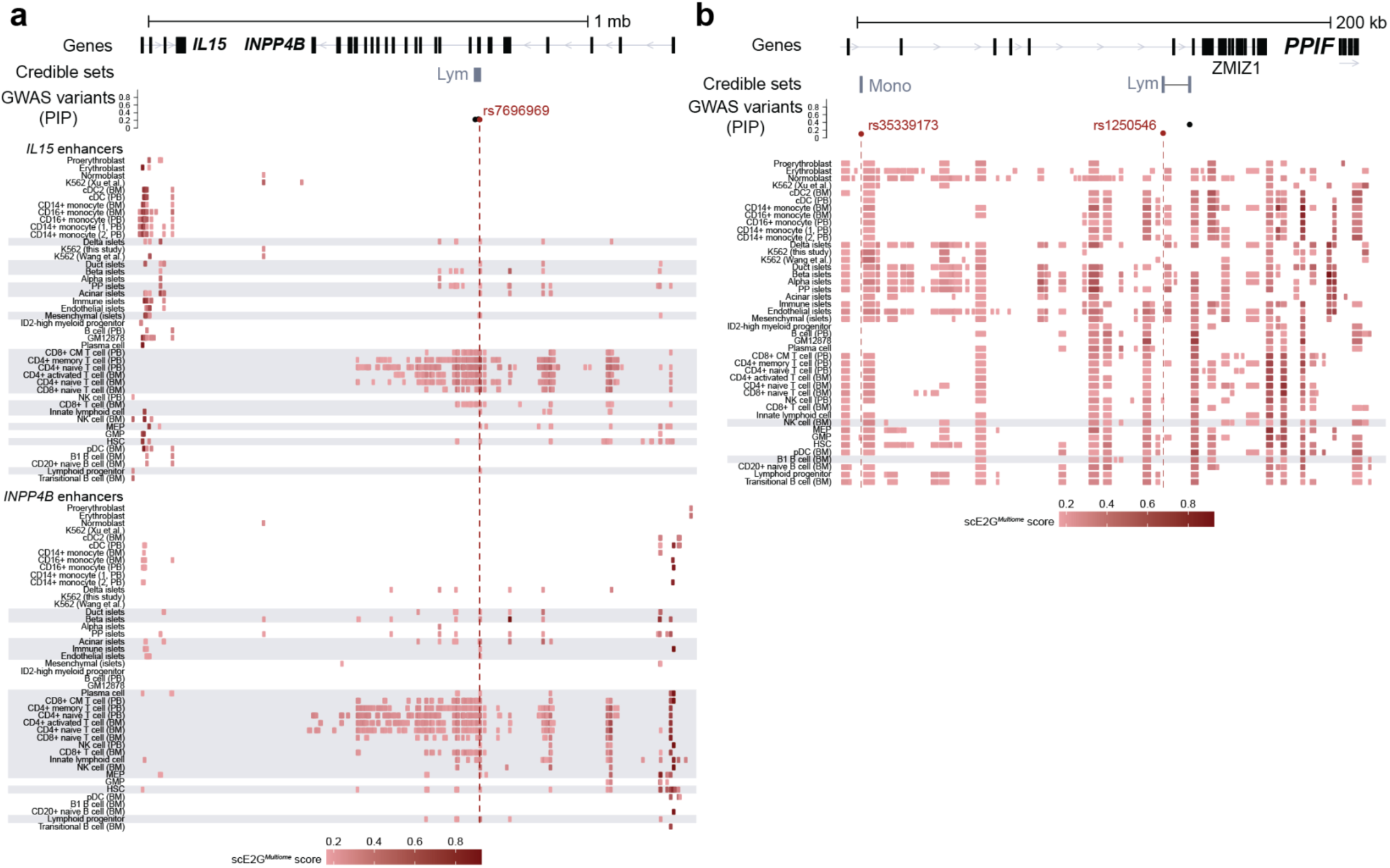
GWAS variant interpretation: enhancer predictions across cell types for selected loci. **a.** Heatmaps showing elements linked to *IL15* (top) and *INPP4B* (bottom) by scE2G*^Multiome^* with a score above the threshold across 44 cell types for the region chr4:141,612,977-142,900,000 (hg38), the locus shown in **Fig. 5c**. Cell types are ordered by similarity in RNA expression profile. The color of the enhancer indicates the magnitude of the scE2G score. Fine-mapped GWAS variants with PIP > 10% and credible sets for lymphocyte count are also shown. The variant rs7696969 is indicated in red. **b.** Heatmap, as described in **a**, showing elements linked to *PPIF* by scE2G*^Multiome^* with a score above the threshold across 44 cell cell types for the region chr10:79,136,379-79,375,225 (hg38), the locus shown in **Fig. 5f**. Fine-mapped GWAS variants with PIP > 10% and credible sets for monocyte count and lymphocyte count are also shown. The variants rs35339173 (monocyte count) and rs1250546 (lymphocyte count) are indicated in red.

## Supplementary Notes

### Note S1. Summary of published predictors

1. *Signac*^34^: Signac computes the Pearson correlation between gene expression and enhancer accessibility for each enhancer-gene pair. For each enhancer, SnapATAC selects 200 peaks with similar GC content, total accessibility level, and sequence length from different chromosomes as the background set. Then, Signac calculates the Pearson correlations between the target gene and these background peaks. A z-score is computed for each enhancer-gene pair using the formula z = (r-μ)/δ, where μ is the mean background Pearson correlation and δ is the standard deviation of the background Pearson correlations for the peak. Signac then computes a p-value for each peak using a one-sided z-test.
2. *FigR*^31^: FigR computes the Spearman correlation between gene expression and enhancer accessibility for each enhancer-gene pair. For each enhancer, FigR selects 100 peaks with similar GC content and total accessibility level as the background set. FigR then calculates the Spearman correlations between the target gene and these background peaks. A z-score is computed for each peak using the formula z = (r-μ)/δ, where μ is the mean background Spearman correlation and δ is the standard deviation of the background Spearman correlations for the peak. FigR computes a p-value for each peak using a one-sided z-test.
3. *SnapATAC*^35^: SnapATAC fits a logistic regression model for each pair of binarized enhancer accessibility and gene expression.
4. *SCENT*^33^: SCENT fits a Poisson regression between gene expression count and variables including enhancer accessibility, total UMI count, batch, and %mitochondrial UMI to identify the association between gene expression and enhancer accessibility, while controlling for interference from other factors. To estimate the error and significance, SCENT applies a bootstrapping approach to calculate the False Discovery Rate (FDR).
5. *SCENIC*+^32^: SCENIC+ uses GRNBoost2^73^, which performs gradient-boosting machine regression, to estimate the importance of candidate enhancers for target gene expression. Then it categorizes the results into positive (Spearman correlation > 0.03) and negative (Spearman correlation < −0.03) interactions.
6. *Cicero*^29^: Cicero aggregates single cells into meta-cells, each composed of 50 cells similarly positioned in trajectory. It computes the correlation between promoters and enhancers across meta-cells within 500kb, and penalizes long-distance correlations using Graphical LASSO^74^.
7. *DIRECT-NET*^30^: DIRECT-NET also aggregates single cells into meta-cells, then employs the XGBoost gradient boosting model to regress gene expression against all candidate enhancers across meta-cells. DIRECT-NET identifies functional enhancers based on the importance scores learned from the XGBoost model.
8. *ArchR*^28^: ArchR adopts the same meta-cell aggregation approach as Cicero, then calculates the Pearson correlation between log2-normalized gene expression and peak accessibility across meta-cells.
9. *ABC*: The ABC (activity-by-contact) model estimates the effect of a specific enhancer by multiplying the enhancer activity by the 3D contact frequency, and then dividing this product by the sum of the products of all candidate enhancer activities and their respective contact frequencies to calculate a relative contribution. In this study, we used pseudo-bulk ATAC intensity to represent enhancer activity, and inferred the 3D contact based on the power-law distribution.
10. *STARE (gABC)*^41^: STARE adopts the ABC model and extends it to the generalized ABC (gABC) model, which accounts for not only all candidate enhancers of the target gene but also all target genes of the target enhancer. In the gABC model, enhancer activity is multiplied by the contact frequency, similar to the ABC model. However, it differs in that the result is divided by the sum of the products of enhancer activity and contact frequency for all candidate enhancers and candidate genes.

### Note S2. Benchmarks against bulk DNase-seq ENCODE-rE2G models

We evaluated scE2G models versus their ENCODE-rE2G counterparts, based on bulk DNase-seq, in the CRISPR, eQTL and GWAS benchmarks. First, we compared the performance of K562 predictions on the CRISPR benchmark (**Fig. N1a,b**). Considering all tested element-gene pairs, ENCODE-rE2G achieves a significantly higher AUPRC than both scE2G*^Multiome^* (delta AUPRC = 0.040, p_bootstrap_ = 0.029) and scE2G*^ATAC^* (delta AUPRC = 0.065, p_bootstrap_ = 0.0001). Suggesting that the activity signal itself is responsible for at least some of the discrepancy, ABC*^A=DNase^* achieves a significantly higher AUPRC than ABC*^A=scATAC^* with an even large delta of 0.080 (p_bootstrap_ = 0.0001). We do see that scE2G*^Multiome^*, but not scE2G*^ATAC^*, performs significantly better than ABC*^A=DNase^* (delta AUPRC = 0.048, p_bootstrap_ = 0.011); both scE2G models have higher precision than ABC*^A=DNase^* (delta precision for scE2G*^Multiome^* = 0.050, p_bootstrap_ = 0.0012; delta precision for scE2G*^ATAC^* = 0.033, p_bootstrap_ = 0.027). Considering element-gene pairs with a distance to TSS greater than 100 kb, scE2G models improve their performance relative to ENCODE-rE2G: the differences in AUPRC and precision between both scE2G models and ENCODE-rE2G is not significant; and the difference in AUPRC between scE2G*^Multiome^* and ENCODE-rE2G*^Extended^* is not significant (**Fig. N1b**).

We next evaluated predictions on GM12878 cells against eQTL variants from lymphoblastoid cell lines (LCLs). When predictions are binarized at the designated thresholds, scE2GM has slightly lower recall than ENCODE-rE2G, but they achieve similar enrichment. scE2GATAC performs worse than ENCODE-rE2G on both metrics (**Fig. N1c**). However, evaluating enrichment of variants at a constant recall of 10%, there is not a significant difference in performance between any of scE2G*^Multiome^*, scE2G*^ATAC^*, ABC*^A=scATAC^*, ENCODE-rE2G*^Extended^*, ENCODE-rE2G, or ABC*^A=DNase^* (**Fig. N1d**).

Finally, we evaluated model predictions from K562 cells and GM12878 lymphoblastoid cells against relevant traits (red blood cell count), mean corpuscular volume, and hemoglobin A1c for K562; lymphocyte count and combined autoimmune disorders for GM12878), which allows for a comparison with bulk models. We observe that scE2G*^Multiome^* performs better than all other models, including the best-performing bulk models, at enrichment and recall of variants (**Fig. N1e**), and comparably to bulk models at linking credible sets to their causal gene (**Fig. N1f,g**).

**Fig. N1.**
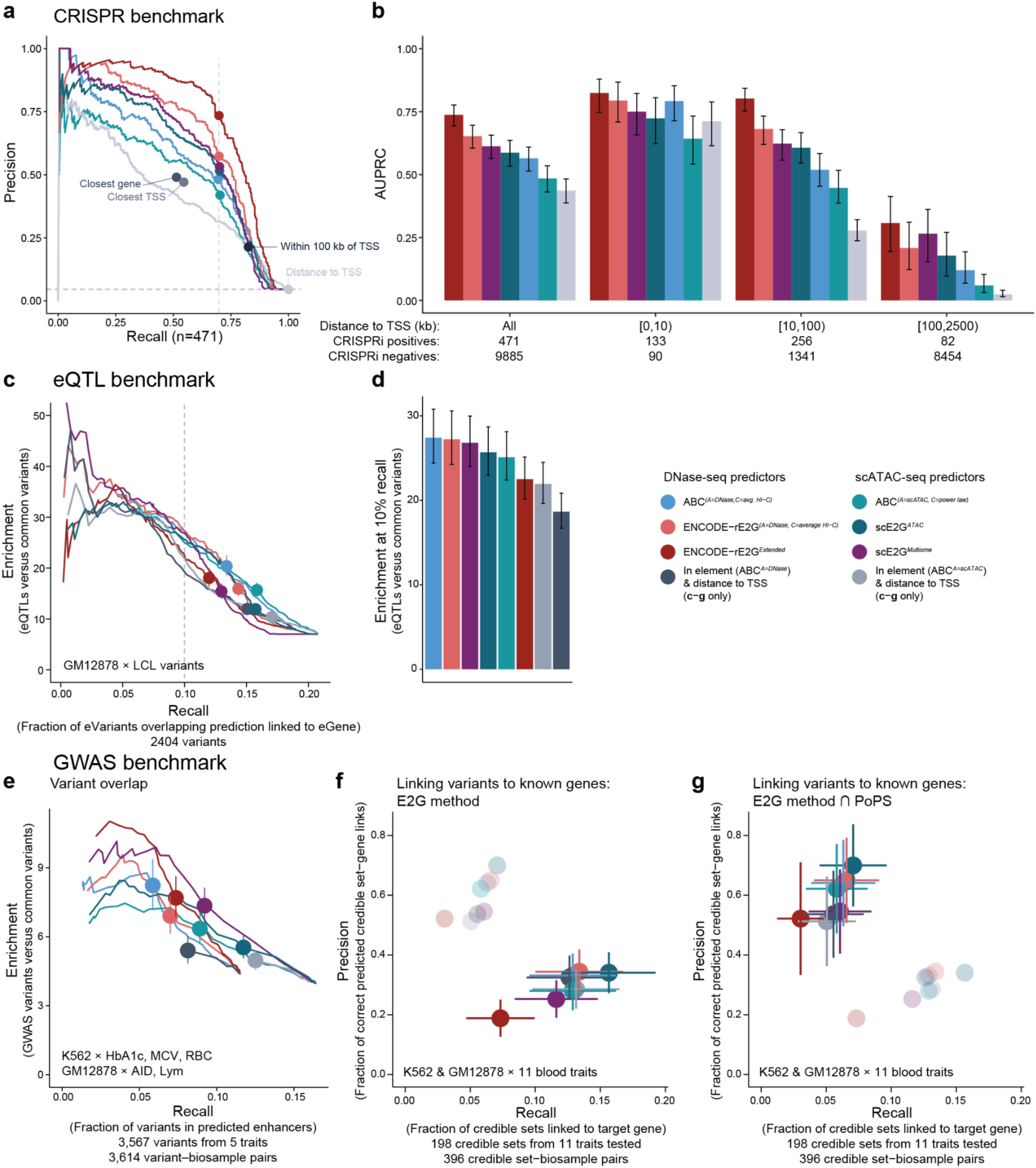
Benchmarking scE2G versus bulk models. **a.** Precision–recall curves display the performance of models on K562 CRISPRi data as in **Fig. 1**. The benchmarking retained all element-gene pairs, including 5 positives and 632 negatives with a TSS distance greater than 1 Mb that were not included in **Fig. 1**. Each curve represents a continuous predictor, and the dot on the curve represents performance at the designated model threshold. Each single dot indicates the performance of a binary predictor. Dashed vertical line marks performance at a threshold corresponding to 70% recall. Dashed horizontal line indicates the proportion of positives. **b.** CRISPRi benchmarking performance (AUPRC) of continuous predictors for element-gene pairs stratified by distance to TSS. Error bars show the 95% confidence interval of AUPRC values inferred via bootstrap (1000 iterations). **c.** Enrichment–recall curves showing performance of models on the eQTL benchmark for enhancer predictions in GM12878 cells and eQTLs from lymphoblastoid cell lines (LCLs, 2,404 variants). Enrichment and recall were calculated for ∼100 threshold values for continuous predictors. Enrichment (y-axis) is the ratio of fine-mapped distal noncoding eQTLs (PIP > 0.5) in predicted enhancers compared to distal noncoding common variants. Recall (x-axis) is the fraction of variants overlapping enhancers linked to the correct gene across the range of score thresholds for enhancers predicted by different predictive models. Points indicate enrichment and recall at designated thresholds for each model. Error bars at the point represent 95% confidence intervals calculated from the natural log of enrichment. Dashed vertical line indicates 10% recall. **d.** Enrichment of variants at 10% recall for the same biosample pairs as **c**. Error bars for both plots represent 95% confidence intervals calculated from the natural log of enrichment. **e.** Enrichment–recall curves showing overlap of fine-mapped GWAS variants for hemoglobin A1c (HbA1c), mean corpuscular volume (MCV), and red blood cell count (RBC) in K562 predictions and variants for autoimmune disorders (AIDs) and lymphocyte count (Lym) in GM12878 predictions. Enrichment (y-axis) is calculated as the ratio of GWAS variants in predicted enhancers compared to distal noncoding variants; recall (x-axis) is calculated as the fraction of variants overlapping enhancers. Performance was evaluated at ∼100 score thresholds for continuous predictors. Points indicate enrichment and recall at suggested thresholds for each model, and error bars represent 95% confidence intervals calculated from the natural log of enrichment. **f.** Precision and recall of models at linking credible sets to known causal genes (inferred from independent signals with coding variants), with enhancers for each model binarized with their suggested threshold. Enhancer predictions for K562 and GM12878 were combined and evaluated against credible sets from 11 blood-related traits (**Table S5**). The top two target genes based on model score for each credible set were considered positive predictions. Error bars represent 95% confidence intervals based on the Wilson score interval. Faded points indicate performance of the models when predictions are intersected with PoPS, as in **h**. **g.** Same as **f**, except enhancer predictions were intersected with the top two genes linked to the credible set by PoPS. Faded points indicate performance of the models when predictions before intersecting with PoPS, as in **f**.

**Fig. N2.**
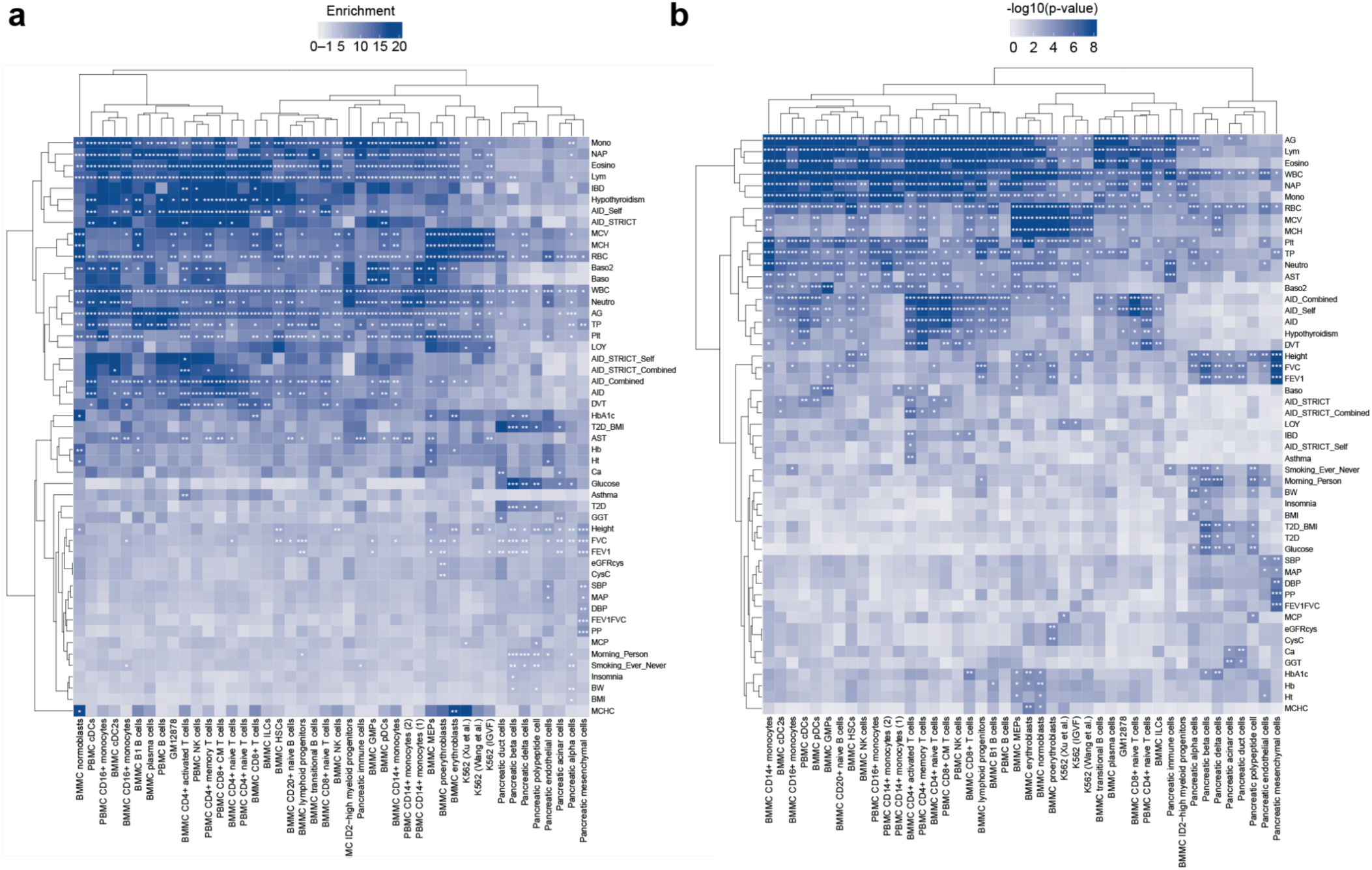
S-LDSC heritability enrichment in scE2G enhancers. **a.** Heatmap showing genome-wide enrichment computed with S-LDSC for 104 GWAS traits in scE2G*^Multiome^* predictions for 44 cell types. Stars indicate significance based on enrichment q-values (FDR-adjusted p-values): three stars indicate q < 0.001, two stars indicate q < 0.01, and one star indicates q < 0.05. The color scale is limited at the low end to an enrichment of 1. Rows and columns are hierarchically clustered with the “complete” method based on Euclidean distance. **b.** Heatmap showing -log10(p-value) for genome-wide enrichment computed with s-LDSC for 104 GWAS traits in scE2G*^Multiome^* predictions for 44 cell types. Stars indicate significance based on enrichment q-values as in **a**. Rows and columns are hierarchically clustered with the “complete” method based on Euclidean distance.

### Note S3. Enrichment of heritability of GWAS traits in scE2G enhancers by S-LDSC

We used stratified linkage disequilibrium score regression (S-LDSC) to identify cell types that had predicted enhancers enriched for genome-wide heritability of GWAS traits^51,52^. We computed enrichment of heritability for 104 traits^43^ in scE2G*^Multiome^* enhancers for 44 cell types and identified 654 significant trait-cell type pairs (**Table S8**, q < 0.05 after multiple hypothesis correction for each trait adjusting for the number of cell types compared, **Fig. N2a,b**) (14.3% of pairs). Subsetting to 25 traits relevant to the cell types analyzed (combined autoimmune disorders, basophil count, body fat percentage, body mass index, body weight, eosinophil count, random glucose, hemoglobin, hemoglobin A1c, hematocrit, hypothyroidism, inflammatory bowel disease, IGF1, lymphocyte count, mean corpuscular hemoglobin, mean corpuscular hemoglobin concentration, mean corpuscular volume, monocyte count, neutrophil count, platelet count, red blood cell count, type 2 diabetes, type 2 diabetes (BMI-adjusted), white blood cell count, waist-height-ratio-adjusted body mass index), 373 trait-cell type pairs had significant enrichment (34.0% of pairs).

## Supplementary Tables

**Table S1. Published models**

Summary of previously-published models benchmarked here, including the score metrics and thresholding criteria used for each analysis.

**Table S2: CRISPR benchmark pairwise comparisons**

Results from bootstrapping (10,000 replicates) of difference in performance between each pair of models on the CRISPR benchmark (element-gene pairs with distance to TSS < 1Mb).

**Table S3: CRISPR benchmark performance intervals**

Table with results from bootstrapping (N = 10,000 replicates) of confidence intervals for performance metrics for each model on the CRISPR benchmark (element-gene pairs with distance to TSS < 1Mb)

**Table S4. eQTL benchmarking cell type pairings**

Description of prediction and eQTL cell type pairings used for eQTL benchmarking.

**Table S5. GWAS benchmarking trait–cell type pairings**

Description of traits and prediction cell type pairings used for GWAS benchmarking

**Table S6. scE2G*^Multiome^* predictions**

Descriptions of each cell type to which scE2G was applied and links to scE2G_Multiome predictions for each cell type. See https://github.com/anderssonlab/scE2G_analysis/blob/v1.0/5.Prediction_Properties/scE2G_prediction_columns.tsv for descriptions of each column in the scE2G prediction files.

**Table S7: scE2G*^Multiome^* and scE2G*^ATAC^* prediction properties**

Table describing genome-wide properties of scE2GMultiome and scATAC predictions in each cell type. See https://github.com/anderssonlab/scE2G_analysis/blob/v1.0/5.Prediction_Properties/prediction_properties.tsv for definitions of each metric.

**Table S8. GWAS trait heritability enrichment in scE2G*^Multiome^* enhancers**

Results from stratified linkage-disequilibrium score regression (S-LDSC) analysis of heritability enrichment of GWAS traits in scE2G*^Multiome^* enhancers. Results are sorted by increasing enrichment p-value.

**Table S9. Gene prioritization for noncoding GWAS variants**

Gene prioritization tables for BMMCs, PBMCs, and pancreatic islets detailing each credible set, causal gene, and cell types are predicted by scE2G*^Multiome^*. For BMMC and PBMC cell types, the following traits were considered: RBC, WBC, Mono, Lym, Eosino, Baso, Neutro, Plt, MCH, MCHC, MCV, Hb, HbA1c, Ht, Glucose. For pancreatic islet cell types, the following traits were considered: Glucose, HbA1c, WHRadjBMI, Ht, T2D_BMI. Only trait-cell type pairs in which variants for the trait were significantly enriched in enhancers by at least 5-fold were prioritized.

**Table S10. 10x Multiome data sources**

Sources for 10x Multiome datasets used in this study

**Table S11. 10x Multiome data quality control thresholds**

Thresholds used to select high-quality cells from each 10x Multiome dataset

## Methods

### 1. Genome build

All coordinates in the human genome are reported using build GRCh38 unless otherwise specified.

### 2. Gene reference file for scE2G and benchmarking analyses

We used the following set of genes and promoter coordinates when building scE2G and conducting all benchmarking analyses: https://github.com/EngreitzLab/ENCODE_rE2G/blob/dev/reference/CollapsedGeneBounds.hg38.TSS500bp.bed. This file contains one promoter per gene symbol, which we defined as a 500 base pair region centered around the RefSeq^1^ TSS with the largest number of coding isoforms. To curate the set of genes, we matched the gene symbols with ENSEMBL IDs using the HUGO database^2^ and manual annotation, then used the GENCODE v29 database^3^ to annotate each gene with its “gene type”. We retained genes that 1) were included in the GENCODE v29 database and 2) had a gene type of “protein_coding,” “processed_transcript,” or “lincRNA” for a total of 20,666 genes.

### 3. Preprocessing of 10x Multiome Data

#### 3.1. Build reference genome

The GRCh38 genome FASTA file was downloaded from the ENCODE Project (https://www.encodeproject.org/files/GRCh38_no_alt_analysis_set_GCA_000001405.15/@@download/GRCh38_no_alt_analysis_set_GCA_000001405.15.fasta.gz), and the gene annotation GTF file was obtained from GENCODE (https://ftp.ebi.ac.uk/pub/databases/gencode/Gencode_human/release_43/gencode.v43.chr_patch_hapl_scaff.annotation.gtf.gz). We created a reference using the `cellranger-arc` (v2.0.2) “`mkref`” tool, following the guidelines provided by 10x Genomics (https://www.10xgenomics.com/support/software/cell-ranger-arc/latest/analysis/inputs/mkref). The Wang *et al.* K562 dataset mixes human K562 and mouse A20 cells. To distinguish human K562 cells from mouse A20 cells, we downloaded the mouse reference (mm10) from https://cf.10xgenomics.com/supp/cell-arc/refdata-cellranger-arc-mm10-2020-A-2.0.0.tar.gz

#### 3.2. Prepare raw FASTQ files

For the Xu *et al.* K562^4^ and PBMC datasets, we directly downloaded the raw FASTQ files (**Table S10**). For the GM12878^5^, BMMC^6^ datasets, we downloaded BAM format files (**Table S10**), and converted the BAM files to FASTQ files using the cellranger-arc “bamtofastq” tool (https://www.10xgenomics.com/support/software/cell-ranger/latest/miscellaneous/cr-bamtofastq). For the Wang *et al.* K562^7^ and pancreatic islets^8^ datasets, we downloaded SRA format files using the sratoolkit (v3.0.0) with the commands “`prefetch -X 100G -pv -O sra SRR_accession_number`”, and converted SRA files to FASTQ files using “`fastq-dump --split-files --gzip --outdir output_directory_path sra_file_path`”. We renamed the FASTQ files in accordance with the 10x Genomics guidelines (https://www.10xgenomics.com/support/software/cell-ranger-arc/latest/analysis/inputs/specifying-input-fastq-count) if the file names did not meet the requirements of cellranger-arc count.

#### 3.3. Obtaining K562 (this study) dataset

For the IGVF K562 dataset, we downloaded raw FASTQ files from the IGVF Data Portal (https://data.igvf.org/analysis-sets/IGVFDS5839UBFZ/). Count matrices were generated using `cellranger count` (v6.0.0) with the GRCh38 reference transcriptome. Fragment files were generated using `cellranger-atac count` (v2.0.0) with the GRCh38 reference genome.

#### 3.4. Run cellranger-arc pipeline

We created a libraries CSV file for each sample to delineate the input file paths, and ran the cellranger-arc count tool following the 10x Genomics guidelines (https://www.10xgenomics.com/support/software/cell-ranger-arc/latest/analysis/running-pipelines/single-library-analysis).

### 4. Quality control for 10x Multiome data

#### 4.1. Quality control metrics

We implemented following metrics to select of high-quality cells from the 10x Multiome data:

- gex_umis_count: UMI count in scRNA-seq
- atac_fragments: fragment count in scATAC-seq
- nucleosome_signal: ratio of mononucleosome fragments to nucleosome-free fragments
- TSS.enrichment: scATAC-seq TSS enrichment score defined by ENCODE (https://www.encodeproject.org/data-standards/terms/)
- percent.mt: percentage of mitochondrial transcripts

The `gex_umis_count` and `atac_fragments` metrics were extracted from the “per_barcode_metrics.csv” file, which is generated by the cellranger-arc count pipeline. The `atac_fragments`, `TSS.enrichment`, and `percent.mt` metrics were calculated using the Signac^9^ and Seurat^10,11^ R functions `Signac::NucleosomeSignal, Signac::TSSEnrichment`, and `Seurat::PercentageFeatureSet(pattern = “^MT-”)`, respectively. We defined thresholds for these metrics based on their distributions within each dataset. These thresholds are detailed in **Table S11**.

#### 4.2. Identifying human K562 cells in Wang et al.^7^ dataset

In the Wang *et al.*^7^ dataset, human K562 cells were mixed with mouse A20 cells. We calculated the percentage of mapped reads (% mapped = # mapped_reads / # raw_reads) to the human genome, and selected cells with % mapped > 95% in the human genome for both scRNA-seq and scATAC-seq as human K562 cells.

#### 4.3. IGVF K562 quality control

Cells were thresholded to exclude cells with fewer than 10^3^ total fragments before a joint peak matrix was computed across two 10x Genomics microfluidic lanes. Peak matrices were then thresholded to exclude cells with fewer than 100 RNA UMIs, merged, and pre-processed using standard Seurat and Signac workflows. Sample of origin demultiplexing was then performed using the deMULTIplex2 ^12^ R package and doublet-enriched clusters and unclassified cells were removed. Cleaned data was then re-processed using Seurat and Signac, and K562 cells were subsetted for downstream analyses.

### 5. Cell-type annotation

#### 5.1. BMMC dataset

We downloaded the processed h5ad file from Luecken *et al.*^6^ (available at NCBI GEO https://www.ncbi.nlm.nih.gov/geo/query/acc.cgi?acc=GSE194122) and extracted the cell type information from it.

#### 5.2. Pancreatic islet dataset

We conducted scRNA-seq analysis using Seurat (v5.0.3)^10^ to perform cell clustering:

i. Normalized the RNA UMI count matrix using `Seurat::NormalizeData`.
ii. Identified variable genes with `Seurat::FindVariableFeatures`.
iii. Scaled the normalized RNA matrix for variable genes using `Seurat::ScaleData`.
iv. Performed PCA using `Seurat::RunPCA`.
v. Defined neighboring cells using `Seurat::FindNeighbors`.
vi. Identify cell clusters using `Seurat::FindClusters`.

Subsequently, we defined the corresponding cell types based on marker gene expression: beta cell (*INS*+), alpha cell (*GCG*+), delta cell (*SST*+), PP cell (*PPY*+), ductal cell (*CFTR*+), acinar cell (*REG1A*+), endothelial cell (*PECAM1*+), mesenchymal cell (*COL3A1*+) and immune cell (*PTPRC*+). Cell clusters expressing multiple markers were considered as doublets. Additionally, beta cells from one donor (A0027) exhibited notable batch effects. Both doublets and beta cell clusters with batch effects were excluded from downstream analyses.

#### 5.3. PBMC dataset

To facilitate the annotation of cell types in PBMC 10x Multiome data, we downloaded the Healthy Hematopoiesis scRNA-seq dataset from https://github.com/GreenleafLab/MPAL-Single-Cell-2019. We then employed Harmony^13^ (v1.2.0) to integrate the PBMC and Healthy Hematopoiesis scRNA-seq data. Based on the results from Harmony, we performed cell clustering using Seurat. Utilizing the cell type annotation information from the Healthy Hematopoiesis scRNA-seq dataset, we identified the most prevalent cell type in our new clustering results as the designated cell type.

### 6. Preparing input files for scE2G

The scE2G pipeline requires input data including RNA count matrices, ATAC fragment files, and their corresponding *.tbi index files for each cell cluster.

#### 6.1. RNA count matrix

We exported the RNA count matrix from the Seurat object in CSV format. Rows of the matrix correspond to genes, while columns correspond to cells. The first row contains the names of the cells, and the first column contains the names of the genes.

#### 6.2. ATAC fragment file

We utilized the R function `Signac::SplitFragments` to split raw fragment files into separate fragment files for each sample and each cell type. We then merged fragment files of the same cell type into a single fragment file. If the cell names in the fourth column of the fragment file do not match the cell names in the RNA count matrix (*e.g.*, cell names in the RNA count matrix include a sample name prefix), it is necessary to modify the cell names in the fragment file to match those in the RNA count matrix. Subsequently, we sorted each cell type’s fragment file using `sortBed` from bedtools (v2.31.0), and then compressed and indexed each fragment file using bgzip and tabix from htslib^14^ (v1.18), respectively. Example data can be found at scE2G repository (https://github.com/EngreitzLab/scE2G/tree/v1.0/resources/example_chr22_multiome_cluster).

### 7. Kendall correlation

We applied a non-parametric measurement, the Kendall correlation (also known as Kendall rank correlation coefficient and Kendall’s τ coefficient)^15^, to quantify the correlation between enhancer accessibility and gene expression. We extracted MACS2^16^ peaks that overlap with candidate enhancers defined in the ABC pipeline (see **Methods 9.1.1**) and generated the ATAC count matrix using the `Signac::FeatureMatrix` function. We binarized the ATAC fragment count matrix using R `Signac::BinarizeCounts` function, and normalized the RNA UMI count matrix using R `Seurat::NormalizeData` function. We then computed the Kendall correlation τ_b_ between binarized enhancer accessibility and normalized gene expression for each candidate enhancer-gene pairs.

Briefly, Kendall’s correlation is based on the concordance and discordance between pairs of observations. A pair is considered concordant if the ranks for both elements agree (i.e., if the enhancer accessibility in one pair is higher than in another, the same relationship holds for the gene expression). Conversely, a pair is discordant if the ranks disagree. When all values for both variables are unique, τ is calculated as 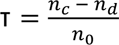, where *n_C_* is the number of concordant pairs, *n_d_* is the number of discordant pairs, and *n_o_* is the total number of pairs. Due to the binarization of enhancer accessibility, there are many ties in enhancer accessibility. To address these ties, we used the τ_b_ formula, which accounts for ties: 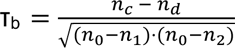, where *n* is the number of cells, 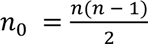, *c* is the number of tied pairs within the first variable, and **n*_2_*, is the number of tied pairs within the second variable.

To accelerate the calculation of Kendall correlation, we have developed code using R and Rcpp specifically for computing binarized matrices (ATAC accessibility) and continuous matrices (gene expression). Briefly, we calculated the Kendall correlation for one gene and multiple enhancers at a time by first reordering the binarized ATAC accessibility matrix based on the descending order of gene expression. This sorted matrix was then processed using an Rcpp function, count_diff, which efficiently computed the difference between concordant and discordant pairs for each enhancer. Below is pseudocode for the `count_diff` function:

*Function count_diff(y_matrix_sorted)*

*Input: y_matrix_sorted, a logical matrix with rows sorted by another variable (e.g., gene expression)*

*Initialize n as number of rows in y_matrix_sorted*

*Initialize m as number of columns in y_matrix_sorted*

*Initialize result as a numeric vector of length m*

*For j from 0 to m-1*

*Initialize concordant to 0*

*Initialize disconcordant to 0*

*Initialize cumsum to 0*

*For i from 0 to n-1*

*Set isAccessible to the value at y_matrix_sorted[i, j] (true for accessible, false for inaccessible)*

*Add isAccessible to cumsum*

*If isAccessible is true*

*Increment disconcordant by (i + 1 - cumsum)*

*Else*

*Increment concordant by cumsum*

*Set result[j] to concordant - disconcordant*

*Return*

*End Function*

Subsequently, our R function adjusted these differences for any ties in gene expression values to ensure accurate correlation calculations. Finally, we calculated Kendall correlation τ_b_ using these adjusted values. See https://github.com/EngreitzLab/scE2G/blob/v1.0/workflow/scripts/compute_kendall.R for the full implementation.

### 8. ARC-E2G score

According to our CRISPRi benchmarking results, the ABC model and the Kendall correlation each performed best as single features dependent and independent of 3D contact or distance, respectively (**Fig. 2b,c**). However, the performance of the Kendall correlation, and therefore its weight in the scE2G logistic regression model, was dependent on sequencing and cell depth. Accordingly, to improve robustness across sequencing depths, we integrated the ABC model and the Kendall correlation into a single feature called Activity, Responsiveness, and Contact (ARC)-E2G scores.

We observed that the distributions of the Kendall correlation τ and the ln(ABC_score) approximate the normal distribution (**Fig. S1a,b**). To make their values comparable in magnitude, we scaled them by subtracting the mean μ and then dividing by the standard deviation σ. We found that the τ_scaled_ and ln(ABC_score)_scaled_ are linearly positively correlated, and that CRISPRi^+^ E-G pairs are enriched in the area where both values are high compared to CRISPRi^-^ E-G pairs (**Fig. S1c,d**). This suggests that we can integrate them based on their linear relationship. Consequently, we fitted a linear regression between scaled Kendall correlation and scaled ln(ABC score):

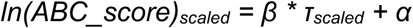

We considered that points on the orthogonal line to this regression line, described by *y = −1 / β * x + α’*, have the same predicted value, where α’ is the intercept of the orthogonal line on the y-axis. Higher values of α’ indicate a stronger likelihood that the line corresponds to CRISPRi^+^ E-G pairs. The intercept α’ is calculated as follows:

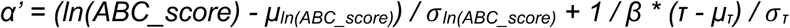

We define the ARC-E2G score as follows to ensure it aligns with the magnitude of the ABC score and increases monotonically with the intercept α’ of the orthogonal line:

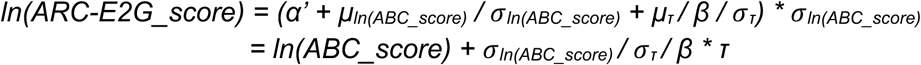

We observed that the ranking of ARC-E2G corresponds well with the density of CRISPRi^+^ relative to CRISPRi^-^ E-G pairs (**Fig. S1d,e**). See https://github.com/EngreitzLab/scE2G/blob/v1.0/workflow/scripts/compute_arc_e2g.R for the full implementation.

### 9. Training and applying scE2G

We implemented a training and application pipeline for scE2G that uses the Activity-by-Contact (ABC)^17^ codebase to define all candidate element-gene pairs, quantify activity at each element, and compute the ABC score for each element-gene pair, and that uses the ENCODE-rE2G^18^ codebase to annotate element-gene pairs with additional activity-based features. In the scE2G pipeline, we compute the Kendall correlation between element accessibility and gene expression, calculate the ARC-E2G score that integrates the Kendall correlation and ABC score (see **Methods 7,8**), and implement additional score normalization and filtering to account for sequencing depth variability and gene expression data. The scE2G pipeline, including ENCODE-rE2G submodule and ABC sub-submodule, is implemented here: https://github.com/EngreitzLab/scE2G/releases/tag/v1.0

#### 9.1. Processing pseudobulk ATAC-seq data (implemented in ABC pipeline)

##### 9.1.1. Define candidate element-gene pairs

We call peaks on a tagAlign file generated from the pseudobulk ATAC fragment file for each cell cluster using MACS2 with parameters from the ENCODE ATAC-seq Processing Pipeline^19^. We count reads in each peak with P<0.1 and identify the 150,000 peaks with the highest read counts. We take 500 base-pair elements centered on the summits of these peaks, merge overlapping elements, remove elements overlapping ENCODE blacklisted regions (regions of the genome that have been observed to accumulate anomalous number of reads in epigenetic sequencing experiments)^20^, add 500 base-pair promoter elements centered on the transcription start site of each gene (from **Methods** 2), and again merge overlapping regions. We define the resulting list of regions as ‘candidate elements,’ and annotate each as a promoter (elements overlapping regions from the promoter reference file), genic (elements within an annotated gene body), or intergenic (all other elements). We define all genic and intergenic elements as ‘distal’ elements. We generate the list of candidate element-gene pairs by computing all combinations of candidate elements with any promoter within 5 Mb.

##### 9.1.2. Annotating candidate elements with normalized ATAC signal

We count reads in each candidate element from the tagAlign file, and double the count for elements on the sex chromosomes to simulate two copies. We then rank-normalize the ATAC reads per million for distal elements and promoters to K562 reference values (derived from the original ABC paper)^17^ for each rank in their respective category. Briefly, we rank candidate elements from each category by ATAC reads per million, and replace the read count with the interpolated read count from the reference list. This normalization is necessary for comparison of values across biosamples and minimizes technical variability from the experimental set up. The ATAC read count after normalization is used in all downstream calculations.

##### 9.1.3. Determining active genes

We remove element-gene pairs from consideration for any gene that has an ‘inactive promoter’ based on the promoter ATAC signal. Here, we exclude genes with a promoter activity quantile less than 0.3.

##### 9.1.4. Estimated 3D contact between each element-gene pair

Because it is rare to have cell-type specific Hi-C data corresponding to cell clusters from single-cell experiments, we estimate 3D contact using a power law relationship with genomic distance. We have previously shown that a relationship in the form of contact = exp(scale + −1 * gamma * ln(distance)) fits Hi-C contact frequencies well^17^. The *gamma* and *scale* parameters were previously computed by fitting a linear regression between log-transformed distance and K562 Hi-C contact, binned at 5-kb resolution^18^. For genomic distances below 5 kb, we set estimated contact as the power law contact at a distance of 5 kb.

##### 9.1.5. Calculate ABC score for each element-gene pair

We calculate the ABC score between each element-gene pair on a per-gene basis. The numerator of the ABC score for a given element-gene pair is the normalized ATAC signal at the element multiplied by the power law contact between the element and gene. The denominator of the ABC score for a given gene is equal to the sum across all candidate element-gene pairs of the normalized ATAC signal at the element multiplied by the power law contact between the element and gene. Finally, the ABC scores for ‘self-promoter’ element-gene pairs (promoters linked to their target gene) are set to 1.

#### 9.2. Annotating genome-wide element-gene pairs with additional features

In addition to the ABC score, the Kendall correlation, and ARC-E2G score (see **Methods 7,8**), we annotated each element-gene pair with nine other values related to chromatin accessibility, genomic distance, and gene density to use as input features for scE2G. We included the following four features that were output directly from the ABC pipeline: distance to transcription start site, ATAC signal at the candidate element, ATAC signal at the promoter, and power law contact. We further computed the number of TSSs and number of other elements between the element and promoter, as well as the number and sum of activity of peaks within 5 kb of the element (implemented in ENCODE-rE2G pipeline). Finally, each promoter was classified based on ‘ubiquitously expression.’

##### 9.2.1. Classifying promoters as ubiquitously-expressed

We annotate promoters as ubiquitously-expressed using a classification obtained from previous work^21^. Briefly, CAGE data from the FANTOM5 consortium across 1,829 biosamples (downloaded from https://fantom.gsc.riken.jp/5/datafiles/latest/extra/gene_level_expression/hg19.gene_phase1and2combined_tpm.osc.txt.gz on 19 February 2022) was analyzed to obtain expression (in TPM) for each gene. Ubiquitously-expressed genes were defined as those expressed at ≧1 TPM in ≧95% of biosamples.

#### 9.3. Training scE2G on CRISPR perturbation dataset (implemented in ENCODE-rE2G pipeline)

We trained logistic regression classifiers on the K562 CRISPR perturbation dataset, where positive element-gene pairs were defined as cases where perturbing the element had a significant and negative effect size on expression of the target gene. Using this framework, we selected final sets of features for two models: scE2G*^ATAC^* which uses just scATAC-seq data and scE2G*^Multiome^* which uses multiomic data.

##### 9.3.1. Training logistic regression classifiers

We annotated the element-gene pairs in the CRISPR dataset by overlapping them with the genome-wide element-gene pairs (at least one base pair overlap) computed on a deeply-sequenced K562 10x Multiome dataset^4^. We applied a variance-stabilizing transformation log(|x| + ϵ), with ϵ = 0.01, to all the features before feeding them into a logistic regression model implemented with sklearn’s `LogisticRegression` function. To maintain calibrated probability scores as an output of the logistic regression, we did not use any penalty. To evaluate the performance of the logistic regression models on the CRISPR dataset, we used chromosome-wise cross validation by holding out one chromosome, training a model on the rest, and using this model to score each element-gene pair on the held-out chromosome. For scE2G*^Multiome^*, we then implement a gene expression filter (specified by a minimum TPM) by setting the score of all element-gene pairs where the gene is expressed below the TPM threshold to 0. We also train a model on all chromosomes, which is used to apply scE2G to new cell types.

##### 9.3.2. Selecting features for final models

To select the final set of features for each scE2G model, we used several approaches. We applied a best-subsets approach to calculate the area under the precision-recall curve (AUPRC) on the CRISPR benchmark for models comprising every possible subset of the input features, and selected the set of features for each model that minimized number of features while maintaining near-optimal performance. We performed additional feature analyses including forward sequential feature selection (iteratively choosing the feature that contributes to the largest increase in AUPRC) and ablation of single features or categories of features to validate the importance of our chosen features in the final models. When choosing features for scE2G*^Multiome^*, we used initial feature sets with ABC score and the Kendall correlation as separate features to determine the contribution of each feature independently, as well as with the ARC-E2G score in place of the two features. We saw similar results with both feature sets.

The final feature set for scE2G*^Multiome^* consists of six features:: ARC-E2G score, ubiquitous expression of promoter, number of candidate elements between element and gene, number of transcription start sites between element and gene, number of candidate elements within 5 kb of the element, and normalized ATAC signal at the promoter. The final feature set for scE2G*^ATAC^* is the same, except with the ABC score in place of the ARC-E2G score.

##### 9.3.3. Applying scE2G to new cell types

To apply the trained scE2G models to new cell types, we first compute a feature table for all candidate element-gene pairs as described above. We then apply the logistic regression model trained on all chromosomes to assign a score to each element-gene pair. We quantile-normalize the raw scores to the scE2G scores computed on the full-depth K562 dataset used for model training to maintain robustness across sequencing depths. For scE2G*^Multiome^*, we then apply the gene expression filter by setting the score for any element-gene pair in which the gene is expressed below the designated TPM threshold to 0. For applications requiring binarized enhancer predictions, we used the score threshold determined as the score yielding 70% recall when evaluating the cross-validated scores on the CRISPR benchmark. The thresholds for scE2G*^Multiome^* and scE2G*^ATAC^* were determined to be 0.164 and 0.183, respectively.

### 10. Additional versions of scE2G

#### 10.1. scE2G downsample analysis

We performed downsampling on Xu *et al.* K562^4^ dataset to reduce the number of cells, the mean RNA UMI count, and the mean ATAC fragment count to the levels listed in **Fig. S10a**. We used the `base::sample` function in R to select a specified number of cell barcodes. We then sampled the RNA count matrix based on a binomial distribution using the `stats::rbinom` function in R. For ATAC fragment downsampling, we used the `base::sample` function to extract a specific number of fragments. See https://github.com/anderssonlab/scE2G_analysis/blob/v1.0/1.Preprocess_Data/K562_Xu/downsample.R for the full implementation.

We ran the scE2G pipeline on each downsampled dataset and assessed all scE2G prediction outcomes on the CRISPR benchmark. To determine the minimal sequencing depth requirement, we calculated the total ATAC fragments based on the mean ATAC fragment count multiplied by the cell count. Then, we explored the relationship between total ATAC fragments and AUPRC, precision, and recall (**Fig. 4**). We also trained scE2G models on each of the downsamples to examine the stability of optimal model weights across cell number and sequencing depth.

#### 10.2. scE2G meta-cell analysis

We followed the SEACells^22^ tutorials https://github.com/dpeerlab/SEACells/blob/main/notebooks/SEACell_computation.ipynb to construct 90 meta-cells based on Xu *et al.*^4^ K562 scRNA-seq data. According to the single-cell to meta-cell assignments determined in SEACells (**Fig. 3f left**), we aggregated single-cell RNA UMI counts and ATAC fragment counts into meta-cell RNA UMI count matrix and ATAC fragment count matrix, respectively. We normalized the meta-cell RNA UMI count matrix using `Seurat::LogNormalize` and normalized the meta-cell ATAC fragment count matrix by dividing by the total counts of the corresponding meta-cell and then multiplying by 1,000,000. Based on the normalized RNA and ATAC matrices, we computed the Kendall correlation (meta-cell) using the `fastkendall::fastkendall` function for each candidate element-gene pair defined in the scE2G pipeline. Subsequently, we retrained a scE2G model using the ABC-score and Kendall correlation (meta-cell) as independent features, along with five other features used in the original scE2G model. Finally, using the retrained model, we performed scE2G (meta-cell) predictions using a hold-one-chromosome-out cross-validation approach.

### 11. Published predictors

We run the published predictors on each 10x Multiome dataset using the same input RNA UMI count matrix and ATAC fragment count matrix, unless otherwise specified. We obtain the RNA UMI count matrix as described in **Methods 6.1**. For datasets containing only one cell type (K562 and GM12878), we obtain the ATAC fragment count matrix as described in **Methods 7**. For datasets containing multiple cell types (PBMC and BMMC), we extracted MACS2 peaks that overlap with candidate enhancers defined in the ABC pipeline (see **Methods 9.1.1**) for each cell type, then merged peaks from all cell types, with peaks being merged if they overlap by at least 1 bp. Subsequently, we generated the ATAC count matrix using the `Signac::FeatureMatrix` function. For fair benchmarking, we uniformly make predictions for peak-gene pairs within 1 Mb TSS distance. We also made a version using the default settings of the corresponding method on the Xu *et al.* K562 dataset (**Fig. S6**).

#### 11.1. Signac^9^

We ran Signac across all cell types within a dataset. For the uniform prediction within a 1Mb TSS distance, we normalized the RNA UMI count matrix using Seurat::NormalizeData function. We then calculated the Pearson correlation between normalized gene expression and enhancer accessibility for each candidate element-gene pair. To select background peaks, we employed `Signac::RegionStats` to compute GC content and region lengths for all peaks, and identified 200 background peaks with similar GC content, sequence length, and total ATAC fragment count using the `Signac::MatchRegionStats` function. Subsequently, we calculated the Pearson correlation between target gene expression and background peak accessibility. A z-score is computed for each enhancer-gene pair using the formula z = (r-μ)/δ, where μ is the mean background Pearson correlation and δ is the standard deviation of the background Pearson correlations for the peak. We then computed a p-value for each enhancer-gene pair using a one-sided z-test. See https://github.com/anderssonlab/scE2G_analysis/blob/v1.0/2.Prediction/K562_Xu/Signac/Signac.240517.ipynb for more details.

For the default setting prediction, we normalize the RNA UMI count matrix using the `Seurat::SCTransform` function, then conduct predictions using `Seurat::LinkPeaks` function. See https://github.com/anderssonlab/scE2G_analysis/blob/v1.0/2.Prediction/K562_Xu/Signac.default/Signac.240611.ipynb for more details.

#### 11.2. FigR^23^

We ran FigR across all cell types within a dataset. For uniform prediction within a 1Mb TSS distance, we normalized the RNA UMI count matrix using the `Seurat::NormalizeData` function, and the ATAC fragment count matrix using the `FigR::centerCounts` function. We then calculated the Spearman correlation between normalized gene expression and enhancer accessibility for each candidate enhancer-gene pair. To select background peaks, we used `chromVAR::addGCBias` to compute GC content for all peaks and identified 100 background peaks with similar GC content and total ATAC fragment count using the `chromVAR::getBackgroundPeaks` function. Subsequently, we calculated the Spearman correlation between target gene expression and background peak accessibility. A z-score for each enhancer-gene pair was computed using the formula z = (r-μ)/δ, where μ is the mean background Spearman correlation and δ is the standard deviation of the background Spearman correlations for the peak. We then computed a p-value for each enhancer-gene pair using a one-sided z-test. See https://github.com/anderssonlab/scE2G_analysis/blob/v1.0/2.Prediction/K562_Xu/FigR/FigR.240521.ipynb for more details.

For the default setting prediction, we normalize the RNA UMI count matrix using the `Seurat::SCTransform` function and then conduct predictions using the `FigR::runGenePeakcorr` function. See https://github.com/anderssonlab/scE2G_analysis/blob/v1.0/2.Prediction/K562_Xu/FigR.default/FigR.240612.ipynb for more details.

#### 11.3. SnapATAC^24^

We ran SnapATAC across all cell types within a dataset. For uniform prediction within a 1Mb TSS distance, we normalized the RNA UMI count matrix using the `Seurat::NormalizeData` function and binarized the ATAC fragment count matrix using the `Signac::BinarizeCounts` function. We then fitted a logistic regression model for each element-gene pair using `stats::glm(ATAC_accessibility ∼ Gene_Expression, family = binomial(link = “logit”))`. See https://github.com/anderssonlab/scE2G_analysis/blob/v1.0/2.Prediction/K562_Xu/SnapATAC/SnapATAC.240522.ipynb for more details.

For the default setting prediction, we normalize the RNA UMI count matrix using the `Seurat::SCTransform` function and binarize the ATAC fragment count matrix using the `Signac::BinarizeCounts` function. Then, we conduct predictions using the `SnapATAC::predictGenePeakPair` function. See https://github.com/anderssonlab/scE2G_analysis/blob/v1.0/2.Prediction/K562_Xu/SnapATAC.default/SnapATAC.240612.ipynb for more details.

#### 11.4. SCENT^25^

We ran SCENT for each cell type. For uniform prediction within a 1Mb TSS distance, we normalized the RNA UMI count matrix using the `Seurat::NormalizeData` function and binarized the ATAC fragment count matrix using the `Signac::BinarizeCounts` function. We then fitted a Poisson regression model for each element-gene pair using `stats::glm(exprs ∼ atac + log.nUMI + percent.mito, family = “poisson”)`, where exprs represents normalized gene expression, atac denotes binarized ATAC accessibility, log.nUMI is the log-transformed total RNA UMI count, and percent.mito is the percentage of mitochondrial UMIs. Subsequently, we performed bootstrapping using the `boot::boot` function and calculated the false discovery rate (FDR) using `SCENT::basic_p`. Due to the time-consuming nature of bootstrapping, we limited the bootstrapping process to 500 iterations. To distinguish between activation and repression effects, we log-transformed the FDR and assigned it a negative value if the beta coefficient of ATAC accessibility in the Poisson regression model is less than zero. See https://github.com/anderssonlab/scE2G_analysis/blob/v1.0/2.Prediction/K562_Xu/SCENT/SCENT_500.240604.ipynb for more details.

For the default setting prediction, we perform similar processing as with uniform prediction, but increase the number of bootstrapping iterations to a maximum of 2,500 times, and limit the TSS distance to within 500 kb. See https://github.com/anderssonlab/scE2G_analysis/blob/v1.0/2.Prediction/K562_Xu/SCENT/SCENT_2500.240606.ipynb and https://github.com/anderssonlab/scE2G_analysis/blob/v1.0/2.Prediction/K562_Xu/SCENT.default/SCENT.extract_2500.240622.ipynb for more details.

#### 11.5. SCENIC+^26^

We ran SCENIC+ across all cell types within a dataset. Following the SCENIC+ tutorials (https://scenicplus.readthedocs.io/en/latest/tutorials.html), we preprocessed scATAC-seq data using pycisTopic (see https://github.com/anderssonlab/scE2G_analysis/blob/v1.0/2.Prediction/K562_Xu/ScenicPlus/preprocessing.ATAC.ipynb) and scRNA-seq data using Scanpy^27^ (see https://github.com/anderssonlab/scE2G_analysis/blob/v1.0/2.Prediction/K562_Xu/ScenicPlus/preprocessing.RNA.ipynb). Subsequently, we used the commands `scenicplus prepare_data prepare_GEX_ACC` to prepare multiome input data, and `scenicplus prepare_data search_spance` to generate candidate element-gene pairs. Finally, we employed the `scenicplus grn_inference region_to_gene` command to make predictions. For uniform predictions, we configured `scenicplus prepare_data search_spance` with `--upstream 1000 1000000` and `--downstream 1000 1000000` to specify a maximum TSS distance as 1Mb. For default settings predictions, we adhered to the default parameters: `--upstream 1000 150000` and `--downstream 1000 150000`. See https://github.com/anderssonlab/scE2G_analysis/blob/v1.0/2.Prediction/K562_Xu/ScenicPlus/region_to_gene.ipynb for more details.

#### 11.6. Cicero^28^

We ran Cicero across all cell types within a dataset. We executed the Cicero pipeline according to the tutorial at https://cole-trapnell-lab.github.io/cicero-release/docs_m3/. For the uniform prediction, we configured the `cicero::estimate_distance_parameter` function with `distance_constraint`

`= 1000000` and `window = 2000000` to specify the maximum TSS distance as 1Mb. For the default setting prediction, we adhered to the default parameters: `distance_constraint = 250000` and `window = 500000`. See https://github.com/anderssonlab/scE2G_analysis/blob/v1.0/2.Prediction/K562_Xu/Cicero/Cicero.240525.ipynb (Uniform Prediction) and https://github.com/anderssonlab/scE2G_analysis/blob/v1.0/2.Prediction/K562_Xu/Cicero.default/Cicero.240525.ipynb (Default Setting) for more details.

#### 11.7. DIRECT-NET^29^

We ran DIRECT-NET across all cell types within a dataset. We ran the DIRECT-NET pipeline following the demo https://htmlpreview.github.io/? https://github.com/zhanglhbioinfor/DIRECT-NET/blob/main/tutorial/demo_DIRECTNET_PBMC.html. We adapted the `DIRECTNET::Aggregate_data` function for compatibility with the new version of Seurat, and modified the `DIRECTNET::Run_DIRECT_NET` function to enable setting the maximum TSS distance. We configured the maximum TSS distance to be 1,000,000 for uniform prediction and 250,000 for default setting prediction, respectively. See https://github.com/anderssonlab/scE2G_analysis/blob/v1.0/2.Prediction/K562_Xu/DIRECTNET/DIRECTNET.240527.ipynb (Uniform Prediction) and https://github.com/anderssonlab/scE2G_analysis/blob/v1.0/2.Prediction/K562_Xu/DIRECTNET.default/DIRECTNET.240527.ipynb (Default Setting) for more details.

#### 11.8. ArchR^30^

We ran ArchR across all cell types within a dataset. We created an ArchR project following the tutorial available at https://www.archrproject.com/articles/Articles/tutorial.html. For the element-gene predictions, we used the `ArchR::addPeak2GeneLinks` function as described in https://www.archrproject.com/bookdown/peak2genelinkage-with-archr.html. In the uniform prediction, we set `maxDist = 1000000` within the ArchR::addPeak2GeneLinks function to specify the maximum TSS distance as 1Mb, while for the default setting prediction, we adhered to the default `maxDist = 250000`. See https://github.com/anderssonlab/scE2G_analysis/blob/v1.0/2.Prediction/K562_Xu/ArchR/ArchR.240528.ipynb (Uniform Prediction) and https://github.com/anderssonlab/scE2G_analysis/blob/v1.0/2.Prediction/K562_Xu/ArchR.default/ArchR.240528.ipynb (Default Setting) for more details.

#### 11.9. ABC model

We used the ABC scores of each cell type generated within our scE2G pipeline as described in **Methods 9.1**, which is implemented in the ABC pipeline.

#### 11.10. STARE (gABC model)^3^

We run the STARE (v1.0.4) using the command “STARE_ABCpp -b Enhancers.ATAC.tagAlign.sort.gz.CountReads.bedgraph -n 4 -a genes.gtf -w 10000000 -f false -t 0 -o [output_dir]” for each cell type. The file Enhancers.ATAC.tagAlign.sort.gz.CountReads.bedgraph is generated in the scE2G pipeline, located at [results_dir]/[cell_type]/[Neighborhoods]/Enhancers.ATAC.tagAlign.sort.gz.CountReads.bedgraph. The first three columns of this file represent the coordinates of the enhancer (chr, start, end), and the fourth column contains the read count for the corresponding enhancer.

#### 11.11. Making cell-type specific predictions

For benchmarks of published predictors required predictions for each cell type, we obtained candidate enhancer-gene pairs for a specific cell type by filtering enhancer-gene predictions made on the full cell population to a specific cluster based on which peaks were accessible in that cell type and which genes were expressed in that cell type. For example, to make predictions for the BMMC erythroblast cell type, we filtered predictions made on the combined BMMC dataset to those which contained peaks accessible in erythroblasts and genes expressed in erythroblasts. For Cicero, we only filtered predictions based on peak accessibility since the model does not use gene expression data. We defined genes detected (UMI count > 0) in more than 1% cells for each cell type as “expressed genes,” and defined MACS2 peaks that overlap with candidate enhancers identified in the ABC pipeline (see **Methods 9.1.1**) for each cell type as “accessible peaks.”

### 12. CRISPR benchmark

We used the CRISPR benchmarking pipeline developed in previous work^18^ to compare predictive models to perturbation data from previous studies in which CRISPR was used to perturb candidate enhancers and effects on gene expression were measured in K562 cells. The CRISPRi dataset is comprised of enhancer screens from 3 previously published CRISPRi enhancer screen datasets: 1) Nasser *et al.*, 2021^32^ (itself comprised most of CRISPRi screens from Fulco et al., 2019^17^), 2) Gasperini *et al.*, 2019^33^ (Perturb-seq) and 3) Schraivogel *et al.*, 2020^34^ (TAP-seq). The CRISPR benchmarking pipeline is implemented here: https://github.com/EngreitzLab/CRISPR_comparison

### 13. eQTL benchmark

We adapted our previously-developed eQTL benchmarking pipeline to assess whether predictive models of enhancer-gene regulatory interactions could (i) identify enhancers enriched for fine-mapped eQTL variants and (ii) link those enhancers to their corresponding eQTL target genes. We analyzed variants from the eQTL Catalogue V7 fine-mapped using the Sum of Single Effects (SuSiE) model^35^. The pipeline is implemented here: https://github.com/EngreitzLab/eQTLEnrichment/blob/integrated. Full results from the benchmarking analyses presented can be found here: https://github.com/anderssonlab/scE2G_analysis/blob/v1.0/3.Benchmarking/eQTL/

#### 13.1. Fine-mapped eQTL variants

We obtained the set of eQTL variants from the eQL Catalogue V7 fine-mapped with SuSiE from untreated biosamples with RNA sequencing quantified by “gene expression” that were filtered to variants to those with eGenes expressed in their respective biosamples^35^. We retained variants to those with a target gene included in the gene reference file described in **Methods 2**. For all analyses, we considered variants with a posterior inclusion probability greater than 50%. We grouped variants based on their biosample of origin at “fine-grained” and “coarse-grained” resolutions and retained biosamples with at least 35 variants. The code and table describing the collation of eQTL variants is available here: https://github.com/anderssonlab/scE2G_analysis/blob/v1.0/3.Benchmarking/eQTL/eQTL_Catalogue_v7

#### 13.2. Filtering to distal noncoding variants

For eQTL benchmarking analyses, we focused on distal noncoding variants by filtering out any variants overlapping (i) coding sequences, (ii) 5′ and 3′ untranslated regions of protein-coding genes, (iii) splice sites (within 10 bp of an intron–exon junction of a protein-coding gene) of protein-coding genes, and (iv) promoters (±250 bp from the gene TSS) of protein-coding genes. The disjoint genome partition used for this categorization is available here: https://github.com/EngreitzLab/eQTLEnrichment/blob/integrated/resources/genome_annotation/PartitionCombined.bed

#### 13.3. Defining prediction–eQTL cell type matches for eQTL benchmarking

We assigned each cell type with predicted regulatory enhancer-gene interactions to one or more closely-related eQTL biosamples (**Table S4**). In the “fine-grained” eQTL analysis (**Fig. 2d**,**e**, **Fig. S7a,b,g**) the following number of matching cell types was used for each method: 67 pairs for scE2G*^Multiome^*, scE2G*^ATAC^*, ABC, in element (ABC) & distance to TSS; 64 pairs for SnapATAC, Signac, Cicero, FigR, SCENIC+, DIRECT-NET, and ArchR; 26 pairs for STARE and SCENT. For the “coarse-grained” eQTL analysis (**Fig. S7c-f,h**), 3 pairs of matching cell types were used for ENCODE-rE2G, and 13 pairs were used for all other models. To calculate aggregate metrics across all pairs, we summed the variant counts for each matched set of cell types.

#### 13.4. Calculating enrichment of eQTLs in predicted enhancers

We defined enrichment as (fraction of distal noncoding eQTL variants with PIP > 50% that overlap predicted enhancers) / (fraction of distal noncoding 1000 Genomes Project SNPs that overlap predicted enhancers) (see **Fig. 2d**). To calculate this enrichment, we obtained a list of approximately 10 million SNPs from the 1000 Genomes Project from the Price Group ^36^ then filtered them to the distal noncoding regions of the genome as described in **Methods 13.2**. We calculated the number of variants from each eQTL biosample. Next, we calculated the number of eQTL variants from each biosample overlapping enhancer predictions defined by the suggested threshold in each biosample and the number of distal noncoding 1000 Genomes Project variants intersecting thresholded predictions in each biosample. The enrichment for each prediction cell type, P and eQTL cell type, Q intersection was then calculated as [number of distal noncoding variants in Q with PIP > 50% overlapping predicted enhancers in P / total number of distal noncoding variants with PIP > 50% in Q]/[number of distal noncoding 1000 Genomes Project SNPs overlapping P / total number of distal noncoding 1000 Genomes Project SNPs]. We calculated confidence intervals for enrichment using the analytical formula for confidence interval of the risk ratio^37^.

#### 13.5. Calculating recall of eQTLs in predicted enhancers

To calculate the recall of a predictive model in linking an eQTL variant to its eGene, we considered each pair of matching prediction and eQTL cell types (**Table S4**). For each pair, we calculated the 1) total recall, defined as the fraction of eQTL variants overlapping predicted enhancers; 2) recall accounting for gene-linking, defined as the fraction of eVariant-eGene pairs in which the eVariant overlaps a predicted enhancer linked to the correct eGene; and 3) for the set of variants that overlap any predicted enhancer, the fraction of eVariant-eGene pairs in which the overlapping enhancer is linked to the variant’s eGene. We calculated the 95% confidence interval of recall using the Wilson score interval implemented with the “plus four rule”^38^.

#### 13.6. Calculating enrichment-recall curves

We calculated a range of predictor score values to construct enrichment-recall curves (**Fig. 2d)** by defining 50 to 100 quantiles across the combined recall (considering gene-linking) of matching cell types for each predictive model with a continuous score. For each matching pair of prediction and eQTL cell types, we found the list of eVariants intersecting any predicted enhancer linked to the correct eGene, without applying a threshold to predictions. We compiled the variant-prediction intersections across all pairs of cell types. Then, we found the model score at the designated number of quantiles for predictions in this list, which estimate model scores required to achieve evenly-spaced recalls between 0 and the maximum recall for that model. These threshold values were used to binarize predictions to evaluate enrichment and recall as described above to generate enrichment-recall curves.

#### 13.7. Calculating enrichment at a specific recall

To compare the enrichment of eQTLs in predicted enhancers at a target recall (as in **Fig. 2e**), we first identified which threshold value out of all that were evaluated yielded a recall closest to the target value. If the recall at this threshold was within 10% of the target recall, the model is included in the comparison. The significance in the difference in enrichment between each pair of models was computed as using a two-side test on the difference in log relative risk^39^ and p-values were adjusted with the Bonferroni correction to account for multiple comparisons.

### 14. GWAS benchmark

Using an approach taken in previous work,^18,32^ we developed a GWAS benchmarking pipeline to evaluate predictive models based on (i) enrichment of fine-mapped GWAS variants and (ii) ability to link GWAS signals to causal genes (**Fig. 2f-h**). For this benchmark, we used GWAS data for 94 traits from the UK Biobank. The pipeline is implemented here: https://github.com/EngreitzLab/GWAS_E2G_benchmarking/tree/main

#### 14.1. Fine-mapped GWAS variants

We obtained GWAS summary statistics and fine-mapping results for 94 traits from the UK Biobank^40^. Detailed methods for the genetic association analysis, fine-mapping with SuSiE, and post-processing as well as the fine-mapping results are available at https://www.finucanelab.org/data. Definitions of each trait used in this study are summarized here: https://github.com/anderssonlab/scE2G_analysis/blob/v1.0/3.Benchmarking/GWAS/UKBB_94_traits.tsv. For all analyses, we retained variants with a fine-mapping posterior inclusion probability greater than 10% and that were located in distal noncoding regions of the genome as in **Methods 13.2**.

#### 14.2. Evaluation set of causal genes for GWAS signals

To compare how well predictive models link GWAS signals to causal genes, we used a “silver-standard” evaluation set developed in previous work using the fine-mapping results described above ^41^. Briefly, probable causal genes for each trait were defined as those containing a protein-coding, fine-mapped variant with a posterior inclusion probability greater than 50%. Then, each causal gene was linked to all independent noncoding credible sets for the same trait located within 500 kb. We removed duplicate credible sets in which the same noncoding variants were linked to the same gene for multiple traits, to avoid duplicates in the benchmark. This evaluation set of locus-gene links is larger, available for more traits, and less biased towards well-studied genes than other alternatives. However, it is limited by the assumptions that genes with protein-coding variants are causal for the trait and that multiple GWAS signals in the same locus have the same causal gene.

#### 14.3. Defining biosample-trait pairs for GWAS benchmarking

For the benchmarking evaluation, we assigned sets of biosamples to which the predictive models were applied with sets of GWAS traits expected to be relevant to those biosamples (**Table S5**). For example, predictions in the five BMMC and PBMC clusters for CD14+ monocytes and CD16+ monocytes were evaluated against the monocyte count trait. For the variant overlap analysis, the variants for a set of traits paired with a set of prediction biosamples were deduplicated before evaluation. In both analyses, a variant was considered to overlap an enhancer if it intersected a predicted enhancer in at least one biosample at a given score threshold. In addition, a model was only evaluated for a given biosample-set, trait-set pair if it had been applied to all biosamples in the set.

#### 14.4. Calculating enrichment and recall of variants in predicted enhancers

We calculated the recall of variants from trait (or set of traits) T in enhancer predictions from biosample (or set of biosamples) P as: [number of distal noncoding GWAS variants in T with PIP > 10% overlapping predicted enhancers in P / total number of distal noncoding variants with PIP > 10% in T]. We calculated the enrichment of variants from trait (or set of traits) T in enhancer predictions from biosample (or set of biosamples) P as: [number of distal noncoding GWAS variants in T with PIP > 10% overlapping predicted enhancers in P / total number of distal noncoding variants with PIP > 10% in T]/[number of distal noncoding 1000 Genomes Project SNPs overlapping P / total number of distal noncoding 1000 Genomes Project SNPs]. The 1000 Genomes Project SNPs were curated as described in **Methods 13.4**. We calculated 95% confidence intervals for recall using the Wilson score interval implemented with the “plus four rule”^38^ and for enrichment using the analytical formula for confidence interval of the risk ratio^37^.

#### 14.5. Calculating enrichment-recall curves

For each continuous predictive model, we calculated a range of model scores at which to threshold predictions. To do so, we first defined 25 thresholds evenly-spaced across the range of possible scores. Next, we found the predicted enhancer with the highest score in any biosample overlapping variants from all 94 traits, and calculated the scores at 25 quantiles for the predictions in this list to estimate thresholds required to achieve evenly-spaced recalls between 0 and the maximum recall for that model. These threshold values were used to binarize predictions to evaluate enrichment and recall as described above to generate enrichment-recall curves.

#### 14.6. Calculating precision and recall of identifying causal genes for GWAS signals

We calculated the precision and recall of model predictions in each biosample and biosample set at identifying the causal genes associated with noncoding credible sets from each trait described in **Methods 14.2**. After intersecting variants with enhancer-gene predictions binarized at the suggested thresholds, we defined credible set-gene predictions as each credible set linked to the two genes linked to enhancers overlapping variants in the credible set with the highest scores (or, for binary predictors, all genes linked to enhancers overlapping variants in the credible set). Recall was calculated as (number of correct credible set-gene predictions / total number of credible sets). Precision was calculated as (number of correct credible set-gene predictions / total number of credible sets-gene predictions). 95% confidence intervals for both metrics were calculated using the Wilson score interval implemented with the “plus four rule”^38^.

#### 14.7. Combining model predictions with PoPS

To calculate precision and recall at identifying causal genes for each model in combination with the polygenic priority score (PoPS), we used the PoPS scores given to genes for each trait^41^ to find the top two genes ranked by PoPS within 1 Mb of each noncoding credible set in the evaluation set. We then defined credible set-gene predictions for a model ∩ PoPS as each credible set linked to any gene that ranked in the top two based on both model score as in **Methods 14.6** and PoPS score. Recall and precision were calculated as described above with this new definition of credible set-gene predictions.

### 15. Analyzing cell-type specificity of scE2G predictions

The code for all analyses of the cell-type specificity of scE2G predictions is available here: https://github.com/anderssonlab/scE2G_analysis/blob/v1.0/6.Cell_Type_Specificity

#### 15.1. Computing Jaccard similarity between pairs of scE2G predictions

To compare the fraction of shared enhancers between scE2G predictions in each pair of cell types, we quantified overlap of predicted enhancer-gene links using the Jaccard similarity. We considered thresholded predictions (scE2G score greater than 0.164 for scE2G*^Multiome^*, no promoter elements) for each cell type. We computed the Jaccard similarity between cell type A and cell type B as: (number of shared predictions) / (number of total predictions for cell type A + number of total predictions for cell type B - number of shared predictions). We defined a “shared prediction” as enhancer-gene links with the same gene and at least a 50% overlap of each enhancer width.

#### 15.2. Computing Pearson correlation between pairs of scE2G predictions

To compare the correlation of scE2G scores between predictions in each pair of cell types, we calculated the Pearson correlation between prediction scores in each pair of cell types. We considered only thresholded predictions and defined “shared predictions” as described above. For enhancer-gene predictions that were not shared between the two cell types, we set the score in the cell type without that prediction to 0. We then computed the Pearson correlation between the matched scores for each pair of cell types.

#### 15.3. Analyzing cell-type specificity of eQTL variant enrichment and recall

We categorized each eQTL cell type, prediction cell type pair as an “exact match” (same cell type, such as eQTLs from CD4+ T cells with predictions in any T cell type), “similar” (same cell lineage, such as a eQTLs from CD4+ T cells with predictions in any lymphoid cell type), or “unmatched” (all other pairs). The categorizations used in **Fig. 5b** and **c** can be found here: https://github.com/anderssonlab/scE2G_analysis/blob/v1.0/6.Cell_Type_Specificity/eQTL_enrichment_recall/all_pred_eQTL_key.tsv.

#### 15.4. Computing number of predicted enhancer-gene links at different cell type resolutions

To determine whether scE2G makes more enhancer-gene links for more fine-grained cell types, we compared the number of scE2G*^Multiome^* predictions made for a cell type supergroup to the number of predictions made across cell types within a supergroup from the BMMC dataset. To control for a change in number of predictions from analyzing more or smaller cell clusters, we ran scE2G on samples of similar cell numbers, selected from either each cell type or from supergroup, and aimed for each sample of cells to contain over 2 million ATAC fragments and close to 1 million RNA UMIs. For example, we divided the three B cell types (B1 B cells, CD20+ naive B cells, and transitional B cells) into 3, 10, and 5 samples of cells with 500–600 cells, 5–8 million ATAC fragments, and 0.75–1.25 million RNA UMIs per sample. We then divided the cells from the entire B cell supergroup into 19 samples of cells of similar size. The sample sizes for these two sets of cell samples can be found https://github.com/anderssonlab/scE2G_analysis/blob/v1.0/6.Cell_Type_Specificity/number_of_predictions/cell_samples_from_cell_types.tsv and https://github.com/anderssonlab/scE2G_analysis/blob/v1.0/6.Cell_Type_Specificity/number_of_predictions/cell_samples_from_supergroups.tsv. We then ran scE2G*^Multiome^* on all of these samples.

We moved forward with the four BMMC supergroups with sufficient samples to compare at least 100 unique combinations of samples per cell type (B, T, Myeloid, Erythroid). For each supergroups, we chose at random one sample that was drawn from each cell type, and the same number of samples from the supergroup-level samples (*e.g.*, one sample each from B1 B cells, CD20+ naive B cells, and transitional B cells, and three samples from the B cell supergroup). We counted the number of unique enhancer-gene predictions across the cell-type-level samples and the number of predictions across the supergroup-level samples, using the same definition of “shared predictions” described in **Methods 15.1**. We repeated this selection and counting process 100 times per supergroup. To summarize the results, we ensured that each replicate comprised a unique set of samples, then counted the number of replicates in which the cell-type-level samples yielded more enhancer-gene links than the supergroup-level samples.

### 16. Applying scE2G to prioritize causal genes and cell types for GWAS loci

#### 16.1. Computing enrichment of heritability of GWAS traits with stratified linkage disequilibrium score regression

To estimate the enrichment of trait heritability in predicted enhancers of each cell type, we used stratified linkage disequilibrium score regression (S-LDSC, v1.0.1^42,43^). We considered all distal enhancers (no promoters) predicted by scE2G*^Multiome^* for each cell type. We prepared summary statistics for 104 traits^40^ (GWAS summary statistics from the UK Biobank obtained from https://www.finucanelab.org/data), using the `munge_sumstats.py` script provided by LDSC and filtered to retain SNPs curated by HapMap 3 as recommended^44^. We performed S-LDSC using the scE2GMultiome enhancers along with the baseline model (v2.2). We used the 1000 Genome EUR Phase 3 genotype data as the LD reference panel. Full results from S-LDSC can be found in **Table S8**.

#### 16.2. Prioritizing causal genes and cell types for GWAS signals

To systematically apply scE2G to prioritize causal genes and cell types for GWAS signals, we first determined which traits had variants that were 1) significantly enriched and 2) enriched at least 5-fold in predicted enhancer from which cell types. We considered blood-related traits for the 31 cell types from BMMCs and PBMCs, and metabolic-related traits for the 9 cell types from pancreatic islets. We found all credible set, cell type, gene combinations for which the credible set has noncoding variants located in a predicted scE2G*^Multiome^* enhancer linked to that gene in that cell type. We recorded the maximum scE2G score supporting that prediction, as well as all other cell types in which the credible set was linked to that gene. We also annotated each credible set with the nearest genes based on distance. We then filtered the predictions to the two genes linked to each credible set with the highest maximum scE2G scores (“scE2G rank” of 1 and 2). The full gene prioritization results (not filtered based on scE2G rank) can be found in **Table S9**.

#### 16.3. Prioritizing causal genes for GWAS traits with the polygenic priority score

We used Polygenic Priority Scores (PoPS) as an orthogonal gene prioritization method to scE2G^41^. We used PoPS results obtained in our previous work^32^.

#### 16.4. Quantifying variant effect on transcription factor motifs

We quantified the effect of a variant on a given transcription factor motif using the match score computed with FIMO^45^. We obtained position-weight matrices for the transcription factor motif of interest from JASPAR^46,47^, and input 101 base pair sequences centered on the reference and alternate alleles. The match scores reported correspond to the same window from each allele.

### 17. Data visualization

#### 17.1. Box plots

Box plots are defined as follows: the middle line or point corresponds to the median; lower and upper hinges correspond to first and third quartiles; the upper whisker extends from the hinge to the largest value no further than 1.5 × IQR from the hinge (where IQR is the interquartile range, or distance between the first and third quartiles); and the lower whisker extends from the hinge to the smallest value, at most 1.5 × IQR of the hinge. Data beyond the end of the whiskers are outlying points and are plotted individually, unless box plots are shown on top of all data points or data distribution.

#### 17.2. Point intervals

Histograms and violin plots containing point interval plots were made with the ggdist package^48^. The middle point corresponds to the median; the thick line corresponds to the quantile interval of the center 66% of data; the thin line corresponds to the quantile interval of center 95% of data.

#### 17.3. Locus plots

All locus plots were made with the GViz^49^ and GenomicInteractions^50^ packages. Pseudobulk ATAC-seq signal tracks are plotted based on the normalized bigwig file generated in the scE2G pipeline, in which the ATAC signal is scaled by the number of total ATAC fragments (see `frag_to_norm_bigWig` rule here: https://github.com/EngreitzLab/scE2G/blob/v1.0/workflow/rules/frag_to_tagAlign.smk).

